# Manganese transporter SLC30A10 and iron transporters SLC40A1 and SLC11A2 impact dietary manganese absorption

**DOI:** 10.1101/2024.07.17.603814

**Authors:** Milankumar Prajapati, Jared Z. Zhang, Grace S. Chong, Lauren Chiu, Courtney J. Mercadante, Heather L. Kowalski, Olga Antipova, Barry Lai, Martina Ralle, Brian P. Jackson, Tracy Punshon, Shuling Guo, Mariam Aghajan, Thomas B. Bartnikas

**Affiliations:** Department of Pathology and Laboratory Medicine, Brown University, Providence, RI, 02912, USA; Advanced Photon Source, Argonne National Laboratory, Argonne, IL, 60439, USA; Department of Molecular and Medical Genetics, Oregon Health & Science University, Portland, OR, 97239, USA; Biomedical National Elemental Imaging Resource, Dartmouth College, Hanover, NH, 03755, USA; Ionis Pharmaceuticals, Inc., Carlsbad, CA, 92010, USA

## Abstract

SLC30A10 deficiency is a disease of severe manganese excess attributed to loss of SLC30A10-dependent manganese excretion via the gastrointestinal tract. Patients develop dystonia, cirrhosis, and polycythemia. They are treated with chelators but also respond to oral iron, suggesting that iron can outcompete manganese for absorption in this disease. Here we explore the latter observation. Intriguingly, manganese absorption is increased in Slc30a10-deficient mice despite manganese excess. Studies of multiple mouse models indicate that increased dietary manganese absorption reflects two processes: loss of manganese export from enterocytes into the gastrointestinal tract lumen by SLC30A10, and increased absorption of dietary manganese by iron transporters SLC11A2 (DMT1) and SLC40A1 (ferroportin). Our work demonstrates that aberrant absorption contributes prominently to SLC30A10 deficiency and expands our understanding of biological interactions between iron and manganese. Based on these results, we propose a reconsideration of the role of iron transporters in manganese homeostasis is warranted.

---

Manganese (Mn) is an essential metal and dietary nutrient^1,2^. It plays key roles in biological processes including antioxidant defense and protein synthesis but in excess causes oxidative stress and mitochondrial dysfunction. Given the essential yet potentially toxic nature of this metal, body Mn levels are carefully regulated, largely by hepatobiliary excretion. Our understanding of the molecular basis of Mn homeostasis has benefited greatly from studies of recently identified inherited diseases. One such disease is SLC30A10 deficiency, characterized by severe Mn excess, neurologic dysfunction, liver failure, polycythemia, and erythropoietin excess^3,4^. SLC30A10 is an apical membrane protein that exports Mn from hepatocytes into bile and from enterocytes into the gastrointestinal tract lumen^5–7^. SLC30A10 deficiency leads to impaired gastrointestinal Mn excretion, resulting in Mn excess. Neurologic and liver dysfunction are attributed to Mn toxicity. Polycythemia reflects excessive synthesis of erythropoietin by the liver due to Mn-induced upregulation of hepatic hypoxia-inducible factor 2 (Hif2) signaling^8^.

Current treatments for SLC30A10 deficiency include chelation and oral iron (Fe) supplementation. The efficacy of the latter is attributed to Fe outcompeting Mn for dietary absorption, which suggests that Fe and Mn share absorption pathways in this disease. Like Mn, Fe is an essential dietary nutrient^2,9^. This metal is essential for multiple biological processes including oxygen transport, energy metabolism, and DNA synthesis. In excess, it is toxic due to its propensity to catalyze the formation of reactive oxygen species. In contrast to Mn, body Fe levels are determined mainly by absorption^9^. SLC11A2 (DMT1) imports non-heme Fe from the gastrointestinal tract lumen into enterocytes. SLC40A1 (ferroportin, FPN) exports Fe from enterocytes into blood. Hepcidin, a hormone synthesized largely by the liver, post-translationally downregulates FPN. Hepcidin suppresses Fe absorption under conditions of Fe excess. Hepcidin deficiency leads to increased Fe absorption under conditions of Fe deficiency. Perturbations in hepcidin expression can have severe impacts on Fe homeostasis, as seen in hereditary hemochromatosis, a disease of Fe excess caused by mutations in hepcidin or genes essential for hepcidin expression^10^. Hepcidin deficiency is also observed in SLC30A10 deficiency. In contrast to hereditary hemochromatosis, hepcidin deficiency in SLC30A10 deficiency reflects indirect suppression of liver hepcidin expression in the setting of erythropoietin excess to ensure adequate Fe supply for erythropoiesis.

The efficacy of oral Fe as a treatment for SLC30A10 deficiency suggests that DMT1 and FPN, both essential for Fe absorption, mediate Mn absorption in SLC30A10 deficiency. The molecular basis of Mn absorption is unknown, although studies have implicated FPN and DMT1 in Mn transport. Several cell culture-based or structural studies have reported that DMT1 can transport Mn^11–16^. The Belgrade rat, harboring a missense DMT1 mutation, has impaired olfactory ^54^Mn uptake and intestinal Mn transport^17,18^. However, Mn absorption is intact in mice with intestinal Dmt1 deficiency^19^. With regards to Mn transport by FPN, increased absorption of gavaged ^54^Mn was noted in a mouse model of hereditary hemochromatosis deficient in Hfe, a protein required for hepcidin expression^20^. Two other studies supportive of a FPN-Mn link employed the flatiron mouse, but this line harbors a dominant Fpn missense mutation found in an atypical variant of hereditary hemochromatosis^21,22^. Other studies demonstrating Mn transport by Fpn employed functional assays in cells, not mammals^23–25^. Two studies reported no evidence of a prominent impact of Fpn on Mn homeostasis, although neither assessed Mn absorption or excretion. One study was based in *Xenopus* oocytes^26^. The other employed mice with intestinal Fpn deficiency and mice deficient in Tmprss6, a transmembrane protease that downregulates hepcidin expression^27^.

In this study, we hypothesize that one context in which DMT1 and FPN do transport Mn is SLC30A10 deficiency, given that patients respond to oral Fe. To test this hypothesis, we first assessed Mn absorption in *Slc30a10^-/-^*mice. Finding increased Mn absorption in mutant mice, we next established the impact of intestinal Dmt1 and Fpn deficiency on Mn absorption and excess in *Slc30a10^-/-^* mice. Finally, to explore a potential mechanism for increased Mn absorption in *Slc30a10^-/-^* mice, we interrogated the role of intestinal Slc30a10 in modulating Mn absorption, given that Slc30a10 exports Mn from enterocytes into the gastrointestinal tract lumen. Results and their implications are presented below.

## Results

### Slc30a10^-/-^ mice have increased Mn absorption

To interrogate the role of intestinal Fe transporters in SLC30A10 deficiency, we first assessed dietary Mn absorption. Intriguingly, two-month-old *Slc30a10^-/-^* mice absorbed more gavaged ^54^Mn than did *Slc30a10^+/+^* mice (Fig. 1a). Increased ^54^Mn absorption and liver Mn excess first occurred on post-natal day (P) 21, while increased red blood cell (RBC) counts were noted first at P28 (Fig. 1b-d). To determine if aberrant Mn absorption solely reflected Mn excess, wild-type mice were weaned onto diets with varied Mn content. This produced expected changes in liver Mn content but did not impact ^54^Mn absorption (Fig. 1e, f). Given the aberrant Mn absorption in mutant mice, we next interrogated Mn levels in adult *Slc30a10^-/-^* duodenum by X-ray fluorescence microscopy. This identified punctate Mn excess in enterocytes and prominent Mn signal throughout goblet cells, consistent with perturbed intestinal Mn homeostasis (Fig. 1g, S1). As mentioned above, Dmt1 and Fpn are both essential for dietary Fe absorption yet have also been implicated in Mn transport by some studies. We observed increased apical Dmt1 and basolateral Fpn levels in *Slc30a10^-/-^* enterocytes (Fig. 1h, S2-9). This was not unexpected given that *Slc30a10^-/-^* mice have increased dietary Fe absorption^8,28^. However, the relevance of intestinal Dmt1 and Fpn to Mn absorption in mutant mice remained to be determined.

**Fig. 1:**
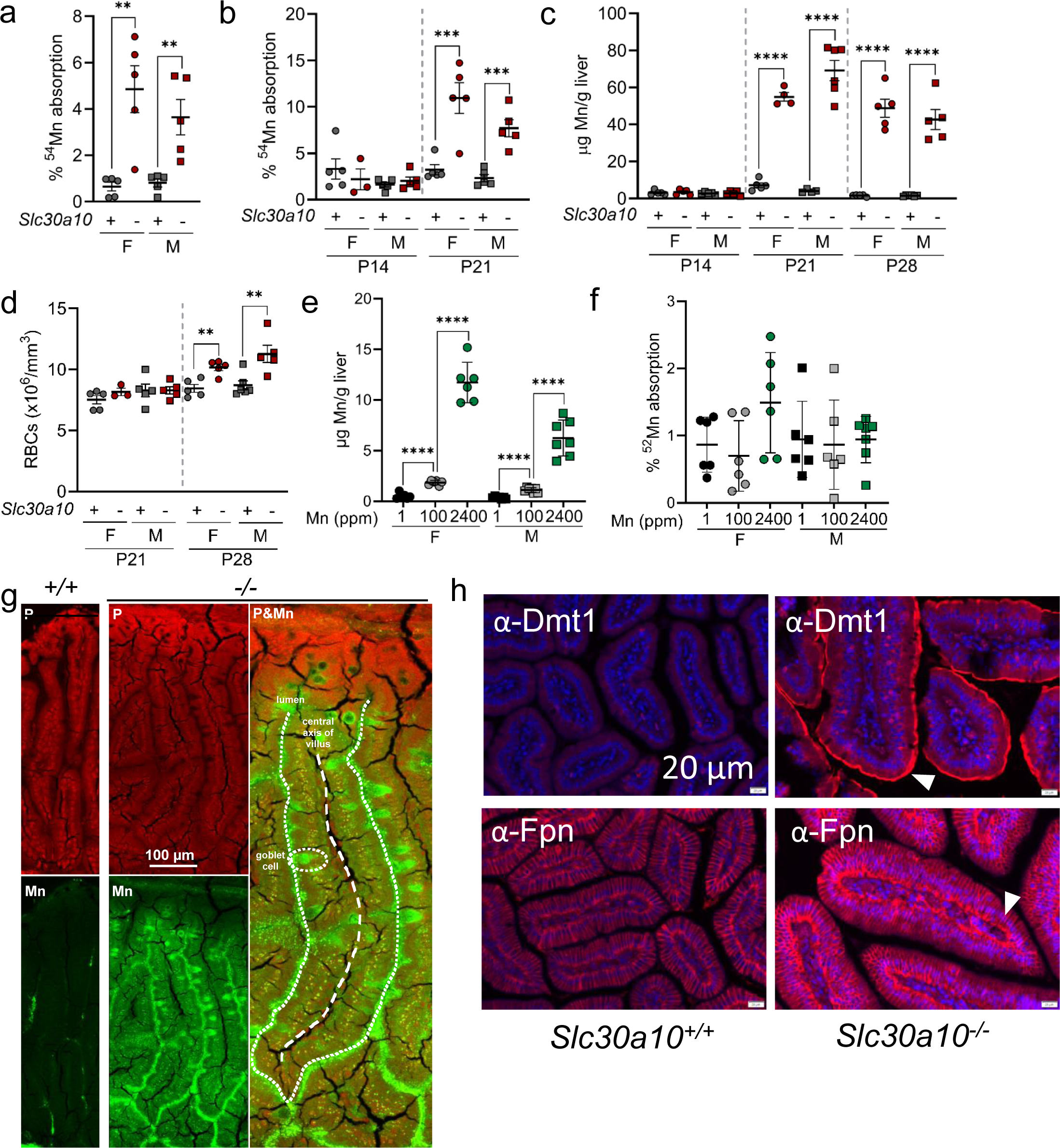
*Slc30a10^-/-^*mice have increased Mn absorption. (**a**) Two-month-old female (F) and male (M) *Slc30a10^+/+^* (‘+’) and *Slc30a10^-/-^* (‘-’) mice were analyzed for Mn absorption by ^54^Mn gavage. (**b-d**) P14-28 *Slc30a10^+/+^* and *Slc30a10^-/-^* mice were analyzed for Mn absorption (b), liver Mn levels by ICP-OES (c), and RBC counts by complete blood counts (d). (**e, f**) Wild-type mice weaned onto diets containing 1, 100, or 2400 ppm Mn were analyzed at two months of age for liver Mn levels (e) and ^52^Mn absorption (f). (**g**) X-ray fluorescence microscopy of small intestinal sections from two-month-old *Slc30a10^+/+^*and *Slc30a10^-/-^* mice. (**h**) Immunofluorescence for Dmt1 (h) and Fpn (i) in small intestinal sections from two-month-old *Slc30a10^+/+^*and *Slc30a10^-/-^* mice. Apical Dmt1 and basolateral Fpn staining are indicated by arrowheads. For (a-f), data are represented as means +/- standard deviation, with at least three animals per group. Data were tested for normal distribution by Shapiro-Wilk test; if not normally distributed, data were log transformed. Within each sex, groups were compared using unpaired, two-tailed tests (a-d) or one-way (e, f) ANOVA with Tukey’s multiple comparisons test. (* P<0.05, ** P<0.01, *** P<0.001, **** P<0.0001)

### Intestinal Dmt1 deficiency attenuates Mn absorption and excess in Slc30a10^-/-^ mice

To explore the impact of intestinal Fe transporters on Mn absorption and excess in *Slc30a10^-/-^* mice, we first focused on Dmt1. Intestinal Dmt1 deficiency was induced prior to the onset of Mn excess in two-week-old mice carrying floxed *Dmt1* alleles (*Dmt1^fl/fl^*) and a tamoxifen-inducible *Vil* Cre transgene (*Vil-ERT2*). After tamoxifen treatment, a set of *Vil-ERT2* mice were injected with Fe dextran to correct Fe deficiency secondary to intestinal Dmt1 deficiency. At four weeks of age, *Slc30a10* RNA levels were decreased in *Slc30a10^-/-^* duodenum and liver, while *Dmt1* RNA levels were decreased in duodenum but not liver of *Vil-ERT2* mice, as expected (Fig. S10). Notably, *Slc30a10* RNA levels were increased in *Slc30a10^+/+^ Dmt1^fl/fl^ Vil-ERT2* mice not treated with Fe dextran. This may reflect hypoxia-inducible factor (Hif)-dependent Slc30a10 upregulation in the setting of Fe deficiency, as Slc30a10 is regulated by Hifs^29^ and Fe deficiency increases Hif levels^9^. Dmt1 protein levels were decreased in intestinal epithelia of *Slc30a10^-/-^ Dmt1^fl/fl^ Vil-ERT2* mice (Fig. S11-14).

Consistent with Dmt1’s role in Fe absorption, *Vil-ERT2* mice had decreased liver non-heme Fe levels, liver hepcidin RNA levels, and RBC parameters but unaltered liver copper and zinc levels (Fig. S15). Fe dextran treatment of *Vil-ERT2* mice increased liver non-heme Fe levels, liver *Hamp* RNA levels, and RBC parameters, although RBC parameters were not impacted in male *Slc30a10^-/-^* mice. Intestinal Dmt1 deficiency had no impact on Mn levels in *Slc30a10^+/+^* mice but decreased tissue Mn levels in *Slc30a10^-/-^* mice irrespective of Fe dextran treatment, suggesting that Fe deficiency was not contributing to the impact of intestinal Dmt1 deficiency on Mn excess (Fig. 2a-f). Intestinal Dmt1 deficiency had no impact on bile flow rates or Mn levels but did decrease ^52^Mn absorption in *Slc30a10^-/-^* mice (data not shown; Fig. 2g-i; absorption data for both sexes pooled in 2g, unpooled in 2i).

**Fig. 2:**
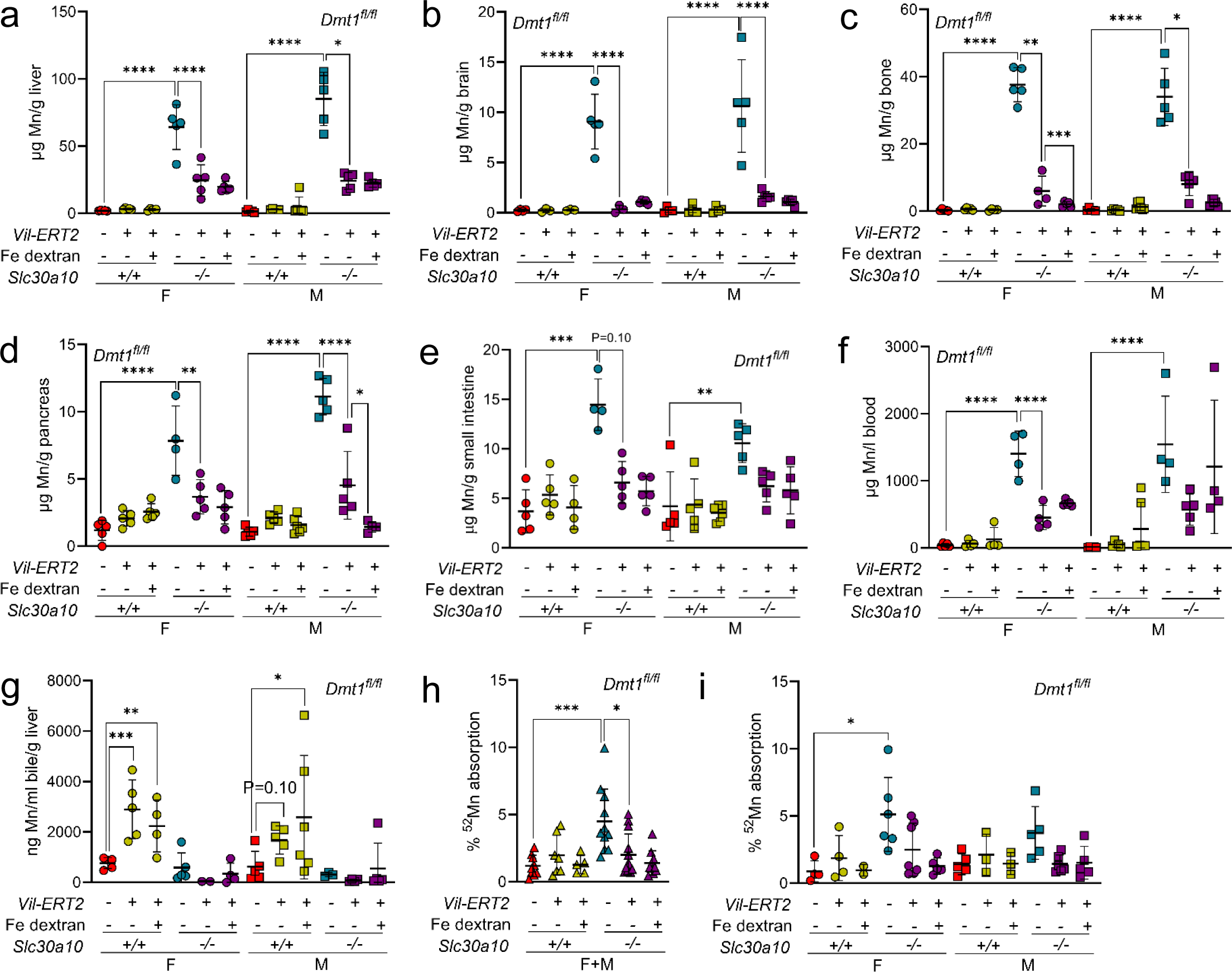
Intestinal Dmt1 deficiency, initiated at P14 before the onset of Mn excess, attenuates Mn excess in *Slc30a10^-/-^* mice. *Slc30a10 Dmt1^fl/fl^*mice with and without *Vil-ERT2* were injected intraperitoneally with a five-day course of tamoxifen starting at P14. Some mice were injected subcutaneously with Fe dextran on P19. At P28, mice were analyzed for: (**a-e**) liver (a), brain (b), bone (c), pancreas (d), and small intestine (e) Mn levels by ICP-OES; (**f**) blood Mn levels by GFAAS; (**g**) bile Mn levels by GFAAS; (**h, i**) Mn absorption by ^52^Mn gavage, with female and male data pooled in (h) and analyzed separately in (i). In all panels, data are represented as means +/- standard deviation, with at least four animals per group. Data were tested for normal distribution by Shapiro-Wilk test; if not normally distributed, data were log transformed. Within each sex, groups were compared using two-way ANOVA with Tukey’s multiple comparisons test. (* P<0.05, ** P<0.01, *** P<0.001, **** P<0.0001)

To examine the impact of intestinal Dmt1 deficiency on the brain, an organ prominently impacted in SLC30A10 deficiency, we performed bulk RNA-sequencing (RNA-seq). We previously demonstrated using this approach that ∼300 genes were differentially expressed in brains of two-month-old *Slc30a10^-/-^* mice^8,28^. Here we detected 76 genes impacted by Slc30a10 deficiency and 16 genes impacted by intestinal Dmt1 deficiency in female *Slc30a10^-/-^* mice (Fig. S16). Nine genes were shared between these groups, with intestinal Dmt1 deficiency normalizing their expression (Fig. S17). The modest number of differentially expressed genes may reflect the fact that mice were only four weeks old when analyzed.

To determine the impact of intestinal Dmt1 deficiency on *Slc30a10^-/-^*phenotypes after the onset of Mn excess, we treated mice with tamoxifen at three weeks of age then aged them to eight weeks. We pooled sexes for analysis, since several *Slc30a10^-/-^ Dmt1^fl/fl^ Vil-ERT2* mice did not survive to eight weeks. *Dmt1* RNA levels were decreased in duodenum but not liver for *Vil-ERT2* mice; *Slc30a10* RNA levels were decreased in *Slc30a10^-/-^* duodenum and liver (Fig. 3a-d). Intestinal Dmt1 deficiency decreased liver Fe levels (*Slc30a10^-/-^*only) and RBC parameters (RBC counts in *Slc30a10^-/-^* only) in *Slc30a10* mice (Fig. 3e-h). Intestinal Dmt1 deficiency also decreased tissue and blood Mn levels in *Slc30a10^-/-^* mice but not *Slc30a10^+/+^*mice (Fig. 3i-n).

**Fig. 3:**
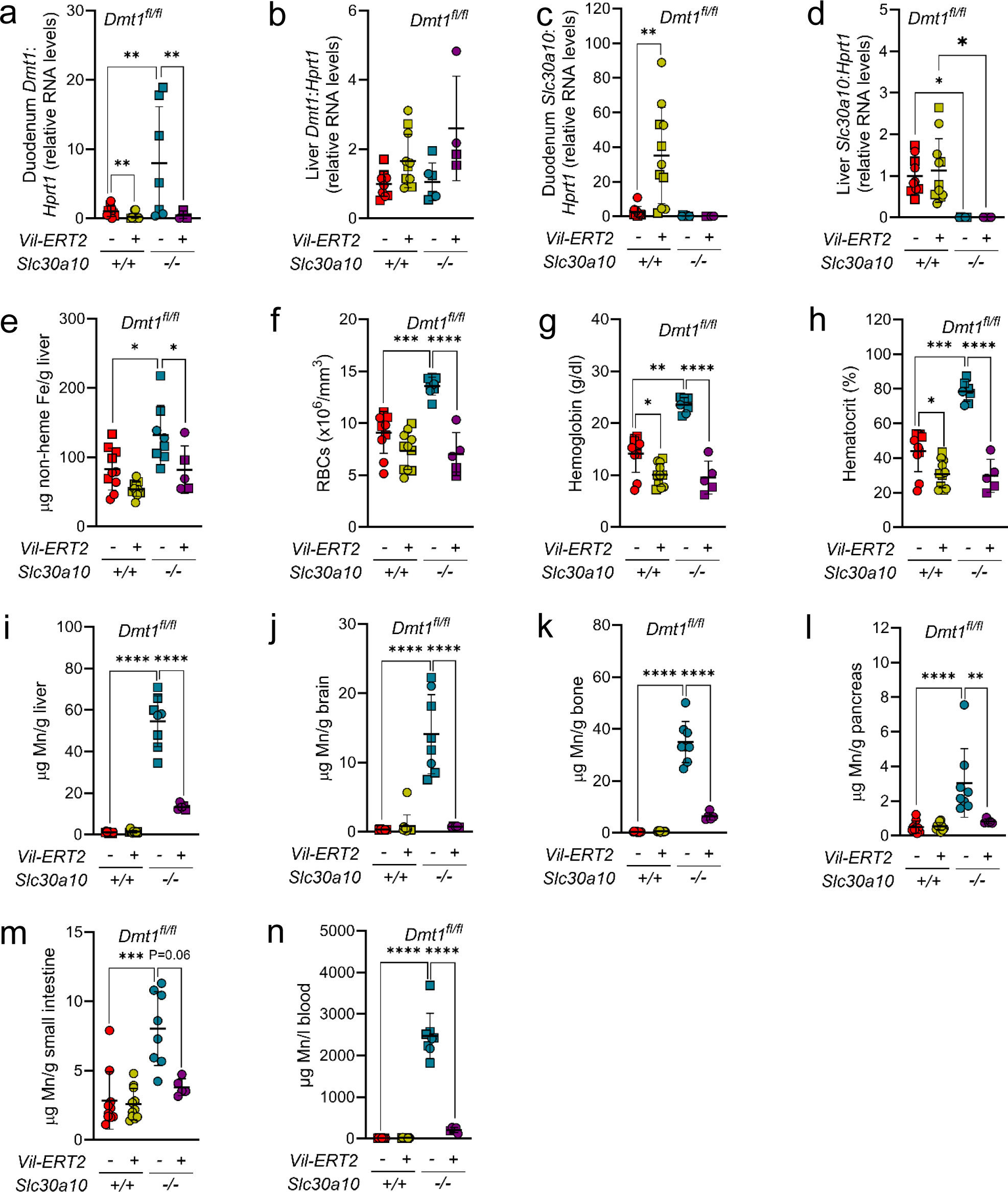
Intestinal Dmt1 deficiency, initiated at P21 after the onset of Mn excess, attenuates Mn excess in *Slc30a10^-/-^* mice. *Slc30a10 Dmt1^fl/fl^* mice with and without *Vil-ERT2* were injected intraperitoneally with a five-day course of tamoxifen starting at P21. At eight weeks of age, mice were analyzed for: (**a-d**) duodenum (a) and liver (b) *Dmt1* and duodenum (c) and liver (d) *Slc30a10* RNA levels by qPCR; (**e**) liver non-heme Fe levels by bathophenanthroline-based assay; (**f-h**) RBC counts (f), hemoglobin levels (g), and hematocrits (h) by complete blood counts; (**i-m**) liver (i), brain (j), bone (k), pancreas (l), and small intestine (m) Mn levels by ICP-OES; (**n**) blood Mn levels by GFAAS. Data are represented as means +/- standard deviation, with at least four animals per group. Data were tested for normal distribution by Shapiro-Wilk test; if not normally distributed, data were log transformed. Within each sex, groups were compared using two-way ANOVA with Tukey’s multiple comparisons test. (* P<0.05, ** P<0.01, *** P<0.001, **** P<0.0001)

While intestinal Dmt1 deficiency had a prominent impact on Mn levels in *Slc30a10^-/-^* mice, it had minimal impact on Mn levels in *Slc30a10^+/+^* mice, consistent with a previous report^19^. We considered that the minimal impact in *Slc30a10^+/+^* mice reflected compensatory changes in expression of other Mn transporters, particularly given that intestinal Dmt1 deficiency in *Slc30a10^+/+^*mice increased bile Mn levels (Fig. 2g). *Slc30a10^+/+^ Dmt1^fl/fl^ Vil-ERT2* mice had increased liver and duodenal *Slc30a10* RNA levels and liver *Slc39a14* and *Slc39a8* RNA levels (Fig. S10a, b, S18). SLC39A14 is essential for liver Mn import, which could contribute to increased bile Mn levels in these mice^30^. SLC39A8 is essential for import of bile Mn into hepatocytes, and mice with intestinal Slc39a8 deficiency do have decreased, but not ablated, absorption of gavaged ^54^Mn^30,31^. SLC39A8 can also transport Fe, so duodenal *Slc39a8* upregulation could represent a response to Fe deficiency^32–34^. Duodenum *Slc30a10* RNA and liver *Slc39a14* and *Slc39a8* RNA levels were not increased in mice treated with Fe dextran, suggesting that expression was driven by Fe deficiency.

### Intestinal Fpn deficiency attenuates Mn absorption and excess in Slc30a10^-/-^ mice

Having established intestinal Dmt1 as essential for Mn excess in *Slc30a10^-/-^*mice, we next explored the contribution of Fpn. We inactivated intestinal Fpn in *Slc30a10* mice carrying floxed *Fpn* alleles (*Fpn^fl/fl^*) and *Vil-ERT2* by treating them with tamoxifen at two weeks of age. Some *Vil-ERT2* mice were injected with Fe dextran. At four weeks of age, *Slc30a10* RNA levels were decreased in *Slc30a10^-/-^*duodenum and liver, while *Fpn* RNA levels were decreased in duodenum but not liver of *Vil-ERT2* (Fig. S19). Fpn protein levels were decreased in intestinal epithelia of all *Vil-ERT2* mice (Fig. S20-23). Consistent with Fpn’s role in Fe absorption, *Vil-ERT2* mice had decreased liver non-heme Fe levels, liver hepcidin RNA levels, and RBC parameters while liver copper and zinc levels were unaffected (Fig. S24). Fe dextran treatment increased liver non-heme Fe levels, liver hepcidin RNA levels (*Slc30a10^+/+^*only), and RBC parameters. Intestinal Fpn deficiency did not impact Mn levels in *Slc30a10^+/+^* mice but did decrease Mn levels in *Slc30a10^-/-^*mice irrespective of Fe dextran treatment (Fig. 4a-f). Intestinal Fpn deficiency did not impact bile flow rates nor did it increase bile Mn levels in *Slc30a10^-/-^* mice (data not shown; Fig. 4g). Intestinal Fpn deficiency decreased ^52^Mn absorption levels in *Slc30a10^-/-^*mice (Fig. 4h, i; female and male data pooled in h, unpooled in i). Bulk RNA-seq on brains detected 66 genes differentially expressed in female *Slc30a10^-/-^* mice and 77 genes differentially expressed in female *Slc30a10^-/-^* mice with intestinal Fpn deficiency (Fig. S25). Seventeen genes were shared between these groups of 66 and 77 genes, with intestinal Fpn deficiency normalizing their expression in mutant mice (Fig. S26). Finally, to determine the impact of intestinal Fpn inactivation after the onset of Mn excess in *Slc30a10^-/-^* mice, *Slc30a10 Fpn^fl/fl^* mice, with and without *Vil-ERT2*, were treated with tamoxifen at three weeks of age then aged to eight weeks. Most *Slc30a10^-/-^ Fpn^fl/fl^ Vil-ERT2* mice did not survive, precluding analysis (data not shown).

**Fig. 4:**
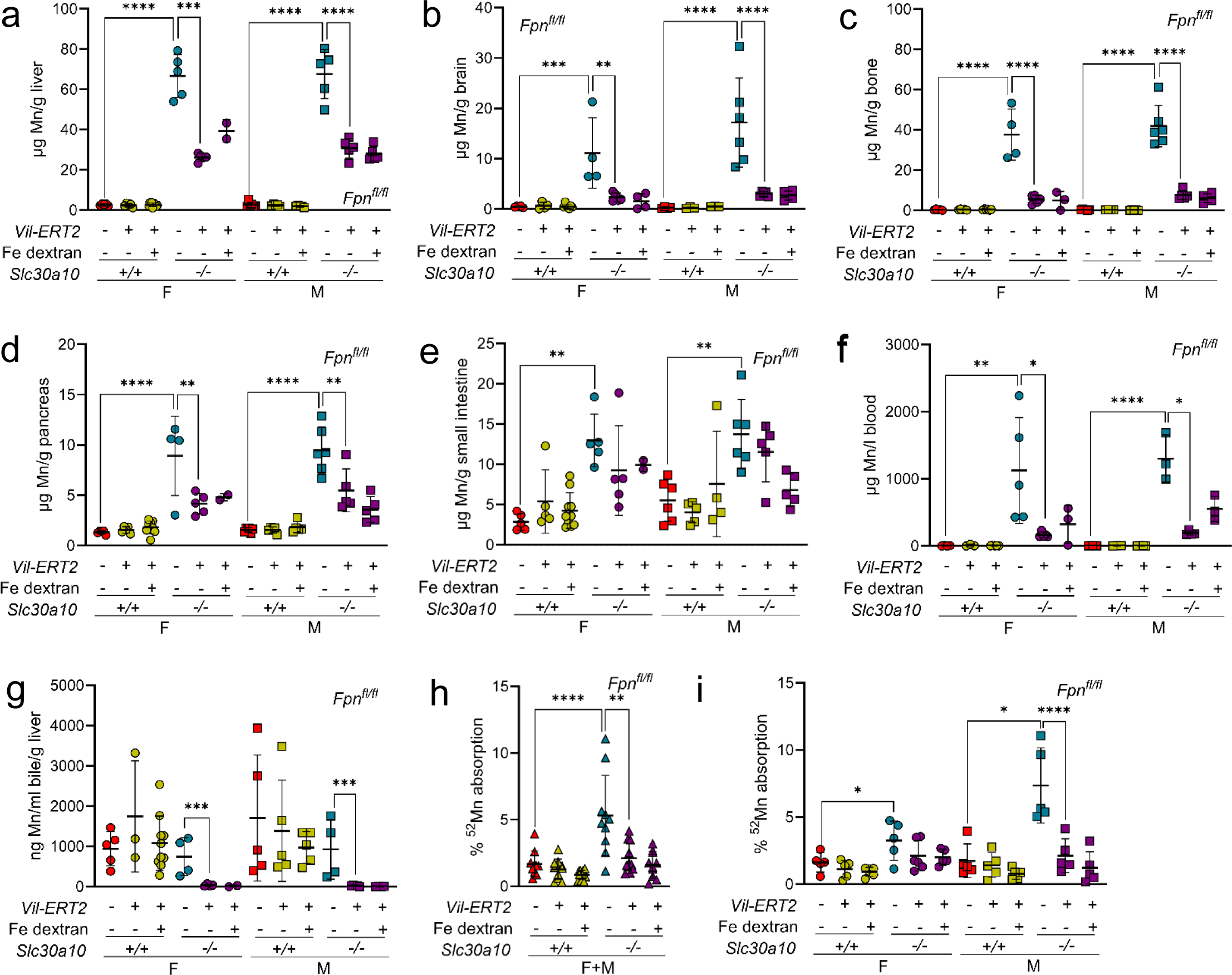
Intestinal Fpn deficiency, initiated at P14 before the onset of Mn excess, attenuates Mn excess in *Slc30a10^-/-^* mice. *Slc30a10 Fpn^fl/fl^* mice with and without *Vil-ERT2* were injected intraperitoneally with a five-day course of tamoxifen starting at P14. Some mice were injected subcutaneously with Fe dextran on P19. At P28, mice were analyzed for: (**a-e**) liver (a), brain (b), bone (c), pancreas (d), and small intestine (e) Mn levels by ICP-OES; (**f**) blood Mn levels by GFAAS; (**g**) bile Mn levels by GFAAS; (**h, i**) Mn absorption by ^52^Mn gavage, with female and male data pooled in (h) and analyzed separately in (i). Data are represented as means +/- standard deviation, with at least four animals per group. Data were tested for normal distribution by Shapiro-Wilk test; if not normally distributed, data were log transformed. Within each sex, groups were compared using two-way ANOVA with Tukey’s multiple comparisons test. (* P<0.05, ** P<0.01, *** P<0.001, **** P<0.0001)

Our observation that intestinal Fpn deficiency minimally impacts Mn levels in wild-type mice is consistent with a previous study^27^. As above, we considered that this may be due to compensatory changes in expression of other Mn transporters. Unlike intestinal Dmt1 deficiency, intestinal Fpn deficiency had minimal impact on duodenum or liver transporter *Slc30a10*, *Slc39a14*, and *Slc39a8* RNA levels in *Slc30a10^+/+^* mice (Fig. S19a, b, S27).

As mentioned above, hepcidin suppresses Fe absorption by negatively regulating Fpn. To determine if increasing hepcidin levels attenuates Mn excess in *Slc30a10^-/-^* mice, we employed anti-sense oligonucleotides (ASOs) against *Tmprss6*, which encodes a transmembrane serine protease that downregulates hepcidin expression^35^. Three weeks of *Tmprss6* ASO treatment in mutant mice, starting at three weeks of age, decreased liver *Tmprss6* RNA levels, increased liver hepcidin RNA levels, and decreased Mn levels in liver and bone but not in brain or small intestine (Fig. S28a-c). Body weights in *Tmprss6* ASO-treated *Slc30a10^-/-^*mice increased to wild-type levels (Fig. S28d, e). However, treatment of mutant mice with *Tmprss6,* but not control, ASOs produced hepatomegaly (Fig. S28f). Decreasing the dose attenuated hepatomegaly but also the impact on Mn levels (data not shown).

### Intestinal deficiency in Dmt1 or Fpn does not impact Mn levels in mice raised on Mn-rich diets

In our final assessment of the role of Dmt1 and Fpn in Mn homeostasis, we determined if intestinal Dmt1 and Fpn contribute to acquired Mn excess. Three-week-old *Dmt1^fl/fl^* and *Fpn^fl/fl^* mice with and without *Vil-ERT2* were treated with tamoxifen and placed on Mn-sufficient or -rich diets. At eight weeks of age, intestinal Dmt1 and Fpn deficiency decreased tissue Fe levels in mice on the Mn-sufficient diet but did not attenuate Mn excess in mice on the Mn-rich diet (Fig. S29).

### Mice with whole-body deficiency in Hjv and Slc30a10 exhibit Fe and Mn excess

While the above data indicate that intestinal Dmt1 and Fpn are essential for Mn absorption and excess in *Slc30a10^-/-^* mice, they do not address the underlying basis for increased Mn absorption in mutant mice. To investigate this, we questioned why aberrant Mn homeostasis does not occur in conditions that share characteristics with SLC30A10 deficiency. Given the key role of intestinal Fpn in Mn absorption and excess in *Slc30a10^-/-^*mice and the fact that hepcidin negatively regulates Fpn, we focused on hereditary hemochromatosis, a disease of Fe excess due to hepcidin deficiency. If Fpn can transport Mn and hepcidin deficiency causes hereditary hemochromatosis, why does hereditary hemochromatosis not result in Mn excess? One possibility is that Fpn does not transport Mn in this condition. Another is that Mn absorption is suppressed. To explore the latter, we considered our previous finding that Slc30a10 exports Mn from enterocytes into the gastrointestinal tract lumen^5^. We hypothesized that SLC30A10 limits Mn absorption in hereditary hemochromatosis by exporting Mn into the gastrointestinal tract lumen, resulting in no net increase in Mn absorption or levels despite the underlying hepcidin deficiency. If correct, this would suggest that loss of intestinal SLC30A10 drives increased Mn absorption in SLC30A10 deficiency.

To explore the role of SLC30A10 in hereditary hemochromatosis, we employed mice with deficiency in hemojuvelin (Hjv), a bone morphogenetic protein coreceptor essential for hepcidin expression^36^. First, we established the impact of whole-body Slc30a10 deficiency on Hjv deficiency. We generated and characterized *Slc30a10 Hjv* mice at five weeks of age, given that most *Slc30a10^-/-^* mice in this cohort did not survive past six weeks of age. Liver *Slc30a10* and *Hjv* RNA levels were decreased as expected (Fig. S30a, b).

We first assessed Mn parameters. Mn levels in most tissues, RBC parameters, and liver *Epo* RNA levels were all increased by Slc30a10 deficiency in *Hjv^+/+^* and *Hjv^-/-^* mice (Fig. S30c-l). Laser ablation inductively coupled plasma mass spectroscopy (LA-ICP-MS) identified increased small intestinal Mn levels in *Slc30a10^-/-^ Hjv^+/+^* and *Slc30a10^-/-^ Hjv^-/-^* mice relative to *Slc30a10^+/+^ Hjv^+/+^*mice (Fig. S31). X-ray fluorescence microscopy showed punctate Mn accumulations in small intestine epithelium in all mutant mice relative to *Slc30a10^+/+^ Hjv^+/+^* mice, although the Mn signal was less pronounced than in *Slc30a10^-/-^*mice (Fig. S32).

We next assessed Fe levels. Liver non-heme Fe levels were increased in *Hjv^-/-^* mice relative to wild-type mice, while levels ranged widely in *Slc30a10^-/-^ Hjv^-/-^* mice and did not differ significantly from other genotypes (Fig. S33a). For pancreas, non-heme Fe levels were increased for *Hjv^-/-^* mice relative to wild-type mice and for *Slc30a10^-/-^ Hjv^-/-^* mice relative to *Slc30a10^-/-^* mice in females only (Fig. S33b). Non-heme Fe levels in spleen, kidney, and small intestine did not differ by genotype (Fig. S33c, data not shown). Diaminobenzidine-enhanced Fe staining showed periportal staining in *Hjv^-/-^* livers, while *Slc30a10^-/-^*and *Slc30a10^-/-^ Hjv^-/-^* livers had variable staining with some displaying diffuse staining and others faint, punctate staining (Fig. S33d-g). The lack of severe Fe perturbations in *Slc30a10^+/+^ Hjv^-/-^* mice may be due to the young age of mice at harvest or the mixed genetic background of *Slc30a10 Hjv* mice, as *Hjv^+/-^*and *Slc30a10^+/-^* mice are maintained on 129S6/SvEvTac and C57BL/6NJ backgrounds respectively.

Finally, we measured metal transport. As expected, liver hepcidin RNA levels were decreased in *Slc30a10^-/-^* females and *Hjv^-/-^*mice relative to wild-type mice and in *Hjv^-/-^ Slc30a10^-/-^* mice relative to *Slc30a10^-/-^* mice (Fig. S34a). ^59^Fe absorption was increased in *Slc30a10^-/-^* mice (males only) and *Hjv^-/-^* mice but not in *Hjv^-/-^ Slc30a10^-/-^* mice (Fig. S34b). The lack of increased ^59^Fe absorption in *Hjv^-/-^ Slc30a10^-/-^* mice is intriguing and may reflect a specific effect of Slc30a10 deficiency on Fe absorption or the impact of metal toxicity on the gut. Bile flow rates and Mn levels did not differ by genotype (Fig. S34c, d). ^52^Mn absorption was increased in all *Slc30a10^-/-^* mice irrespective of Hjv genotype (Fig. S34e).

### Intestinal Slc30a10 deficiency perturbs Mn homeostasis in Hjv deficiency

Having established that whole-body Slc30a10 deficiency increased Mn absorption and levels in *Hjv^-/-^* mice, we next generated *Hjv^-/-^*mice with intestinal Slc30a10 deficiency using floxed *Slc30a10* alleles (*Slc30a10^fl/fl^*) and a *villin* promoter-driven Cre recombinase transgene (*Vil*). Mice were analyzed at two months of age. *Hjv*, hepcidin, and *Slc30a10* RNA levels were decreased as expected by genotype (Fig. 5a-d). Intestinal Slc30a10 deficiency increased Mn levels in multiple tissues (Fig. 5e-j). Mn did not reach *Slc30a10^-/-^* levels but that was expected given that Slc30a10 was inactivated only in intestines. Intestinal Slc30a10 deficiency did not impact bile flow rates in any genotype but did increase bile Mn levels in wild-type and *Hjv^-/-^* mice (data not shown; Fig. 5k). Intestinal Slc30a10 deficiency also increased ^52^Mn absorption in female wild-type, female *Hjv^-/-^*, and male *Hjv^-/-^*mice (Fig. 5l). We next assessed Fe parameters. Intestinal Slc30a10 deficiency did not perturb RBC parameters (Fig. S35a-c). Liver non-heme Fe levels were increased in *Hjv^-/-^*mice irrespective of Slc30a10 genotype (Fig. S35d). Non-heme Fe levels in pancreas or spleen did not differ between genotypes except for increased levels in female intestine Slc30a10-deficient mice (Fig. S35e, f). Diaminobenzidine-enhanced Fe stains of tissues mirrored tissue non-heme Fe levels (Fig. S35g, data not shown). As we postulated above, the lack of perturbation in pancreas and spleen non-heme Fe levels may reflect the mixed genetic background of these mice—*Hjv^+/-^*, *Slc30a10^+/-^*, and *Vil* mice originated on 129S6/SvEvTac, C57BL/6NJ, and C57BL/6J backgrounds respectively. Taken together, these results indicate that intestinal Slc30a10 deficiency increases Mn absorption and Mn levels in *Hjv^-/-^*mice.

**Fig. 5:**
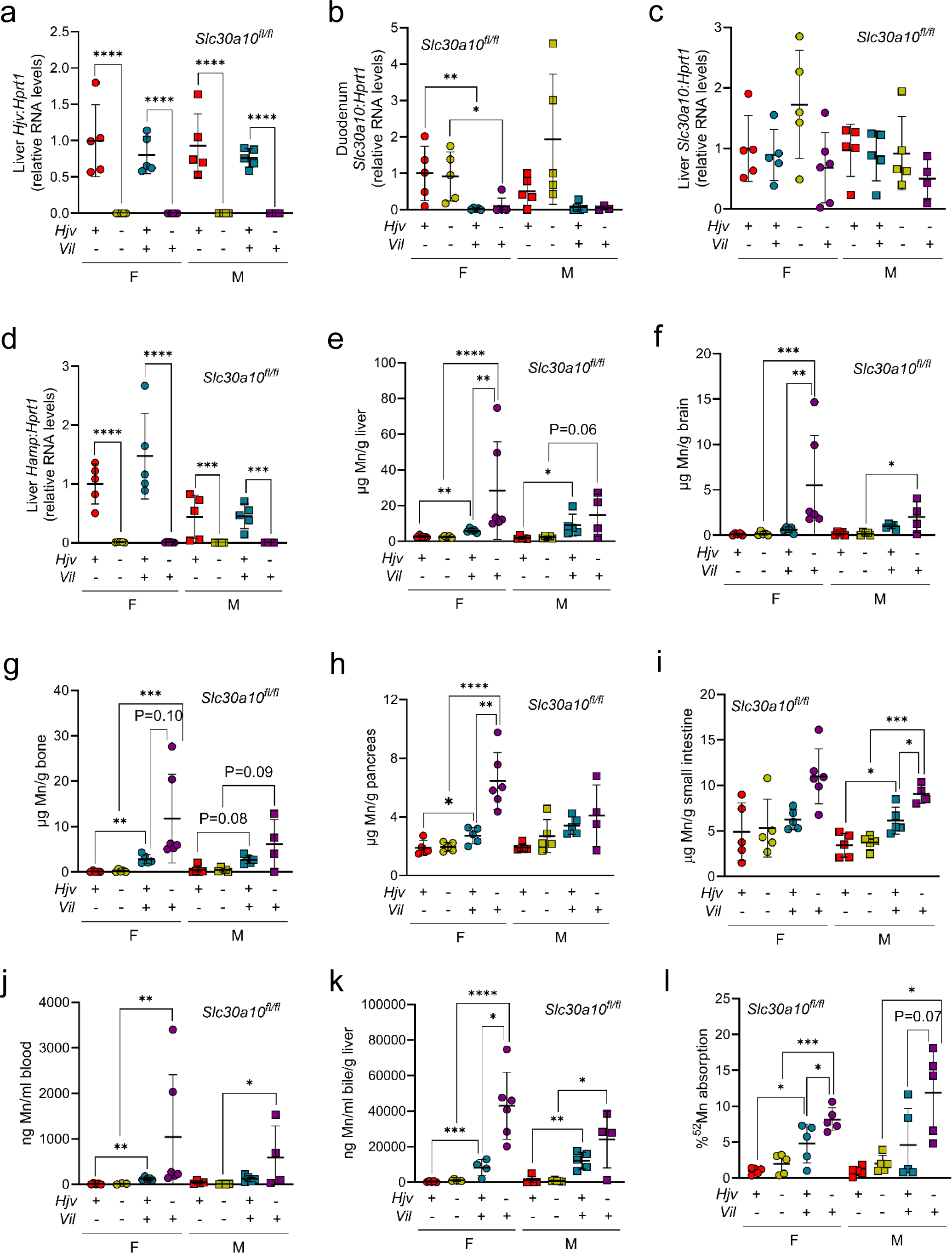
Intestinal Slc30a10 deficiency increases Mn levels in *Hjv^-/-^*mice. Eight-week-old *Hjv Slc30a10^fl/fl^* +/- *Vil* were characterized for: (**a-d**) liver *Hjv* (a), liver *Hamp* (b), duodenum *Slc30a10* (c), and liver *Slc30a10* (d) RNA levels by qPCR; (**e-i**) Mn levels in liver (e), brain (f), bone (g), pancreas (h), and small intestine (i) by ICP-OES; (**j**) blood Mn levels by GFAAS; (**k**) bile Mn levels by GFAAS; (**l**) ^52^Mn absorption by gastric gavage. Data are represented as means +/- standard deviation, with at least five animals per group. Data were tested for normal distribution by Shapiro-Wilk test; if not normally distributed, data were log transformed. Within each sex, groups were compared using two-way ANOVA with Tukey’s multiple comparisons test. (* P<0.05, ** P<0.01, *** P<0.001, **** P<0.0001)

## Discussion

The goal of this study was to interrogate the observation that oral Fe supplementation can be used to treat patients with SLC30A10 deficiency. We first measured Mn absorption in *Slc30a10^-/-^* mice. Intriguingly, it was increased. This phenotype was detected concomitantly with liver Mn excess in pups, suggesting that aberrant Mn absorption is an early event in SLC30A10 deficiency. We next identified the transporters required for increased absorption. Intestinal Dmt1 and Fpn were both essential for increased Mn absorption and excess in *Slc30a10^-/-^* mice. The prominent impact of intestinal Dmt1 and Fpn deficiency on Mn excess indicates that excessive absorption contributes as much as, if not more than, impaired excretion to Mn excess in mutant mice. This represents a shift in our understanding of this disease, which is currently attributed solely to impaired gastrointestinal Mn excretion. Our results explain why oral Fe administration is effective in patients, as this treatment blunts the impact of increased Mn absorption. A recent case report on a patient with SLC30A10 deficiency noted that phlebotomy is contraindicated in this disease given that the resulting Fe deficiency can worsen Mn absorption^37^. Our study provides a mechanistic basis for this concept—phlebotomy-induced Fe deficiency Fe deficiency could further upregulate dietary Fe absorption, thereby worsening Mn absorption by intestinal Dmt1 and Fpn.

In this study, we relied on the use of intestinal Dmt1 and Fpn deficiency to determine if Dmt1 and Fpn play a role in Mn excess in *Slc30a10^-/-^*mice. One inevitable consequence of this approach was Fe deficiency due to impaired Fe absorption. Could the decreased Mn levels in *Slc30a10^-/-^*mice with intestinal Dmt1 or Fpn deficiency reflect Fe deficiency, rather than a direct role for Dmt1 or Fpn in Mn absorption? We injected a subset of mice with Fe dextran to address this specific question. Intestinal Dmt1 and Fpn deficiency still decreased Mn levels in *Slc30a10^-/-^* mice treated with Fe dextran. Furthermore, we did not observe increased bile Mn levels in *Slc30a10^-/-^* mice with intestinal Dmt1 or Fpn deficiency, suggesting that increased excretion was not contributing to decreased Mn levels. This was not unexpected, as there is no precedent for Fe deficiency leading to increased Mn excretion—to the contrary, Fe deficiency is a known risk factor for Mn excess, as it is proposed that upregulation of Fe absorption pathways in response to Fe deficiency leads to increased Mn absorption^38^.

While intestinal Dmt1 and Fpn deficiency had notable impacts on Mn absorption and levels in *Slc30a10^-/-^* mice, they had no effect on these parameters in *Slc30a10^+/+^* mice, similar to previous observations by other groups^19,27^. We propose two potential explanations for this difference in *Slc30a10^-/-^* versus *Slc30a10^+/+^*mice. In the first, intestinal Dmt1 and Fpn transport Mn in *Slc30a10^-/-^*but not *Slc30a10^+/+^* mice. This would require unique cell biology in *Slc30a10^-/-^* enterocytes that confers an essential capacity for Mn transport upon Dmt1 and Fpn. This could involve distinct Mn compartmentalization in *Slc30a10^-/-^* enterocytes. We did observe punctate Mn accumulations in *Slc30a10^-/-^* enterocytes, reminiscent of accumulation within Golgi vesicles in Mn-exposed HeLa cells expressing mutant SLC30A10^39^. In the second explanation, intestinal Dmt1 and Fpn transport Mn in *Slc30a10^+/+^*and *Slc30a10^-/-^* mice but only in an essential manner in the latter. This would require that other transporters can absorb Mn in wild-type mice. SLC39A8 is a viable candidate, as this apical Mn importer is expressed in intestines and mice with intestinal Slc39a8 deficiency do have decreased absorption of gavaged ^54^Mn^31,40^. Both explanations will be explored in future studies.

While the first half of our study focused on the role of intestinal Dmt1 and Fpn in Mn absorption and excess in *Slc30a10^-/-^*mice, the second half of our study asked why mutant mice develop increased Mn absorption at all. To answer this question, we exploited the fact that hereditary hemochromatosis, a disease of Fe excess due to aberrant upregulation of Fpn-dependent Fe absorption, is not associated with Mn excess. In 2019, we demonstrated in mice that hepatocyte Slc30a10 is essential for biliary Mn excretion and that intestinal Slc30a10 exports Mn when hepatic Slc30a10 is lacking^5^. More recently, we showed that AAV-mediated, liver-specific SLC30A10 overexpression is sufficient to attenuate Mn excess and normalize erythropoietin levels, RBC parameters, and liver and brain gene expression in *Slc30a10^-/-^* mice^28^. Together, these two studies indicate that hepatic Slc30a10 is sufficient to carry out Mn excretion. If intestinal Slc30a10 plays a secondary role in Mn excretion, what is its primary role? In this study, we hypothesized that intestinal Slc30a10 functions to attenuate Mn absorption when Fe absorption pathways are upregulated. *Hjv^-/-^* mice with intestinal Slc30a10 deficiency did indeed have increased Mn absorption and levels. Absorption and levels were not increased to the extent observed with whole body Slc30a10 deficiency, but this was expected given that mice with intestinal Slc30a10 deficiency still expressed hepatic Slc30a10. Overall, our results supported the notion that intestinal Slc30a10 minimizes Mn absorption under conditions of hepcidin deficiency, which explains why Mn absorption is increased in *Slc30a10^-/-^* mice: intestinal Slc30a10 is not present to attenuate Mn absorption when Fe absorption pathways are upregulated due to the increased demand for Fe to supply erythropoiesis.

While loss-of-function SLC30A10 mutations are exceedingly rare, non-coding variants in SLC30A10 are much more common and are one of the key determinants of blood Mn levels in otherwise healthy individuals^41–44^. We speculate that SLC30A10 polymorphisms could modify Mn levels in individuals with hepcidin deficiency in diseases such as hereditary hemochromatosis. Polymorphisms that enhance SLC30A10 expression would minimize the development of Mn excess, while those that dampen expression would promote it. Hereditary hemochromatosis is known for its clinical variability and it is possible that SLC30A10 polymorphisms could contribute to this variability^45^. Another condition of potential relevance to SLC30A10 and Mn in settings of aberrant Fe homeostasis is dietary Fe deficiency, the most common nutritional disorder in the world^46^. Notably, Fe deficiency is a risk factor for development of acquired Mn excess^47^. The mechanism of Mn excess in dietary Fe deficiency has yet to be reported but is ascribed to upregulation of Fe absorption pathways leading to increased Mn absorption. As with hereditary hemochromatosis, we speculate that SLC30A10 polymorphisms will impact the susceptibility of individuals with dietary Fe deficiency to development of Mn excess. While we would not expect severe Mn excess in this situation, even moderate increases in blood Mn levels correlate with impaired cognition and therefore may be relevant to the long-term health of individuals with Fe deficiency^48,49^.

We recently reported that Mn-induced Hif2 upregulation in livers of *Slc30a10^-/-^* mice perturbs expression of multiple genes including hepcidin and erythropoietin^8^. We also observed that Hif2a deficiency attenuated Mn excess in *Slc30a10^-/-^* mice. Our current study explains this latter result. Hepatic Hif2a deficiency corrected erythropoietin excess and hepcidin deficiency, thereby attenuating Fe absorption pathways and limiting Mn absorption in mutant mice. Using the results from that study and the present one, we now propose a revised model for Mn excess in SLC30A10 deficiency (Fig. 6). Loss of SLC30A10-dependent gastrointestinal Mn excretion increases Mn levels. Liver Mn excess increases HIF2 levels and erythropoietin synthesis. Erythropoietin excess stimulates erythropoiesis, which indirectly suppresses liver hepcidin expression and stimulates Fe absorption by DMT1 and FPN. Without intestinal SLC30A10 present to export Mn back into the gastrointestinal tract lumen, DMT1 and FPN also mediate increased Mn absorption, furthering exacerbating systemic Mn excess.

**Fig. 6:**
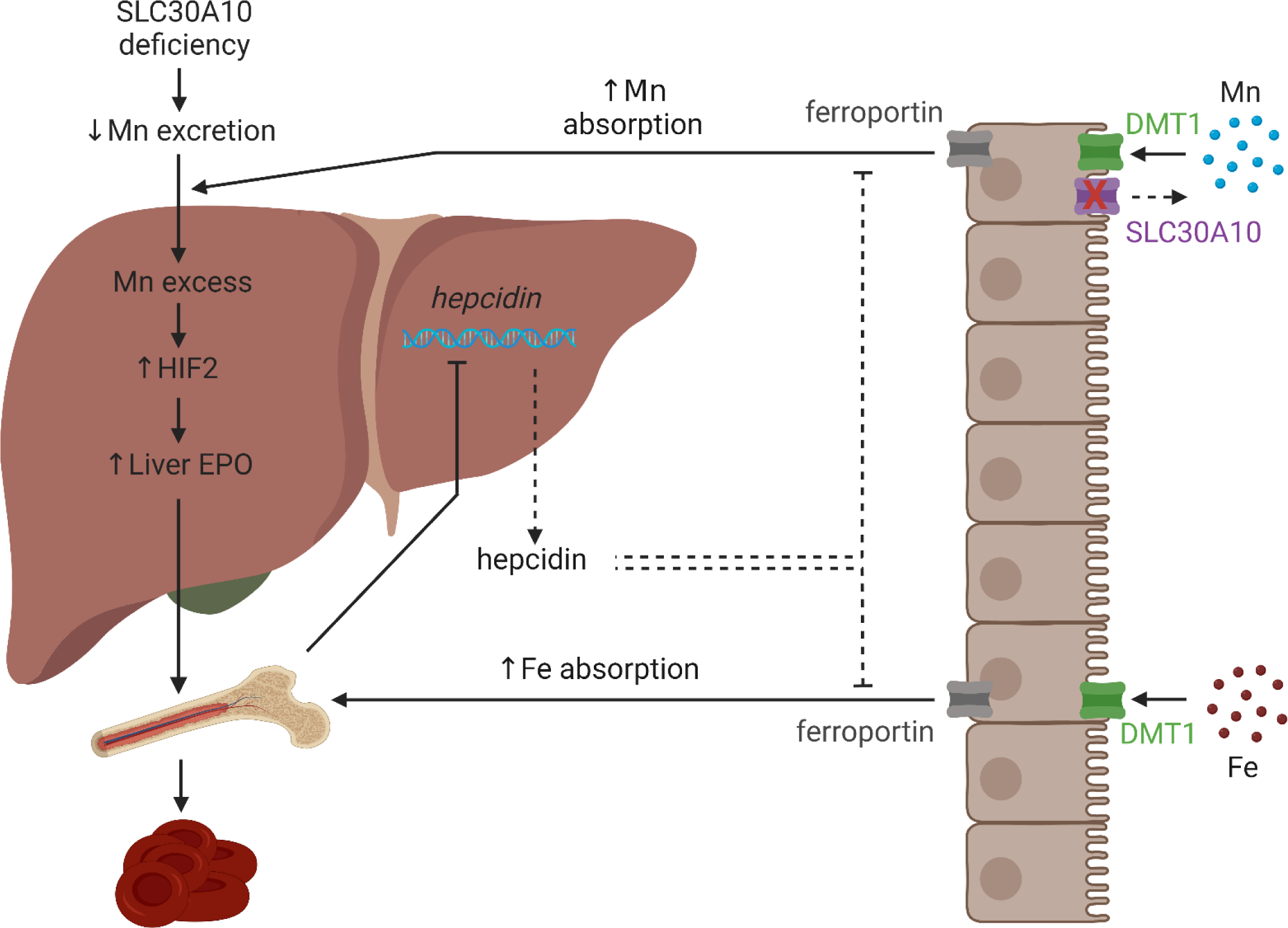
Model of Mn excess in SLC30A10 deficiency. Loss of SLC30A10-dependent gastrointestinal Mn excretion increases Mn levels. Liver Mn excess increases HIF2 levels and erythropoietin (EPO) synthesis. EPO excess stimulates erythropoiesis, which indirectly suppresses liver hepcidin expression and stimulates Fe absorption by DMT1 and FPN. Without intestinal SLC30A10 present to export Mn back into the gastrointestinal tract lumen, DMT1 and FPN also mediate increased Mn absorption, furthering exacerbating systemic Mn excess. Created with Biorender.com

Overall, our study has produced several key observations. First, SLC30A10 deficiency is not just a disease of impaired gastrointestinal Mn excretion—excessive Mn absorption also contributes prominently. Second, DMT1 and FPN can transport Mn *in vivo*. Third, while hepatic SLC30A10 is essential for biliary Mn excretion, intestinal SLC30A10 minimizes Mn absorption when Fe absorption pathways are upregulated. These results expand our understanding of Fe and Mn homeostasis and the overlaps between the two. While the relevance of DMT1 and FPN to Mn homeostasis is still contentious in the literature, we propose that a reconsideration of the role of these Fe transporters in Mn transport is warranted based upon our results. Even if DMT1 and FPN are only essential for Mn transport in SLC30A10 deficiency, determining why this occurs would enhance our understanding of the mechanisms by which metal specificity is determined for Fe transporters *in vivo* and, more broadly, enhance our understanding of how the body balances absorption of two nutrients when overlaps can exist in their means of transport.

## Methods

### Mouse care, generation, and treatment and sample collection

Mice were bred and maintained in the animal care facility at Brown University. Mouse studies were approved by the Institutional Animal Care and Use Committee at Brown University. Mice were group-housed in ventilated cage racks, maintained on a 12-hour light-dark cycle with controlled temperature and humidity, and provided standard chow (LabDiet 5010 containing 120 ppm Mn) and water *ad libitum* unless otherwise specified. Littermates of the same sex were randomly assigned to experimental groups. *Slc30a10^-/-^*and *Slc30a10^fl/fl^* mice were generated as previously described and maintained by backcrossing onto C57BL/6NJ mice (Jackson Laboratory #005304)^5^. *Hjv^+/-^*mice (Jackson Laboratory #017788) were maintained by backcrossing onto 129S6/SvEvTac (Taconic Biosciences #129SVE). *Slc30a10^+/+^* and *Slc30a10^-/-^*mice were generated by crossing *Slc30a10^+/-^* mice. To generate mice with inducible intestinal deficiency in Dmt1 or Fpn, *Slc30a10^+/fl^*mice were bred to mice carrying floxed *Dmt1* (*Dmt1^fl/fl^*, Jackson Laboratory #017789) or *Fpn* (*Fpn^fl/fl^*, Jackson Laboratory #017790) alleles and a tamoxifen-inducible, villin promoter-driven Cre recombinase transgene (*Vil-ERT2*, Jackson Laboratory #020282) in a multi-step breeding scheme. *Slc30a10^+/-^ Dmt1^fl/fl^*and *Slc30a10^+/-^ Dmt1^fl/fl^ Vil-ERT2* mice or *Slc30a10^+/-^ Fpn^fl/fl^* and *Slc30a10^+/-^ Fpn^fl/fl^ Vil-ERT2* mice were crossed to generate mice for characterization. Mice were injected intraperitoneally with tamoxifen (75 mg/kg) for five days at P14 or P21 to inactivate floxed alleles. Treatment with corn oil, the solvent used for tamoxifen, had no impact on liver Mn levels in mutant mice (data not shown). A subset of mice were also injected subcutaneously with Fe dextran treatments (100 mg/kg) at P19. *Hjv Slc30a10* mice were generated by crossing *Hjv^+/-^* and *Slc30a10^+/-^*mice, then crossing *Hjv^+/-^ Slc30a10^+/-^* mice. To generate *Hjv* mice with intestinal Slc30a10 deficiency, mice carrying a *villin* promoter-driven Cre recombinase transgene (*Vil*, Jackson Laboratory #004586) were employed. *Hjv^+/-^*, *Slc30a10^fl/fl^*, and *Vil* mice were bred together in a multi-step breeding scheme, with mice for study generated by crossing *Hjv^+/-^ Slc30a10^fl/fl^*and *Hjv^+/-^ Slc30a10^+/fl^ Vil* mice.

For ASO-based studies, *Slc30a10^+/+^* mice were treated with sterile-filtered PBS and *Slc30a10^-/-^* mice were treated with sterile-filtered GalNAc-control or *Tmprss6* ASO (10 µl/gram body weight, stock 0.25 mg/ml, Ionis Pharmaceuticals). ASOs were administered by intraperitoneal injection twice a week beginning at weaning until six weeks of age for a total of 12 doses. Animals were harvested two days after the last dose. To raise mice on Mn-deficient diets, mice were weaned onto diets containing 1, 100, or 2400 ppm Mn (Envigo TD.140497, TD.140499, TD.180124).

To collect samples for analysis, mice were anesthetized with isoflurane then underwent bile, blood, and tissue collection. Bile was collected surgically by ligation of the common bile duct, cannulation of the gallbladder, and collection over 60 minutes as previously described^5^. Bile volumes were measured every five minutes to assess bile flow rates. Blood was then collected by cardiac puncture into EDTA-coated tubes (BD) and serum collection tubes (BD). Mice were euthanized by decapitation and tissues collected for metal, DNA, RNA, and protein analysis. Complete blood counts were performed on freshly isolated anticoagulated blood using VetAbcPlus (Sci).

### RNA analysis

For gene-specific qPCR, 100-200 mg tissue was homogenized in TRIzol (ThermoFisher) using 0.5 mm zirconium beads and Bullet Blender (Next Advance), followed by chloroform extraction, isopropanol precipitation, and 70% ethanol wash, as previously described^5^. Standards were made by serially diluting mixtures of control and experimental samples, then processed identically as experimental samples. Samples underwent DNase treatment and cDNA synthesis using the High Capacity cDNA Reverse Transcription Kit with RNase Inhibitor (ThermoFisher). qPCR was performed using PowerUP SYBR Green Master Mix (ThermoFisher). Primers listed below were designed using Primer3 and Geneious or used from previously published literature. Amplicon fidelity was confirmed by Sanger sequencing. Primer concentrations were optimized by testing a series of forward and reverse primer combinations followed by melt curve analyses. Primers used were as follows: *Hprt1* (5’GCCCCAAAATGGTTAAGGTT, 5’TGGCAACATCAACAGGACTC, exons 6-9); *Slc30a10* (5’AGAGACTGCTGTCATCTTGCTG, 5’TGTGCACTTCATGCACACTG, exons 3-4); *Dmt1* (5’TCCTCATCACCATCGCAGACACTT, 5’TCCAAACGTGAGGGCCATGATAGT, exons 7-8); *Fpn* (5’CTTGCTCTGGAAGGTTTACC, 5’TGGAGTCTTTCTCACCCATT, exons 6-7); *Slc39a14* (5’GGAACCCTCTACTCCAACGC, 5’ATGGTTATGCCCGTGATGGT, exons 5-7); *Slc39a8* (5’CAACGCAAAGCCCAGTCTTT, 5’GCGTTTGAGAAAAGAGTCCCAA, exons 3-4); *Hamp* (5’TGTCTCCTGCTTCTCCTCCT, 5’CTCTGTAGTCTGTCTCATCTGTTG, exons 1-2); *Epo* (5’ACTCTCCTTGCTACTGATTCCT, 5’ATCGTGACATTTTCTGCCTCC, exons 2-3); *Hjv* (5’ACGACTTCCTCTTTGTCCAGGC, 5’TCCACCTCAGCCTGGTAGACTT, exons 3-4).

For bulk RNA-seq, RNA extraction, library preparation, sequencing, and analysis was conducted at Azenta Life Sciences (South Plainfield, NJ, USA). Total RNA was extracted from fresh frozen tissue samples using Qiagen RNeasy Plus Universal mini kit (Qiagen). RNA samples were quantified using Qubit 2.0 Fluorometer (Life Technologies) and RNA integrity was checked using TapeStation 4200 (Agilent Technologies). RNA sequencing libraries were prepared using the NEBNext Ultra II RNA Library Prep for Illumina (NEB). Briefly, mRNAs were initially enriched with Oligod(T) beads. Enriched mRNAs were fragmented for 15 minutes at 94 °C. First strand and second strand cDNA were subsequently synthesized. cDNA fragments were end-repaired and adenylated at 3’ ends and universal adapters were ligated to cDNA fragments, followed by index addition and library enrichment by PCR with limited cycles. Sequencing libraries were validated on TapeStation 4200 and quantified using Qubit 2.0 Fluorometer (Invitrogen) as well as by quantitative PCR (KAPA Biosystems). Sequencing libraries were clustered on a flowcell, which was then loaded on the Illumina instrument (4000 or equivalent). Samples were sequenced using a 2x150bp Paired End configuration. Image analysis and base calling were conducted by the Control software. Raw sequence data (.bcl files) generated by the sequencer were converted into fastq files and de-multiplexed using Illumina’s bcl2fastq 2.17 software. One mismatch was allowed for index sequence identification. After investigating the quality of raw data, sequence reads were trimmed to remove possible adapter sequences and nucleotides with poor quality. Trimmed reads were mapped to the reference genome available on ENSEMBL using the STAR aligner v.2.5.2b, a splice aligner that detects splice junctions and incorporates them to help align the entire read sequences. BAM files were generated as a result of this step. Unique gene hit counts were calculated by using feature Counts from the Subread package v.1.5.2. Only unique reads that fell within exon regions were counted. After extraction of gene hit counts, the gene hit counts table was used for downstream differential expression analysis. Using DESeq2, a comparison of gene expression between the groups of samples was performed. The Wald test was used to generate p-values and Log2 fold changes. Genes with adjusted p-values < 0.05 and absolute log2 fold changes > 1 were called as differentially expressed genes for each comparison. A PCA analysis was performed using the "plotPCA" function within the DESeq2 R package. The top 500 genes, selected by highest row variance, were used to generate PCA plots. Venn diagrams were generated using bioinformatics.psb.ugent.be/webtools/Venn/. Gene ontology was performed using ShinyGO at http://bioinformatics.sdstate.edu/go/^50^. Heatmaps on rlog-transformed gene counts were generated using Morpheus at https://software.broadinstitute.org/morpheus/.

### Immunofluorescence

Formalin-fixed, paraffin-embedded tissue sections were deparaffinized in xylene and serially re-hydrated in ethanol. Antigen retrieval was performed using citrate- (for Fpn; Vector Laboratories H-3300-250) or Tris- (for Dmt1; Vector Laboratorie H-3301-250) based buffers in a rice steamer for 20 minutes. Slides were then blocked for 1 hour at room temperature in 10% goat serum/1% BSA/PBS followed by overnight incubation with/without primary antibody (Dmt1: Proteintech #20507-1-AP lot 0014129 at 1:500; Fpn: Novus Bio #NBP1-21502 lot D127012-2 at 1:200) in a humidified chamber at 4 °C. Slides were then washed for 2 minutes in PBS 3 times then incubated for 2 hours with goat anti–rabbit IgG/Alexa Fluor 594 secondary antibody (Thermo Fisher Scientific A-11012 at 1:1000) in blocking buffer at room temperature. Slides were washed for 2 minutes in PBS 3 times, then coverslips were applied using mounting media containing DAPI (Vector Labs H-1800-10). Slides were scanned with an Olympus VS200 slide scanner with 10x objective.

### Metal analysis

For measurement of tissue total metal levels, 10-200 mg tissue were digested in 1000 µl 70% trace-metal-grade nitric acid (Fisher) at 65°C for two-three hours then diluted 25-fold in MilliQ water (Millipore Sigma) and analyzed by inductively coupled plasma optical emission spectroscopy (ICP-OES) on an iCAP 7400 DUO (ThermoScientific) or graphite furnace atomic absorption spectroscopy (GFAAS) on an AAnalyst 600 (Perkin Elmer). For measurement of blood metal levels, samples were digested with two volumes nitric acid at 65 °C for four hours, diluted 25-fold with water, then analyzed by GFAAS. For measurement of bile metal levels, samples were digested twice with one volume nitric acid at 65 °C until dry, twice with one volume hydrogen peroxide at 65 °C until dry, resuspended in 2% nitric acid, then analyzed by GFAAS. For ICP-OES, a series of standards were analyzed and sample values were extrapolated from the generated curve. A quality control standard (IV-28, Inorganic Ventures) was run every ten samples to assess changes in sensitivity. If tissue size was small or metal levels too low, GFAAS was used. For GFAAS, standards were measured to create a standard curve. To correct for variations in sensitivity, quality control standard SRM 1640a (NIST) was analyzed every ten samples. Irrespective of instrumentation, a correction curve was calculated based on quality control analysis and correction factor applied to each sample to control for changes in instrument sensitivity.

For measurement of tissue non-heme iron levels, 10-200 mg tissue was digested in 1 ml 3 N hydrochloric acid (Fisher)/10% trichloroacetic acid (Millipore Sigma) at 65 °C for two days, with 30 minutes vortexing each day, followed by centrifugation. Iron levels were measured by mixing 10 μl supernatants with 200 μl chromagen (five volumes MilliQ water; five volumes saturated sodium acetate (Fisher); one volume chromagen stock, consisting of 0.1% bathophenanthroline sulfonate (Millipore Sigma) and 1% thioglycolic acid (Millipore Sigma)) in a 96-well plate. Iron standards (Fisher) were included. After a ten-minute incubation, absorbances were measured at 535 nm. Mock digests without samples were included for this and all other metal analyses.

For X-ray fluorescence microscopy (XFM), data were collected at beamlines 2-IDE and 2-IDD at the Advanced Photon Source part of the Argonne National Laboratory. OCT-embedded tissue was sectioned to 10 µm thickness using a cryostat, placed directly onto 4 µm thick Ultralene and air dried. The tissue containing Ultralene was glued onto an in-house made sample holder. At the beamline, visual light microscope images were collected of the tissue sections and coordinates saved to beamline computers using the program uProbeX (unpublished, Arthur Glowacki). Coordinates for regions of interest were selected in uProbeX. A double layer monochromator produced an incident energy of 10 keV and samples were raster scanned in on-the-fly mode. X-ray fluorescence photons were collected with an energy dispersive silicon drift detector (Vortex). High resolution scans were recorded at 500 nm step size. Data were background subtracted and fitted pixel by pixel using the software MAPS^51^. Quantitation was performed by calibrating the X-ray fluorescence yield of each element to that of a calibration standard (Axo, Dresden).

For laser ablation inductively-coupled mass spectroscopy (LA-ICP-MS), samples were analyzed on an imageBIO266 laser (ESL) using a TV3 sample chamber and coupled with a Vitesse TOF-ICP-MS (Nu Instruments) at the Biomedical National Elemental Imaging Resource (BNEIR) at Dartmouth College. Helium was used as the transport gas, flowing through the laser chamber at 200 ml/min and the laser cup at 200 ml/min. On the Vitesse, the nebulizer gas flow was set to 1250 ml/min and the reaction cell gases, hydrogen and helium, were set to 6 and 17 ml/min respectfully. Elemental images were collected with an 8 µm spot size, a 4 µm spot overlap, a 400 Hz rep rate, and a laser fluence of 3.5 J/cm^2^. Data was exported to Iolite for image processing.

To assess metal absorption, mice were fasted in metabolic cages (Tecniplast) for four hours to clear the upper gastrointestinal (GI) tract of most contents prior to gavage. To assess Mn absorption, mice were gavaged with 1 µCi ^54^MnCl_2_ (Perkin Elmer) or ^52^MnCl_2_ (University of Wisconsin-Madison) then returned to the metabolic cages for 15 minutes. (Perkin Elmer discontinued ^54^MnCl_2_ production midway through this study.) To assess Fe absorption, mice were gavaged with 1 µCi ^59^FeCl_2_ (Perkin Elmer) in 2.21 mM FeCl_3_ in 1 M ascorbic acid then returned to the metabolic cage for one hour. Mice were then anesthetized with isoflurane and euthanized by cervical dislocation. GI tracts and gallbladders were removed, then GI tracts, gallbladders, and carcasses were counted using a Triathler Gamma Counter with external NaI well-type crystal detector (Hidex). GI tracts were then separated into stomach, small intestine, cecum, and large intestine and each compartment cleaned and rinsed. Percent absorption was calculated as sum of radioactivity in whole carcass as percent of total radioactivity counted.

To perform histologic assessment of tissue Fe, formalin-fixed, paraffin-embedded tissue sections were deparaffinized in xylene, serially re-hydrated in ethanol, followed by staining with Iron Stain kit (Millipore Sigma). Slides were then subjected to diaminobenzidine enhancement (Millipore Sigma) followed by dehydration with ethanol and xylene. Coverslips were mounted using Cytoseal XYL mounting media (Fisher). Slides were scanned with Olympus VS200 slide scanner with 20x objective.

### Statistics

Statistics were performed using GraphPad Prism 9. Data were tested for normal distribution by Shapiro-Wilk test. If not normally distributed, data were log transformed. Groups within each sex were compared by one-or two-way ANOVA with Tukey’s multiple comparisons test or by unpaired, two-tailed t test as indicated in figure legends. P<0.05 was considered significant. Data are represented as means +/- standard deviation.

### Study approval

Mouse studies were approved by the Institutional Animal Care and Use Committee at Brown University.

### Data availability

All non-sequencing quantitative data are contained within the “Supporting data values” XLS file. For sequencing data, BAM files can be found under accessions PRJNA1111893 on the NCBI Sequence Read Archive.

### Author contributions

Study design: MP, TBB; execution of experiments and data analysis: MP, JZZ, GSC, LC, CJM, HLK, OA, BL, MR, BPJ, TP, SG, MA, TBB; manuscript writing: TBB; manuscript revision: all authors.

## Acknowledgements

Funding: NIH DK84122 (T.B.B.), DK110049 (T.B.B.), DK117524 (C.J.M.), ES007272-24 T32 (C.J.M.), GM077995 T32 (H.L.C.). We thank Christoph Schorl and the Genomics Core for assistance with qPCR, and Joseph Orchardo, David Murray, and Marcelo da Rosa Alexandre for assistance with metal analysis. The Genomics Core has received partial support from the National Institutes of Health (P30GM103410, P30RR031153, P20RR018728, S10RR02763), National Science Foundation (EPSCoR 0554548), Lifespan Rhode Island Hospital, Brown University’s Division of Biology and Medicine and Provost’s office. Use of the Advanced Photon Source, an Office of Science User Facility operated for the U.S. Department of Energy (DOE) Office of Science by Argonne National Laboratory, was supported by the U.S. DOE under Contract No. DE-AC02-06CH11357. LA-ICP-MS work at Dartmouth was supported by NIH OD032352. The Dartmouth Biomedical National Elemental Imaging Resource is supported by NIH GM141194 and S10ODO32352.

---

**Fig. S1. Small intestine epithelial cells in *Slc30a10^-/-^*mice have excess Mn.** Additional images from x-ray fluorescence of small intestinal sections from two-month-old *Slc30a10^+/+^* and *Slc30a10^-/-^*mice are shown for phosphorus (P), Mn, zinc (Zn), sulfur (S), Fe, potassium (K), and copper (Cu).

**Fig. S2-5. Enterocyte Dmt1 protein levels are increased in *Slc30a10^-/-^*mice.** Fixed small intestine sections from two-month-old female *Slc30a10^+/+^*(Fig. S2), male *Slc30a10^+/+^* (Fig. S3), female *Slc30a10^-/-^*(Fig. S4), and male *Slc30a10^-/-^* (Fig. S5) mice were analyzed for Dmt1 expression by immunofluorescence. Red signal corresponds to anti-Dmt1 antibody binding. Dmt1 signal on apical enterocyte membranes in mutant mice indicated by white arrowheads in Fig. S4 and S5. Blue signal represents DAPI staining. Images on left correspond to 10x magnification with scale bar set to 50 μm; images on right correspond to magnified view of area outlined by white box in left images, with scale bar set to 20 μm.

**Fig. S6-9. Enterocyte Fpn protein levels are increased in *Slc30a10^-/-^*mice.** Fixed small intestine sections from two-month-old female *Slc30a10^+/+^*(Fig. S6), male *Slc30a10^+/+^* (Fig. S7), female *Slc30a10^-/-^*(Fig. S8), and male *Slc30a10^-/-^* (Fig. S9) mice were analyzed for Fpn expression by immunofluorescence. Red signal corresponds to anti-Fpn antibody binding. Fpn signal on basolateral enterocyte membranes in mutant mice indicated by white arrowheads in Fig. S8 and S9. Blue signal represents DAPI staining. Images on left correspond to 10x magnification with scale bar set to 50 μm; images on right correspond to magnified view of area outlined by white box in left images, with scale bar set to 20 μm.

**Fig. S10: Tamoxifen treatment of *Slc30a10^+/+^ Dmt1^fl/fl^ Vil-ERT2* at P14 decreases intestinal *Dmt1* RNA levels.** *Slc30a10 Dmt1^fl/fl^* mice with and without *Vil-ERT2* were injected intraperitoneally with a five-day course of tamoxifen starting at P14. Some mice were injected subcutaneously with Fe dextran on P19. At P28, mice were analyzed for duodenum (**a**) and liver (**b**) *Slc30a10* and duodenum (**c**) and liver (**d**) *Dmt1* RNA levels (relative to *Hprt1* RNA levels) by qRT-PCR. Data are represented as means +/- standard deviation, with at least four animals per group. Data were tested for normal distribution by Shapiro-Wilk test; if not normally distributed, data were log transformed. Within each sex, groups were compared using two-way ANOVA with Tukey’s multiple comparisons test. (* P<0.05, ** P<0.01, *** P<0.001, **** P<0.0001)

**Fig. S11-14: Tamoxifen treatment of Slc30a10 Dmt1^fl/fl^ Vil-ERT2 at P14 decreases intestinal Dmt1 protein levels.** Slc30a10^+/+^ Dmt1^fl/fl^ (Fig. S11), *Slc30a10^+/+^ Dmt1^fl/fl^ Vil-ERT2* (Fig. S12), *Slc30a10^-/-^ Dmt1^fl/fl^* (Fig. S13), and *Slc30a10^-/-^ Dmt1^fl/fl^ Vil-ERT2* (Fig. S14) mice were injected intraperitoneally with a five-day course of tamoxifen starting at P14. At P28, mice were analyzed for Dmt1 expression by immunofluorescence of fixed small intestine sections. Red signal corresponds to anti-Dmt1 antibody binding. Dmt1 signal on apical enterocyte membrane indicated in Fig. S13 by white arrowheads. Blue signal represents DAPI staining. Images on left correspond to 10x magnification with scale bar set to 50 μm; images on right correspond to magnified view of area outlined by white box in left images, with scale bar set to 20 μm.

**Fig. S15: Tamoxifen treatment of *Slc30a10^+/+^ Dmt1^fl/fl^ Vil-ERT2* at P14 leads to Fe deficiency.** *Slc30a10 Dmt1^fl/fl^* mice with and without *Vil-ERT2* were injected intraperitoneally with a five-day course of tamoxifen starting at P14. Some mice were injected subcutaneously with Fe dextran on P19. At P28, mice were analyzed for: (**a**) liver non-heme Fe levels by bathophenanthroline-based assay; (**b**) liver *Hamp* RNA levels; (**c-e**) RBC counts (c), hemoglobin levels (d), and hematocrits (e) by complete blood counts; (**f, g**) liver copper (Cu) (f) and zinc (Zn) (g) by ICP-OES. Data are represented as means +/- standard deviation, with at least four animals per group. Data were tested for normal distribution by Shapiro-Wilk test; if not normally distributed, data were log transformed. Within each sex, groups were compared using two-way ANOVA with Tukey’s multiple comparisons test. (* P<0.05, ** P<0.01, *** P<0.001, **** P<0.0001)

**Fig. S16: Intestinal Dmt1 deficiency, initiated at P14, modestly impacts whole brain gene expression in *Slc30a10^-/-^* mice.** (**a-c**) Bulk RNA-seq was performed on whole brain samples from three female *Slc30a10^-/-^ Dmt1^fl/fl^* and *Slc30a10^+/+^ Dmt1^fl/fl^*, mice then assessed by similarity analysis (a), principal component analysis (b), and volcano plot (c) where genes more abundantly expressed in *Slc30a10^-/-^ Dmt1^fl/fl^* mice have log_2_(fold change)>1. (**d-f**) Bulk RNA-seq was performed on whole brain samples from three female *Slc30a10^-/-^ Dmt1^fl/fl^ Vil-ERT2* and *Slc30a10^-/-^ Dmt1^fl/fl^* mice then assessed by similarity analysis (d), principal component analysis (e), and volcano plot (f) where genes more abundantly expressed in *Slc30a10^-/-^ Dmt1^fl/fl^ Vil-ERT2* mice have log_2_(fold change)>1. In volcano plots, differentially expressed genes (adjusted P value<0.05 and absolute value of log_2_(fold change)>1) are shown as light orange points with gene names shown adjacent as space permitted; non-differentially expressed genes are shown as blue points.

**Fig. S17: Intestinal Dmt1 deficiency, initiated at P14, modestly impacts whole brain gene expression in *Slc30a10^-/-^* mice.** Venn analysis was performed on RNA-seq results from Fig. S16. Heatmaps are shown for 67 genes differentially expressed only in *Slc30a10^-/-^ Dmt1^fl/fl^*vs. *Slc30a10^+/+^ Dmt1^fl/fl^*, mice, 7 genes differentially expressed only in *Slc30a10^-/-^ Dmt1^fl/fl^ Vil-ERT2* and *Slc30a10^-/-^ Dmt1^fl/fl^*mice, and 9 genes differentially expressed in both comparisons.

**Fig. S18: Tamoxifen treatment of *Slc30a10^+/+^ Dmt1^fl/fl^ Vil-ERT2* at P14 increases liver *Slc39a14* and *Slc39a8* RNA levels.** *Slc30a10 Dmt1^fl/fl^* mice with and without *Vil-ERT2* were injected intraperitoneally with a five-day course of tamoxifen starting at P14. Some mice were injected subcutaneously with Fe dextran on P19. At P28, mice were analyzed for duodenum (**a**) and liver (**b**) *Slc39a14* and duodenum (**c**) and liver (**d**) *Slc39a8* RNA levels (relative to *Hprt1* RNA levels) by qRT-PCR. Data are represented as means +/- standard deviation, with at least four animals per group. Data were tested for normal distribution by Shapiro-Wilk test; if not normally distributed, data were log transformed. Within each sex, groups were compared using two-way ANOVA with Tukey’s multiple comparisons test. (* P<0.05, ** P<0.01, *** P<0.001, **** P<0.0001)

**Fig. S19: Tamoxifen treatment of *Slc30a10^+/+^ Fpn^fl/fl^ Vil-ERT2* at P14 decreases intestinal *Fpn* RNA levels.** *Slc30a10 Fpn^fl/fl^* mice with and without *Vil-ERT2* were injected intraperitoneally with a five-day course of tamoxifen starting at P14. Some mice were injected subcutaneously with Fe dextran on P19. At P28, mice were analyzed for duodenum (**a**) and liver (**b**) *Slc30a10* and duodenum (**c**) and liver (**d**) *Fpn* RNA levels (relative to *Hprt1* RNA levels) by qRT-PCR. Data are represented as means +/- standard deviation, with at least four animals per group. Data were tested for normal distribution by Shapiro-Wilk test; if not normally distributed, data were log transformed. Within each sex, groups were compared using two-way ANOVA with Tukey’s multiple comparisons test. (* P<0.05, ** P<0.01, *** P<0.001, **** P<0.0001)

**Fig. S20-23: Tamoxifen treatment of Slc30a10 Fpn^fl/fl^ Vil-ERT2 at P14 decreases intestinal Fpn protein levels.** Slc30a10^+/+^ Fpn^fl/fl^ (Fig. S20), *Slc30a10^+/+^ Fpn^fl/fl^ Vil-ERT2* (Fig. S21), *Slc30a10^-/-^ Fpn^fl/fl^* (Fig. S22), and *Slc30a10^-/-^ Fpn^fl/fl^ Vil-ERT2* (Fig. S23) mice were injected intraperitoneally with a five-day course of tamoxifen starting at P14. At P28, mice were analyzed for Fpn expression by immunofluorescence of fixed small intestine sections. Red signal corresponds to anti-Fpn antibody binding. Fpn signal on basolateral enterocyte membrane indicated in Fig. S20 and S22 by white arrowheads. Blue signal represents DAPI staining. Images on left correspond to 10x magnification with scale bar set to 50 μm; images on right correspond to magnified view of area outlined by white box in left images, with scale bar set to 20 μm.

**Fig. S24: Tamoxifen treatment of *Slc30a10^+/+^ Fpn^fl/fl^ Vil-ERT2* at P14 leads to Fe deficiency.** *Slc30a10 Fpn^fl/fl^*mice with and without *Vil-ERT2* were injected intraperitoneally with a five-day course of tamoxifen starting at P14. Some mice were injected subcutaneously with Fe dextran on P19. At P28, mice were analyzed for: (**a**) liver non-heme Fe levels by bathophenanthroline-based assay; (**b**) liver *Hamp* RNA levels; (**c-e**) RBC counts (c), hemoglobin levels (d), and hematocrits (e) by complete blood counts; (**f, g**) liver copper (Cu) (f) and zinc (Zn) (g) by ICP-OES. Data are represented as means +/- standard deviation, with at least four animals per group. Data were tested for normal distribution by Shapiro-Wilk test; if not normally distributed, data were log transformed. Within each sex, groups were compared using two-way ANOVA with Tukey’s multiple comparisons test. (* P<0.05, ** P<0.01, *** P<0.001, **** P<0.0001)

**Fig. S25: Intestinal Fpn deficiency, initiated at P14, modestly impacts whole brain gene expression in *Slc30a10^-/-^* mice.** (**a-c**) Bulk RNA-seq was performed on whole brain samples from three female *Slc30a10^-/-^ Fpn^fl/fl^* and *Slc30a10^+/+^ Fpn^fl/fl^*, mice then assessed by similarity analysis (a), principal component analysis (b), and volcano plot (c) where genes more abundantly expressed in *Slc30a10^-/-^ Fpn^fl/fl^* mice have log_2_(fold change)>1. (**d-f**) Bulk RNA-seq was performed on whole brain samples from three female *Slc30a10^-/-^ Fpn^fl/fl^ Vil-ERT2* and *Slc30a10^-/-^ Fpn^fl/fl^* mice then assessed by similarity analysis (d), principal component analysis (e), and volcano plot (f) where genes more abundantly expressed in *Slc30a10^-/-^ Fpn^fl/fl^ Vil-ERT2* mice have log_2_(fold change)>1. In volcano plots, differentially expressed genes (adjusted P value<0.05 and absolute value of log_2_(fold change)>1) are shown as light orange points with gene names shown adjacent as space permitted; non-differentially expressed genes are shown as blue points.

**Fig. S26: Intestinal Fpn deficiency, initiated at P14, modestly impacts whole brain gene expression in *Slc30a10^-/-^* mice.** Venn analysis was performed on RNA-seq results from Fig. S25. Heatmaps are shown for 48 genes differentially expressed only in *Slc30a10^-/-^ Fpn^fl/fl^* vs. *Slc30a10^+/+^ Fpn^fl/fl^*, mice, 60 genes differentially expressed only in *Slc30a10^-/-^ Fpn^fl/fl^ Vil-ERT2* and *Slc30a10^-/-^ Fpn^fl/fl^*mice, and 17 genes differentially expressed in both comparisons.

**Fig. S27: Tamoxifen treatment of *Slc30a10^+/+^ Fpn^fl/fl^ Vil-ERT2* at P14 minimally impacts duodenum and liver *Slc39a14* and *Slc39a8* RNA levels.** *Slc30a10 Dmt1^fl/fl^* mice with and without *Vil-ERT2* were injected intraperitoneally with a five-day course of tamoxifen starting at P14. Some mice were injected subcutaneously with Fe dextran on P19. At P28, mice were analyzed for duodenum (**a**) and liver (**b**) *Slc39a14* and duodenum (**c**) and liver (**d**) *Slc39a8* RNA levels (relative to *Hprt1* RNA levels) by qRT-PCR. Data are represented as means +/- standard deviation, with at least four animals per group. Data were tested for normal distribution by Shapiro-Wilk test; if not normally distributed, data were log transformed. Within each sex, groups were compared using two-way ANOVA with Tukey’s multiple comparisons test. (* P<0.05, ** P<0.01, *** P<0.001, **** P<0.0001)

**Fig. S28: Treatment of *Slc30a10^-/-^* mice with *Tmprss6* ASOs attenuates Mn excess, increases body mass, and leads to hepatomegaly.** *Slc30a10^+/+^*mice were treated with sterile-filtered PBS and *Slc30a10^-/-^* mice were treated with sterile-filtered GalNAc-control or *Tmprss6* ASO (2.5 µg/g). ASOs were administered by intraperitoneal injection twice a week beginning at weaning until six weeks of age for a total of 12 doses. Animals were harvested two days after the last dose then analyzed for: (**a, b**) liver *Tmprss6* (a) and *Hamp* (b) RNA levels by qRT-PCR; (**c**) tissue Mn levels by ICP-OES; (**d, e**) body mass (throughout the treatment course); (**f**) liver body mass. Data are represented as means +/- standard deviation, with at least four animals per group. Data were tested for normal distribution by Shapiro-Wilk test; if not normally distributed, data were log transformed. Within each sex, groups were compared using two-way ANOVA with Tukey’s multiple comparisons test; in (d, e), groups were compared at each age. (* P<0.05, ** P<0.01, *** P<0.001, **** P<0.0001)

**Fig. S29: Intestinal deficiency in Dmt1 or Fpn minimally impacts Mn levels in mice raised on Mn-rich diets.** *Dmt1^fl/fl^* (**a-d, i-l**) or *Fpn^fl/fl^* (**e-h, m-p**) mice with and without *Vil-ERT2* were injected intraperitoneally with a five-day course of tamoxifen starting at P21, then weaned onto diets containing 100 or 2400 ppm Mn. At eight weeks of age, mice were analyzed by ICP-OES for Fe levels in liver (**a, e**), bone (**b, f**), brain (**c, g**), and pancreas (**d, h**) and for Mn levels in liver (**i, m**), bone (**j, n**), brain (**k, o**), and pancreas (**l, p**). Data are represented as means +/- standard deviation, with at least four animals per group. Data were tested for normal distribution by Shapiro-Wilk test; if not normally distributed, data were log transformed. Within each sex, groups were compared using two-way ANOVA with Tukey’s multiple comparisons test. (* P<0.05, ** P<0.01, *** P<0.001, **** P<0.0001)

**Fig. S30: Whole-body Slc30a10 deficiency leads to Mn excess in *Hjv^-/-^* mice.** Five-week-old *Slc30a10 Hjv* mice were characterized for: (**a,b**) liver *Slc30a10* (a) and *Hjv* (b) RNA levels, relative to *Hprt1* RNA levels, by qPCR; (**c-g**) Mn levels in liver (c), brain (d), bone (e), pancreas (f), and small intestine (g) by ICP-OES; (**h**) blood Mn levels by GFAAS; (**i-k**) RBC counts (i), hemoglobin levels (j), and hematocrit (k) by complete blood count; (**l**) liver *Epo* RNA levels by qPCR. Data are represented as means +/- standard deviation, with at least five animals per group. Data were tested for normal distribution by Shapiro-Wilk test; if not normally distributed, data were log transformed. Within each sex, groups were compared using two-way ANOVA with Tukey’s multiple comparisons test. (* P<0.05, ** P<0.01, *** P<0.001, **** P<0.0001)

**Fig. S31: Laser ablation inductively coupled mass spectrometry of *Slc30a10 Hjv* small intestine.** Laser ablation inductively coupled mass spectrometry was performed on small intestine sections from five-week-old *Slc30a10 Hjv* mice. Shown are images for Mn, Fe, copper (Cu), and zinc (Zn). Scale bar in top left image applies to all images.

**Fig. S32: X-ray fluorescence microscopy of *Slc30a10 Hjv* small intestine.** X-ray fluorescence microscopy was performed on small intestine sections from five-week-old *Slc30a10 Hjv* mice. Shown are images for phosphorus (P), sulfur (S), potassium (K), Mn, Fe, copper (Cu), and zinc (Zn).

**Fig. S33: *Slc30a10^-/-^ Hjv^-/-^* mice have Fe excess.** Five-week-old *Slc30a10 Hjv* mice were characterized for: (**a-c**) non-heme Fe levels in liver (a), pancreas (b), and spleen (c) by bathophenanthroline-based assay; (**d-i**) liver Fe levels and distribution by diaminobenzidine-enhanced Fe stains. Note that two images are shown each for *Slc30a10^-/-^ Hjv^+/+^* (f,g) and *Slc30a10^-/-^ Hjv^-/-^* (h,i) to show variability within these genotypes. For (a-c), data are represented as means +/- standard deviation, with at least five animals per group. Data were tested for normal distribution by Shapiro-Wilk test; if not normally distributed, data were log transformed. Within each sex, groups were compared using two-way ANOVA with Tukey’s multiple comparisons test. (* P<0.05, ** P<0.01, *** P<0.001, **** P<0.0001)

**Fig. S34: *Slc30a10^-/-^ Hjv^-/-^* mice have increased Mn absorption.** Five-week-old *Slc30a10 Hjv* mice were characterized for: (**a**) liver hepcidin (*Hamp*) RNA levels by qPCR; (**b**) ^59^Fe absorption by gastric gavage; (**c, d**) bile flow rates (c) and Mn levels (d) by surgical bile collection and GFAAS respectively; (**e**) ^52^Mn absorption by gastric gavage. Data are represented as means +/- standard deviation, with at least five animals per group. Data were tested for normal distribution by Shapiro-Wilk test; if not normally distributed, data were log transformed. Within each sex, groups were compared using two-way ANOVA with Tukey’s multiple comparisons test. (* P<0.05, ** P<0.01, *** P<0.001, **** P<0.0001)

**Fig. S35: Intestinal Slc30a10 deficiency does not impact Fe levels in *Hjv^-/-^* mice.** Eight-week-old *Hjv Slc30a10^fl/fl^*+/- *Vil* were characterized for: (**a-c**) RBC counts (a), hemoglobin levels (b), and hematocrits (c) by complete blood counts; (**d-f**) non-heme Fe levels in liver (d), pancreas (e), and spleen (f) by bathophenanthroline-based assay; (**g**) diaminobenzidine-enhanced Fe stains of livers at 20x magnification with 20 μm scale bar. For (a-f), data are represented as means +/- standard deviation, with at least five animals per group. Data were tested for normal distribution by Shapiro-Wilk test; if not normally distributed, data were log transformed. Within each sex, groups were compared using two-way ANOVA with Tukey’s multiple comparisons test. (* P<0.05, ** P<0.01, *** P<0.001, **** P<0.0001)

## References

1. Baj, J. et al. Consequences of Disturbing Manganese Homeostasis. Int J Mol Sci 24, 14959 (2023).

2. Jomova, K. et al. Essential metals in health and disease. Chem Biol Interact 367, 110173 (2022).

3. Tuschl, K. et al. Syndrome of hepatic cirrhosis, dystonia, polycythemia, and hypermanganesemia caused by mutations in SLC30A10, a manganese transporter in man. Am. J. Hum. Genet. 90, 457–466 (2012).

4. Quadri, M. et al. Mutations in SLC30A10 cause parkinsonism and dystonia with hypermanganesemia, polycythemia, and chronic liver disease. Am. J. Hum. Genet. 90, 467–477 (2012).

5. Mercadante, C. J. et al. Manganese transporter Slc30a10 controls physiological manganese excretion and toxicity. J. Clin. Invest. 129, 5442–5461 (2019).

6. Hutchens, S. et al. Deficiency in the manganese efflux transporter SLC30A10 induces severe hypothyroidism in mice. J. Biol. Chem. 292, 9760–9773 (2017).

7. Taylor, C. A. et al. SLC30A10 transporter in the digestive system regulates brain manganese under basal conditions while brain SLC30A10 protects against neurotoxicity. J. Biol. Chem. 294, 1860–1876 (2019).

8. Prajapati, M. et al. Hepatic HIF2 is a key determinant of manganese excess and polycythemia in SLC30A10 deficiency. JCI Insight e169738 (2024) doi:10.1172/jci.insight.169738.

9. Galy, B., Conrad, M. & Muckenthaler, M. Mechanisms controlling cellular and systemic iron homeostasis. Nat Rev Mol Cell Biol 25, 133–155 (2024).

10. Adams, P. C., Jeffrey, G. & Ryan, J. Haemochromatosis. Lancet 401, 1811–1821 (2023).

11. Wolff, N. A. et al. A role for divalent metal transporter (DMT1) in mitochondrial uptake of iron and manganese. Sci Rep 8, 211 (2018).

12. Tai, Y. K., Chew, K. C. M., Tan, B. W. Q., Lim, K.-L. & Soong, T. W. Iron mitigates DMT1-mediated manganese cytotoxicity via the ASK1-JNK signaling axis: Implications of iron supplementation for manganese toxicity. Sci Rep 6, 21113 (2016).

13. Illing, A. C., Shawki, A., Cunningham, C. L. & Mackenzie, B. Substrate profile and metal-ion selectivity of human divalent metal-ion transporter-1. J Biol Chem 287, 30485–30496 (2012).

14. Fujishiro, H., Yano, Y., Takada, Y., Tanihara, M. & Himeno, S. Roles of ZIP8, ZIP14, and DMT1 in transport of cadmium and manganese in mouse kidney proximal tubule cells. Metallomics 4, 700–708 (2012).

15. Wang, D., Song, Y., Li, J., Wang, C. & Li, F. Structure and metal ion binding of the first transmembrane domain of DMT1. Biochim Biophys Acta 1808, 1639–1644 (2011).

16. Garrick, M. D. et al. Comparison of mammalian cell lines expressing distinct isoforms of divalent metal transporter 1 in a tetracycline-regulated fashion. Biochem. J. 398, 539–546 (2006).

17. Thompson, K. et al. Olfactory uptake of manganese requires DMT1 and is enhanced by anemia. FASEB J. 21, 223–230 (2007).

18. Knöpfel, M., Zhao, L. & Garrick, M. D. Transport of divalent transition-metal ions is lost in small-intestinal tissue of b/b Belgrade rats. Biochemistry 44, 3454–3465 (2005).

19. Shawki, A. et al. Intestinal DMT1 is critical for iron absorption in the mouse but is not required for the absorption of copper or manganese. Am. J. Physiol. Gastrointest. Liver Physiol. 309, G635–G647 (2015).

20. Kim, J., Buckett, P. D. & Wessling-Resnick, M. Absorption of manganese and iron in a mouse model of hemochromatosis. PLoS ONE 8, e64944 (2013).

21. Seo, Y. A., Elkhader, J. A. & Wessling-Resnick, M. Distribution of manganese and other biometals in flatiron mice. Biometals 29, 147–155 (2016).

22. Seo, Y. A. & Wessling-Resnick, M. Ferroportin deficiency impairs manganese metabolism in flatiron mice. FASEB J. 29, 2726–2733 (2015).

23. Choi, E.-K., Nguyen, T.-T., Iwase, S. & Seo, Y. A. Ferroportin disease mutations influence manganese accumulation and cytotoxicity. FASEB J. 33, 2228–2240 (2019).

24. Madejczyk, M. S. & Ballatori, N. The iron transporter ferroportin can also function as a manganese exporter. Biochim. Biophys. Acta 1818, 651–657 (2012).

25. Yin, Z. et al. Ferroportin is a manganese-responsive protein that decreases manganese cytotoxicity and accumulation. J. Neurochem. 112, 1190–1198 (2010).

26. Cr, A. et al. Comparative analysis of the functional properties of human and mouse ferroportin. American journal of physiology. Cell physiology 324, (2023).

27. Jin, L. et al. Mice overexpressing hepcidin suggest ferroportin does not play a major role in Mn homeostasis. Metallomics 11, 959–967 (2019).

28. Prajapati, M. et al. AAV-mediated hepatic expression of SLC30A10 and the Thr95Ile variant attenuates manganese excess and other phenotypes in Slc30a10-deficient mice. J Biol Chem 105732 (2024) doi:10.1016/j.jbc.2024.105732.

29. Liu, C., Jursa, T., Aschner, M., Smith, D. R. & Mukhopadhyay, S. Up-regulation of the manganese transporter SLC30A10 by hypoxia-inducible factors defines a homeostatic response to manganese toxicity. Proc Natl Acad Sci U S A 118, e2107673118 (2021).

30. Anagianni, S. & Tuschl, K. Genetic Disorders of Manganese Metabolism. Curr Neurol Neurosci Rep 19, 33 (2019).

31. Choi, E.-K. et al. The manganese transporter SLC39A8 links alkaline ceramidase 1 to inflammatory bowel disease. Nat Commun 15, 4775 (2024).

32. Fisher, A. L. et al. Endothelial ZIP8 plays a minor role in BMP6 regulation by iron in mice. Blood blood.2023023385 (2024) doi:10.1182/blood.2023023385.

33. Zhang, V. et al. A mouse model characterizes the roles of ZIP8 in systemic iron recycling and lung inflammation and infection. Blood Adv 7, 1336–1349 (2023).

34. Pasquadibisceglie, A., Bonaccorsi di Patti, M. C., Musci, G. & Polticelli, F. Membrane Transporters Involved in Iron Trafficking: Physiological and Pathological Aspects. Biomolecules 13, 1172 (2023).

35. Camaschella, C., Nai, A. & Silvestri, L. Iron metabolism and iron disorders revisited in the hepcidin era. Haematologica 105, 260–272 (2020).

36. Silvestri, L. et al. Managing the Dual Nature of Iron to Preserve Health. Int J Mol Sci 24, 3995 (2023).

37. Coppola, T. et al. Case Report: Childhood Erythrocytosis due to Hypermanganesemia Caused by Homozygous SLC30A10 Mutation. Front. Hematol. 3, (2024).

38. Bjørklund, G., Dadar, M., Peana, M., Rahaman, M. S. & Aaseth, J. Interactions between iron and manganese in neurotoxicity. Arch Toxicol 94, 725–734 (2020).

39. Carmona, A. et al. SLC30A10 Mutation Involved in Parkinsonism Results in Manganese Accumulation within Nanovesicles of the Golgi Apparatus. ACS Chem Neurosci 10, 599–609 (2019).

40. Fujishiro, H. & Kambe, T. Manganese transport in mammals by zinc transporter family proteins, ZNT and ZIP. J Pharmacol Sci 148, 125–133 (2022).

41. Wahlberg, K. et al. Common Polymorphisms in the Solute Carrier SLC30A10 are Associated With Blood Manganese and Neurological Function. Toxicol Sci 149, 473–483 (2016).

42. Ng, E. et al. Genome-wide association study of toxic metals and trace elements reveals novel associations. Hum. Mol. Genet. 24, 4739–4745 (2015).

43. Wahlberg, K. E. et al. Polymorphisms in Manganese Transporters SLC30A10 and SLC39A8 Are Associated With Children’s Neurodevelopment by Influencing Manganese Homeostasis. Front Genet 9, 664 (2018).

44. Wahlberg, K. et al. Polymorphisms in manganese transporters show developmental stage and sex specific associations with manganese concentrations in primary teeth. Neurotoxicology 64, 103–109 (2018).

45. Sohal, A. & Kowdley, K. V. A Review of New Concepts in Iron Overload. Gastroenterol Hepatol (N Y) 20, 98–107 (2024).

46. Pasricha, S.-R., Tye-Din, J., Muckenthaler, M. U. & Swinkels, D. W. Iron deficiency. Lancet 397, 233– 248 (2021).

47. Bjørklund, G., Dadar, M., Peana, M., Rahaman, M. S. & Aaseth, J. Interactions between iron and manganese in neurotoxicity. Arch Toxicol 94, 725–734 (2020).

48. Lu, K. et al. Association between serum iron, blood lead, cadmium, mercury, selenium, manganese and low cognitive performance in old adults from National Health and Nutrition Examination Survey (NHANES): a cross-sectional study. Br J Nutr 130, 1743–1753 (2023).

49. Larvie, D. Y., Erikson, K. M. & Armah, S. M. Elevated whole blood manganese is associated with impaired cognition in older adults, NHANES 2013-2014 cycle. Neurotoxicology 91, 94–99 (2022).

50. Ge, S. X., Jung, D. & Yao, R. ShinyGO: a graphical gene-set enrichment tool for animals and plants. Bioinformatics 36, 2628–2629 (2020).

51. Vogt, S. MAPSL: A set of software tools for analysis and visualization of 3D X-ray fluorescence data sets. Journal de Physique IV 104, 635–638 (2003).

