## Supplemental Figures for "Manganese transporter SLC30A10 and iron transporters SLC40A1 and SLC11A2 impact dietary manganese absorption"

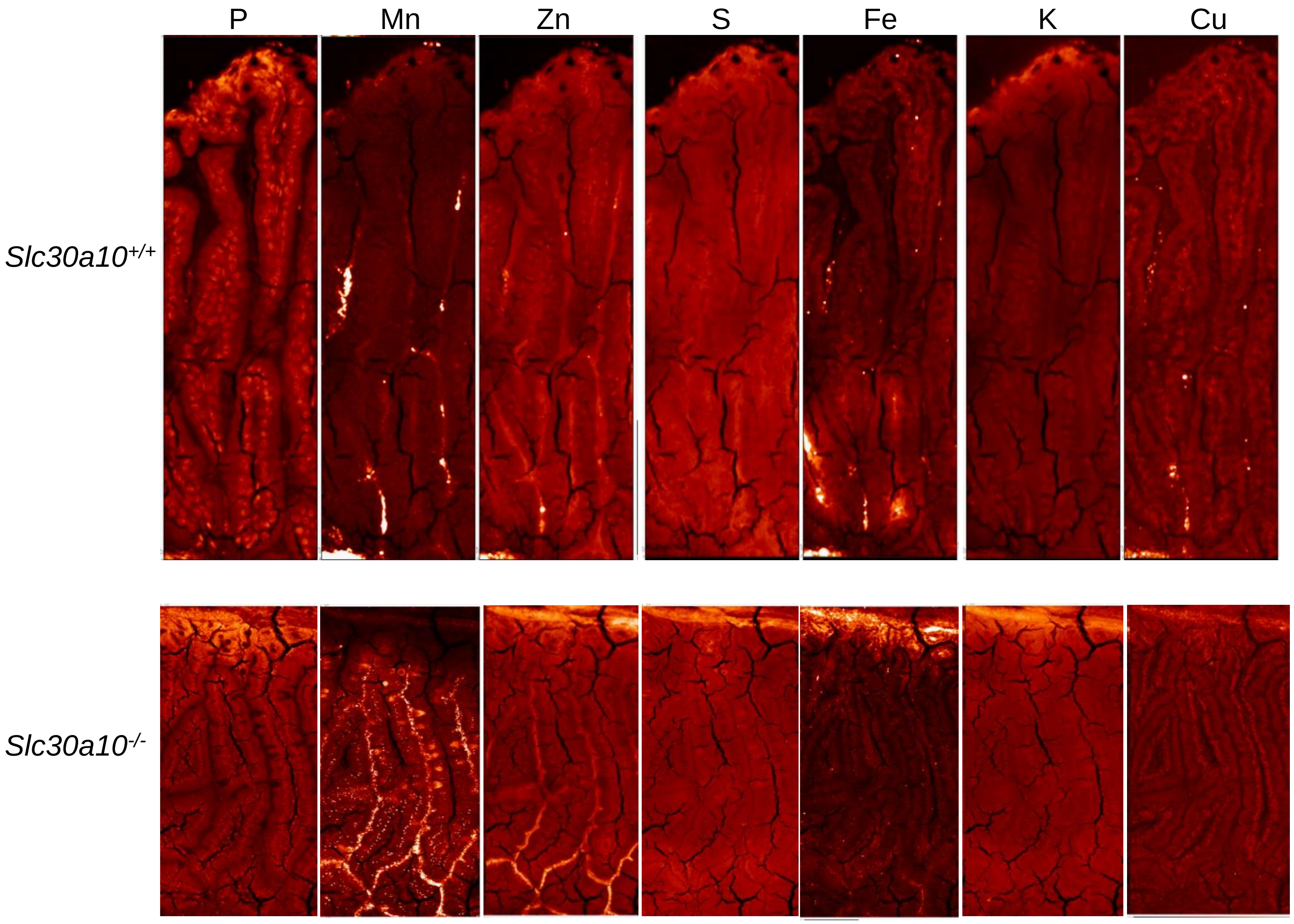


Figure S1

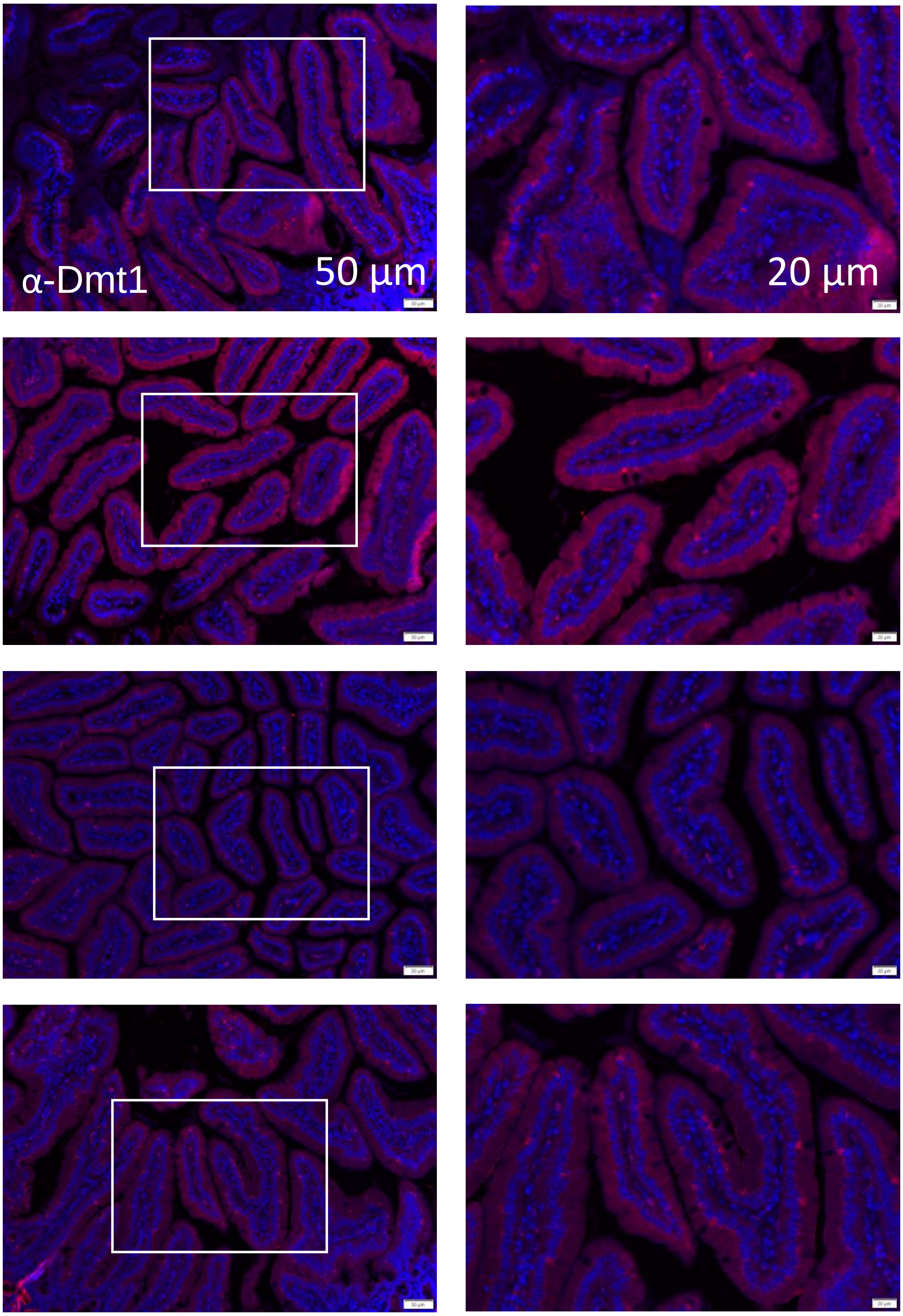


Female *Slc30a10*$^{+/+}$

Figure S2

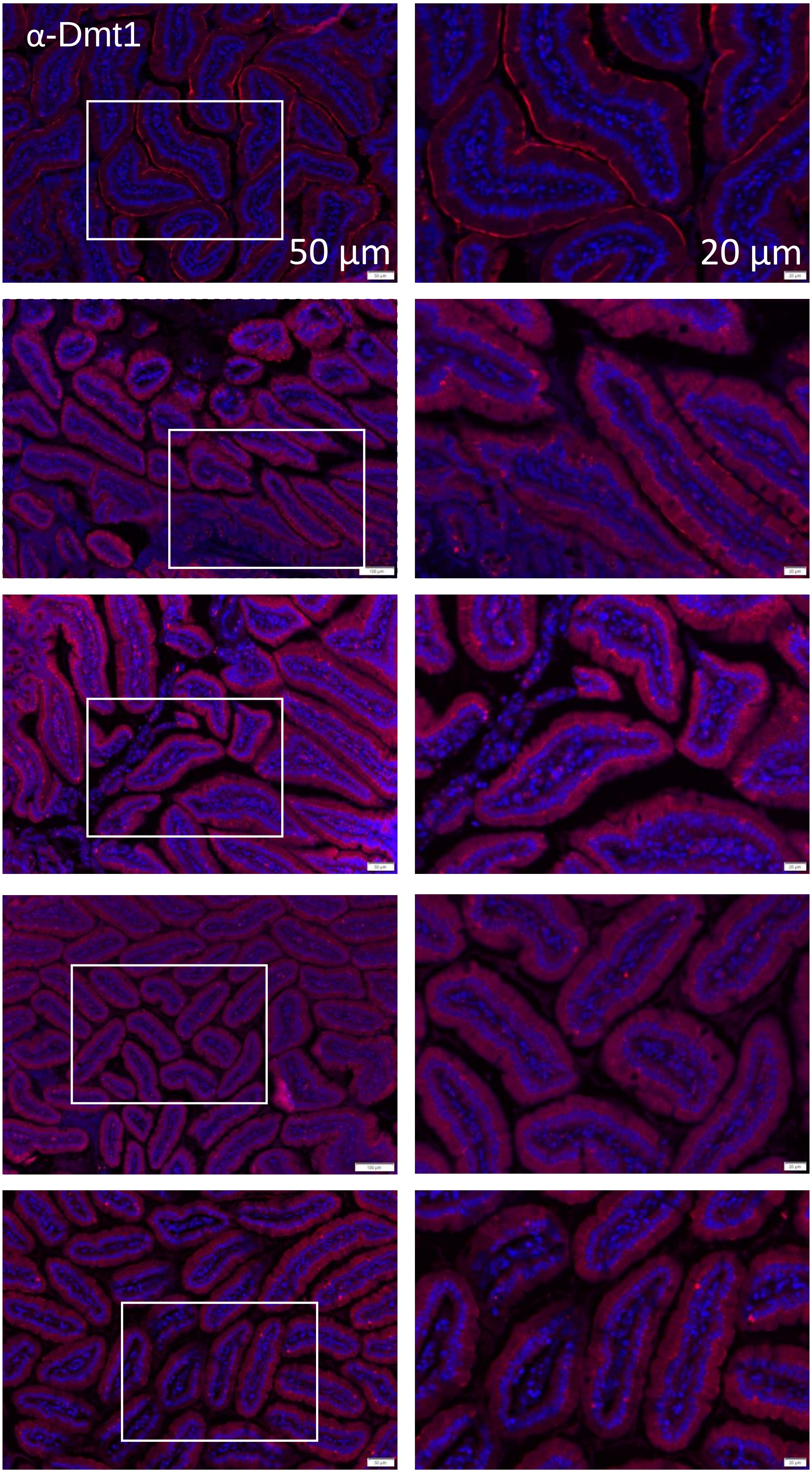


Male *Slc30a10*$^{+/+}$

Figure S3

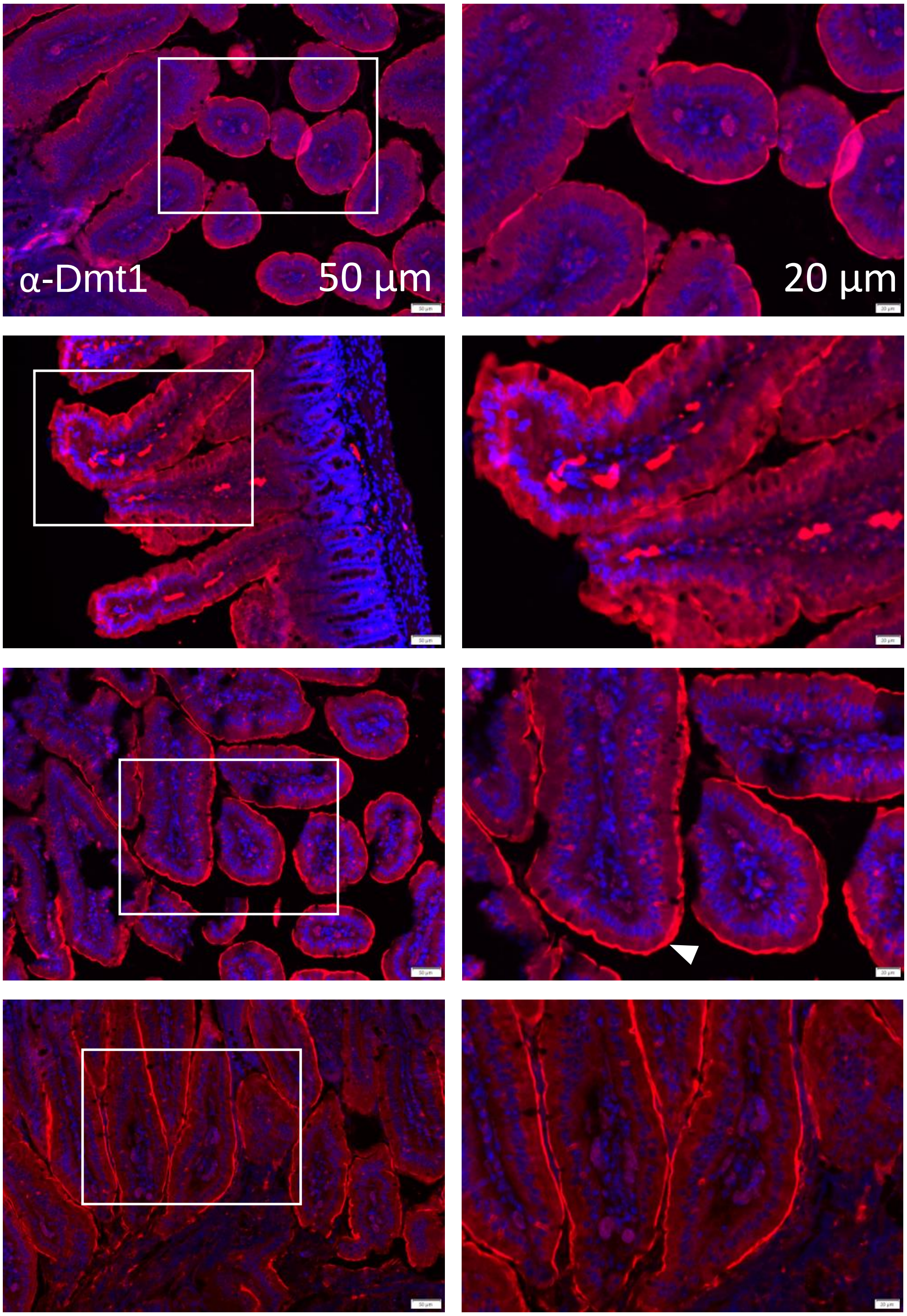


Female *Slc30a10*$^{-/-}$

Figure S4

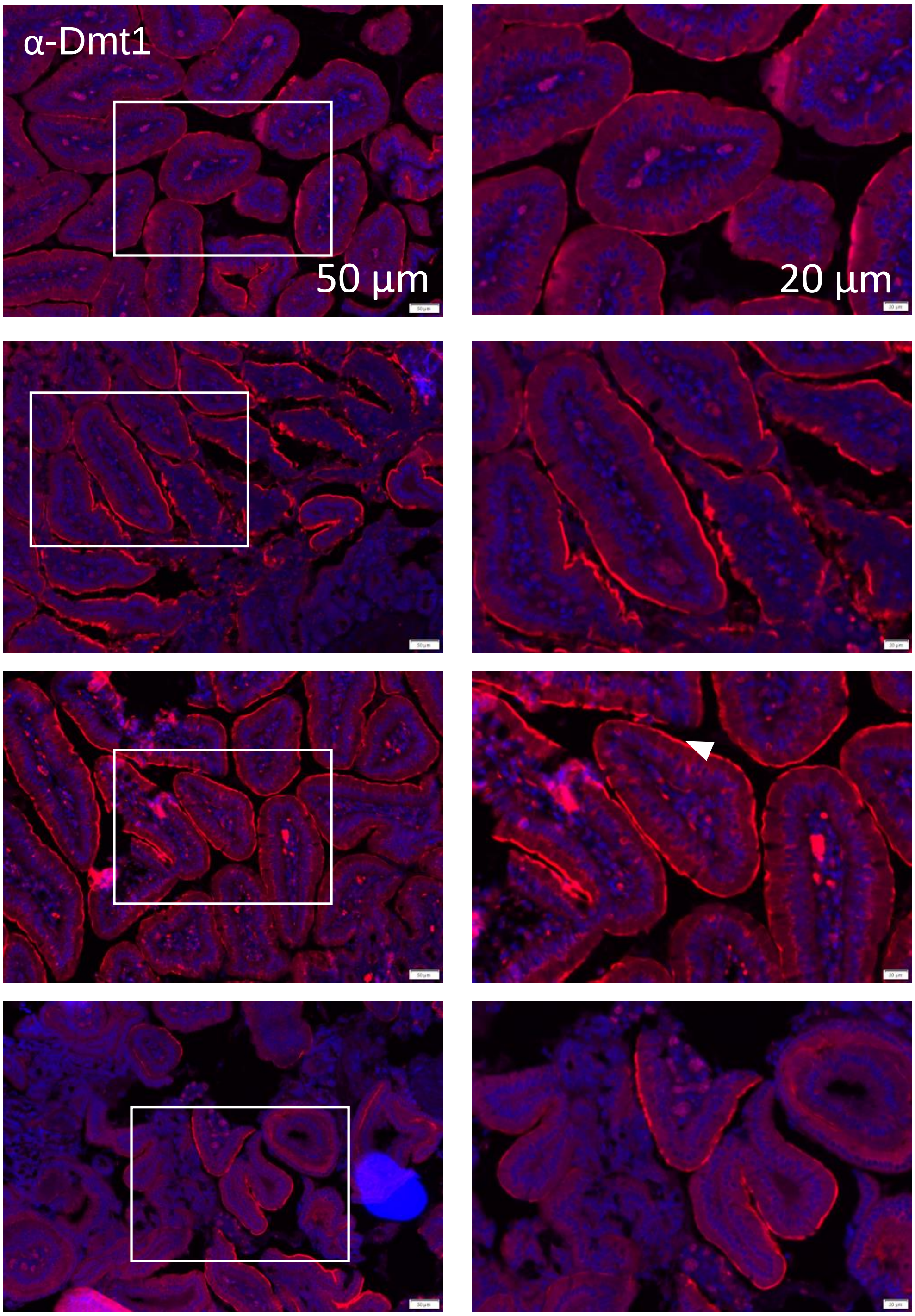


Male *Slc30a10*$^{-/-}$

Figure S5

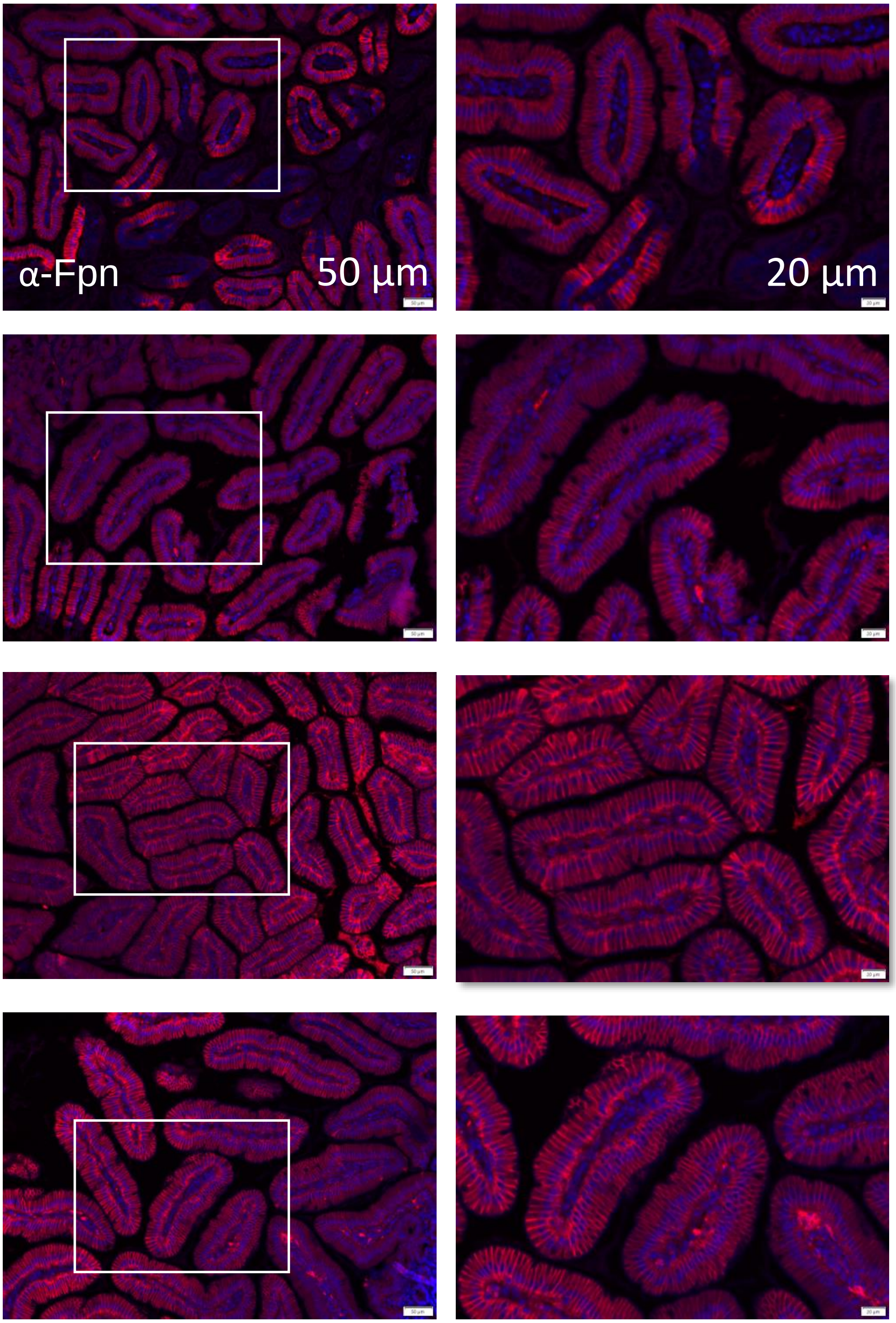


Female *Slc30a10*$^{+/+}$

Figure S6

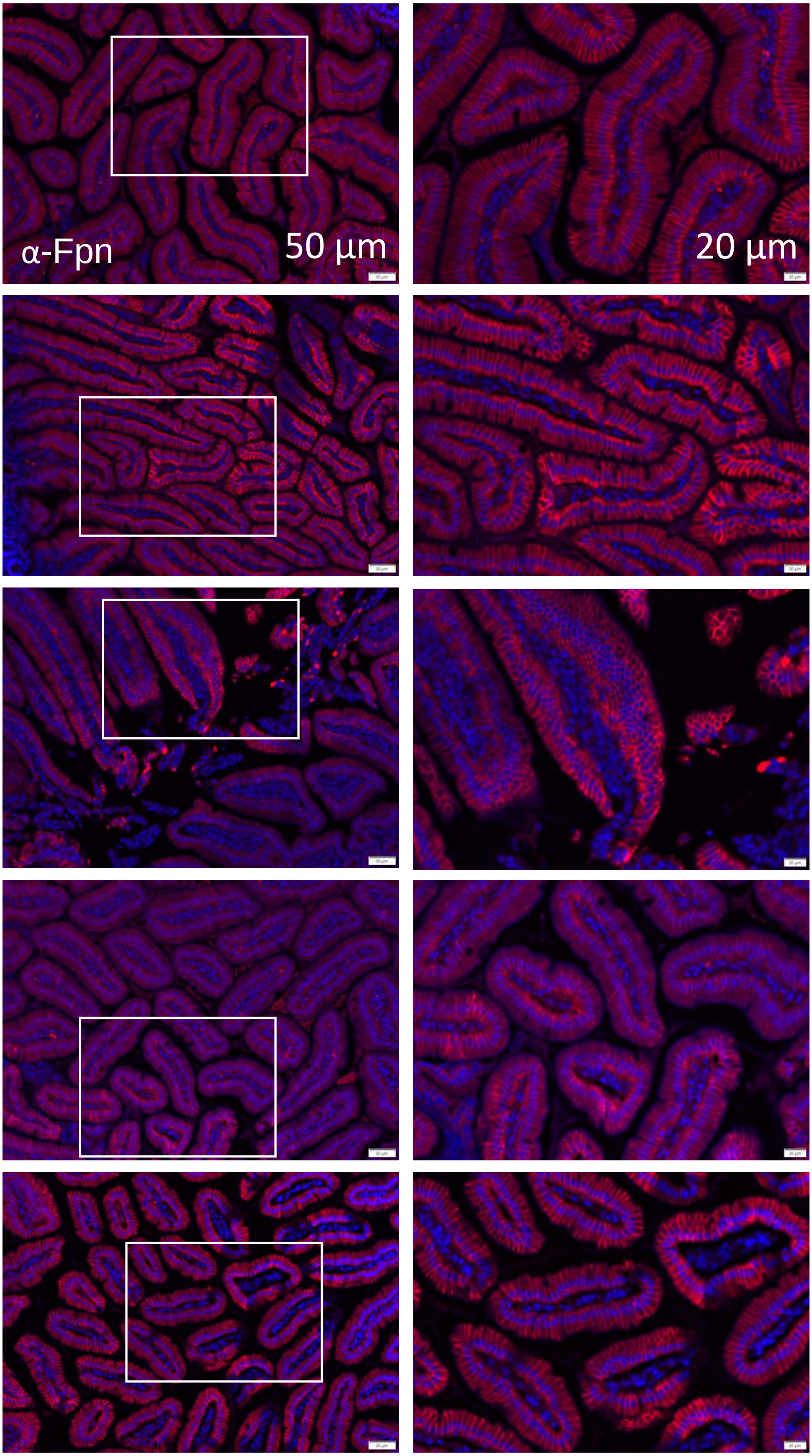

Male *Slc30a10*$^{+/+}$

Figure S7

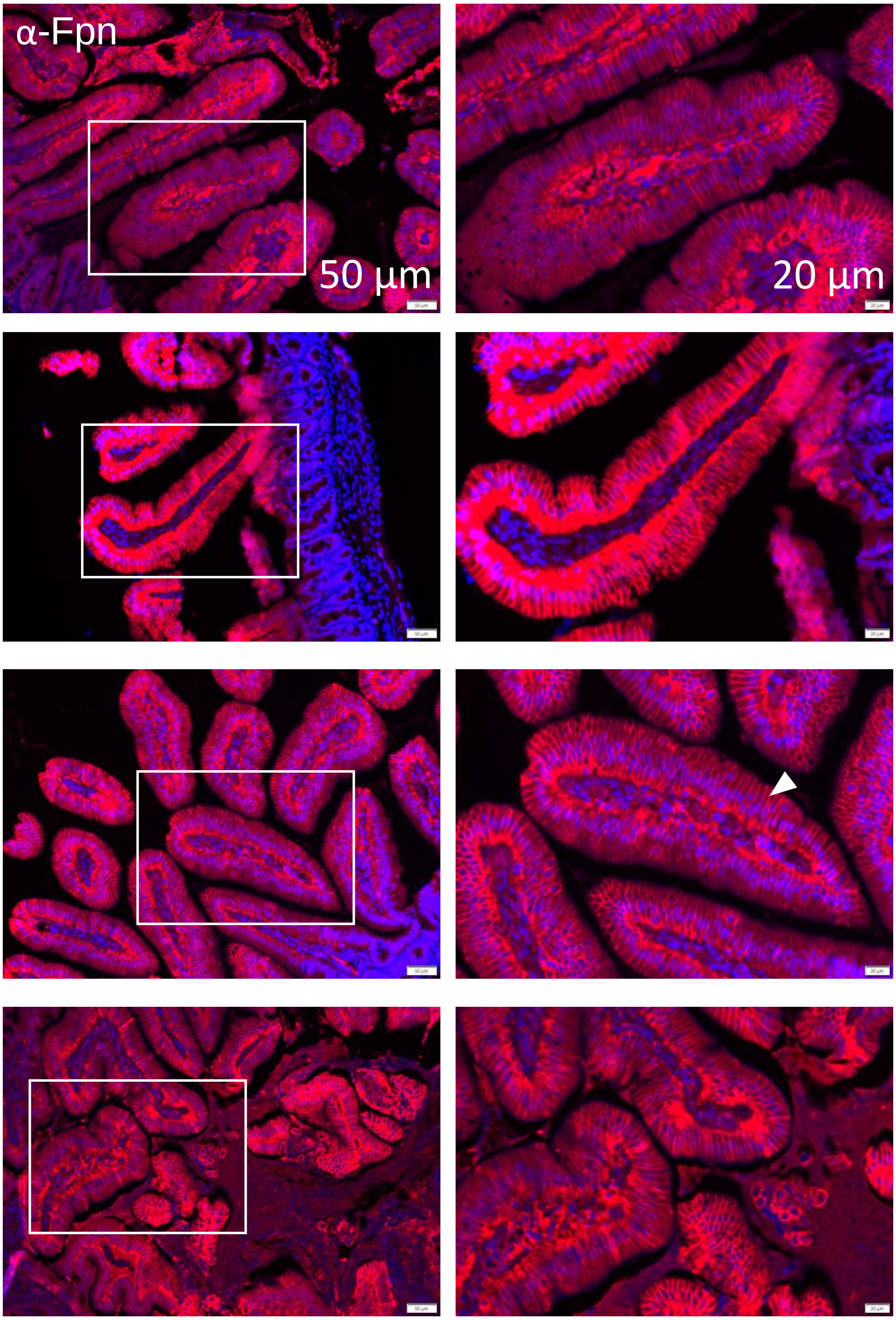

Female *Slc30a10*$^{-/-}$

Figure S8

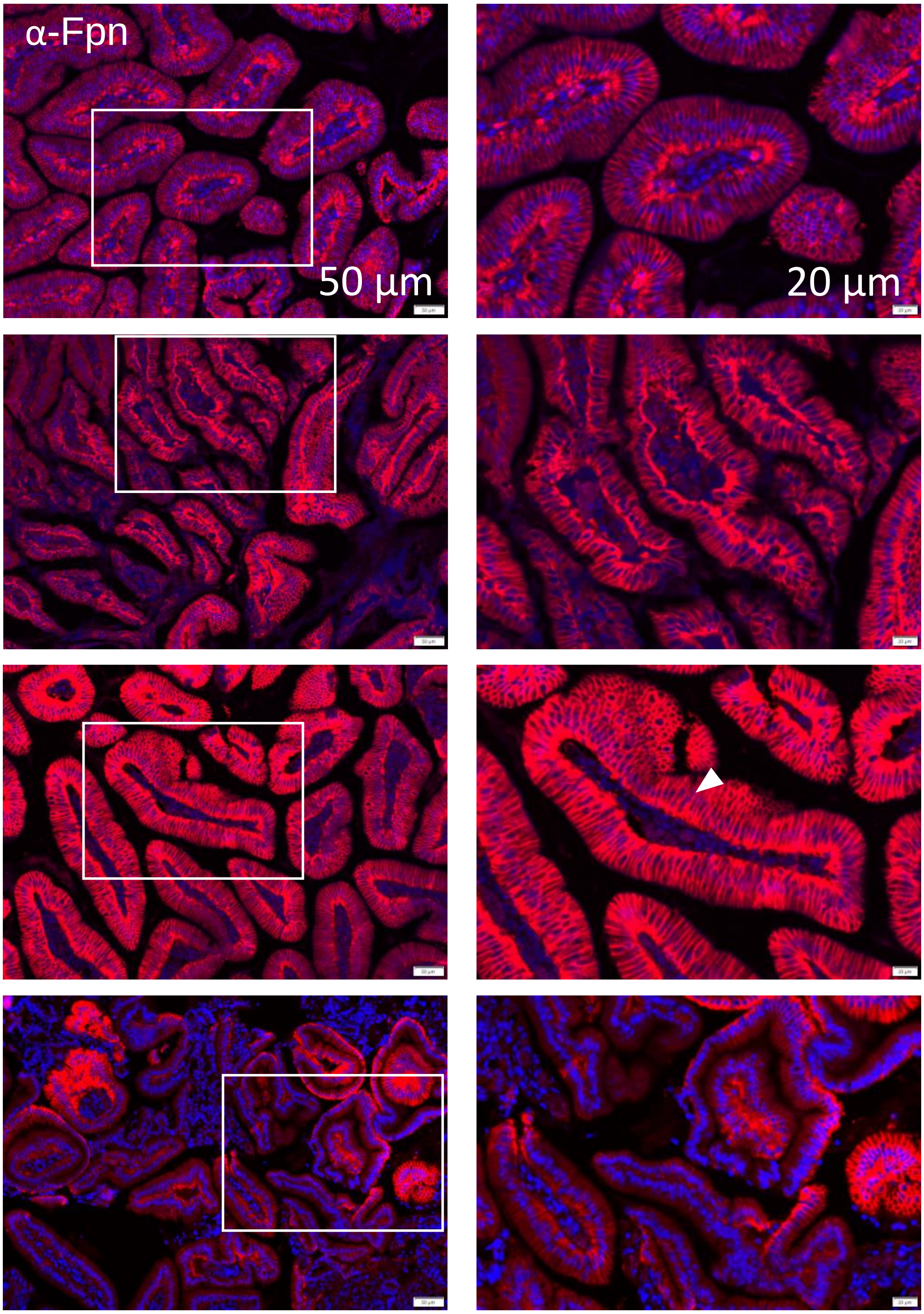


Male *Slc30a10*$^{-/-}$

Figure S9

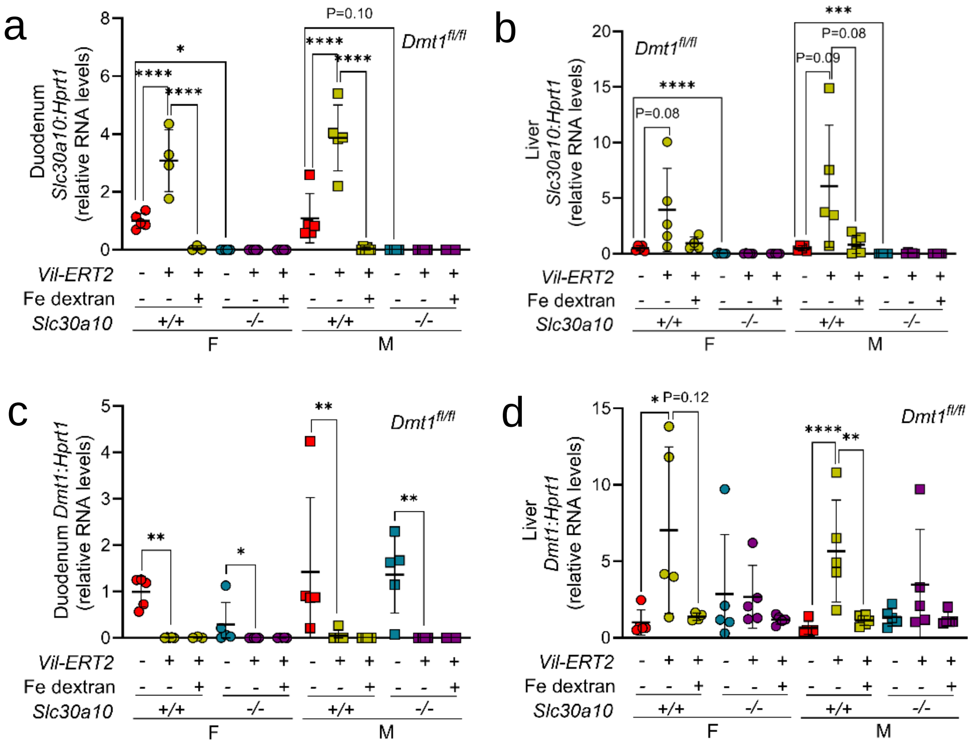
a
Duodenum Slc30a10:Hprt1 (relative RNA levels)
Dmt1fl/fl
P=0.10
Vil-ERT2
Fe dextran
Slc30a10
+/+
-/-
F
M
b
Liver Slc30a10:Hprt1 (relative RNA levels)
P=0.08
P=0.09
c
Duodenum Dmt1:Hprt1 (relative RNA levels)
d
Liver Dmt1:Hprt1 (relative RNA levels)
P=0.12

Figure S10

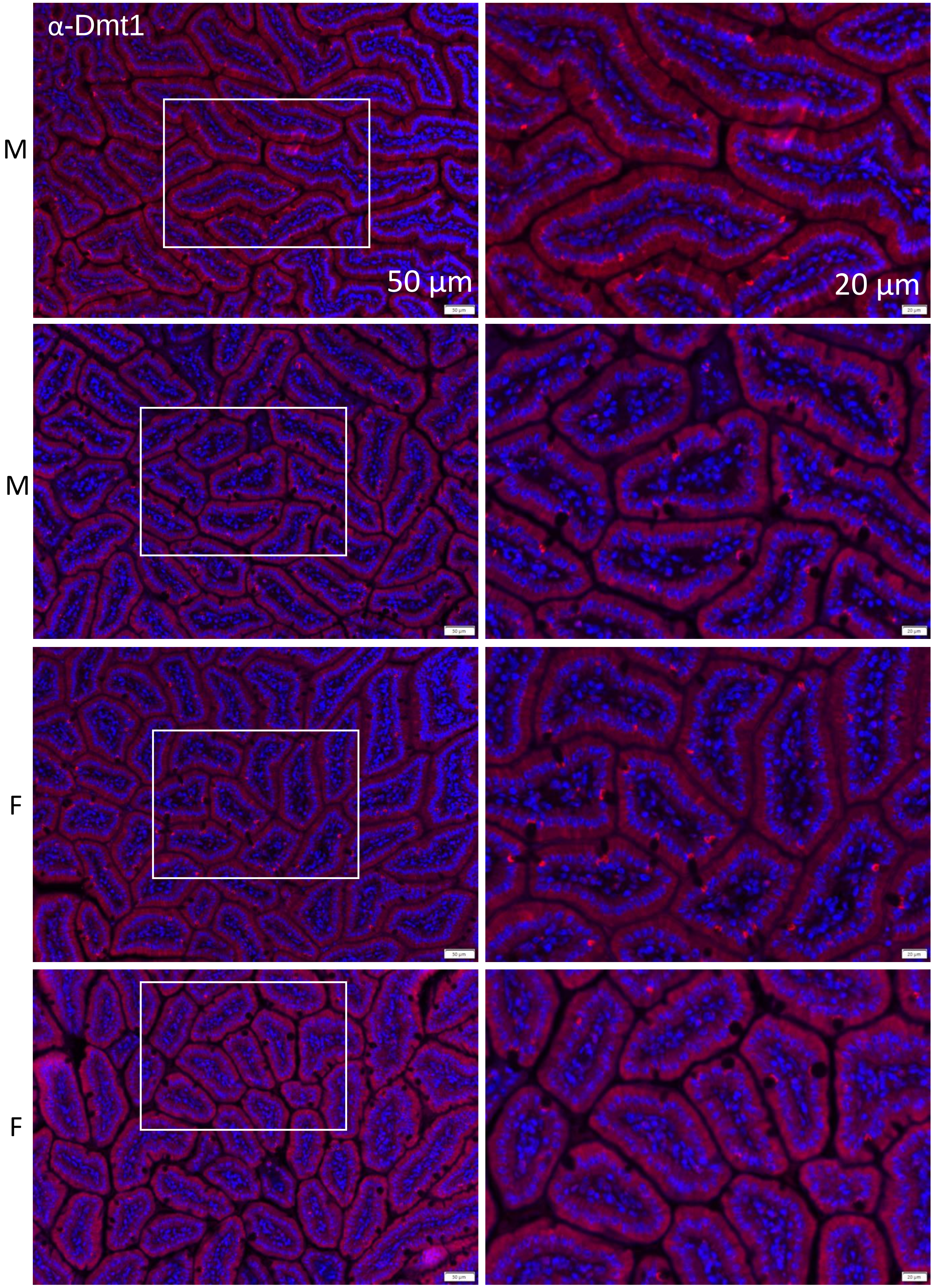


*Slc30a10*$^{+/+}$ *Dmt1*$^{fl/fl}$

Figure S11

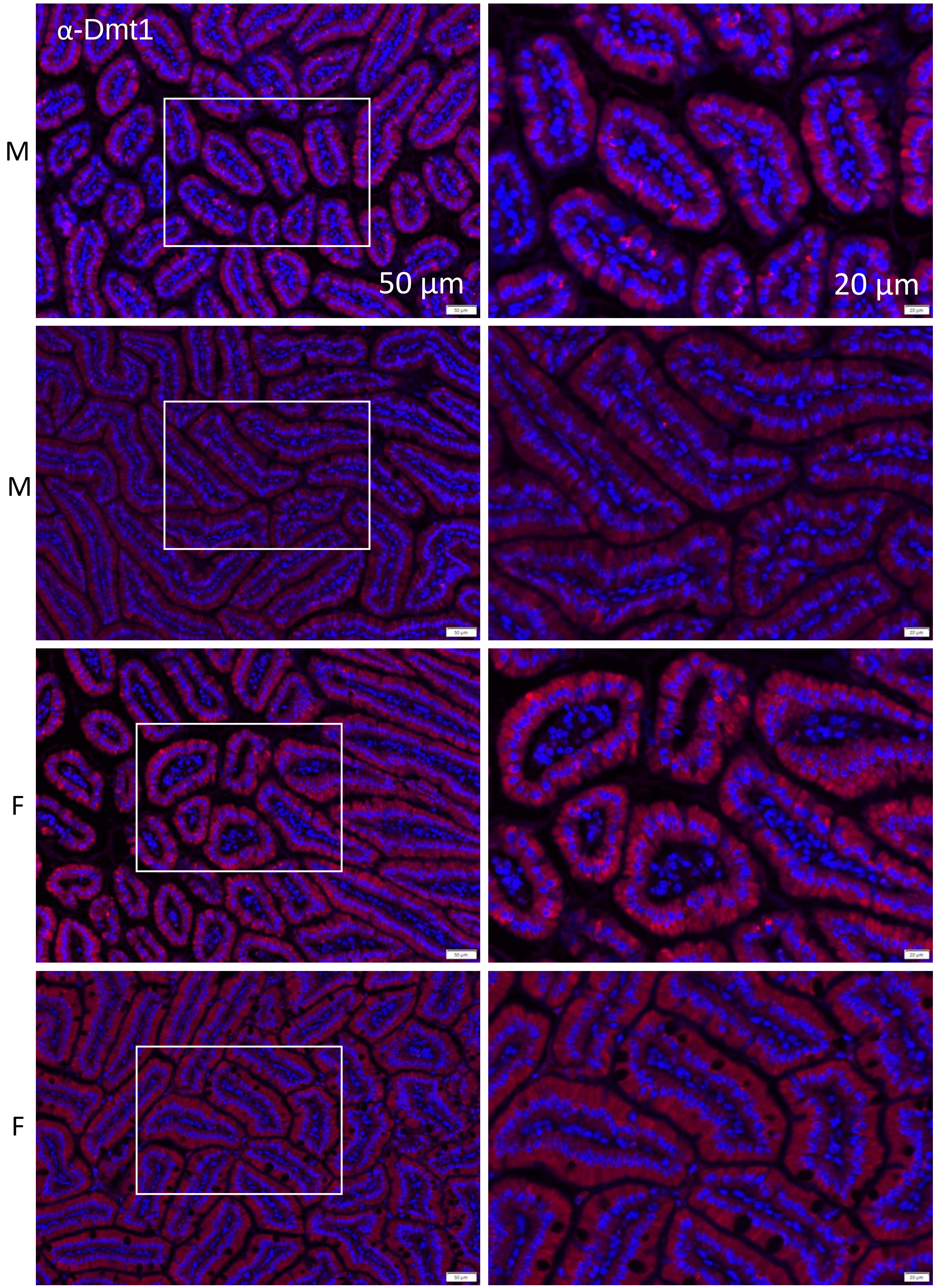


*Slc30a10*$^{+/+}$ *Dmt1*$^{fl/fl}$ *Vil-ERT2*

Figure S12

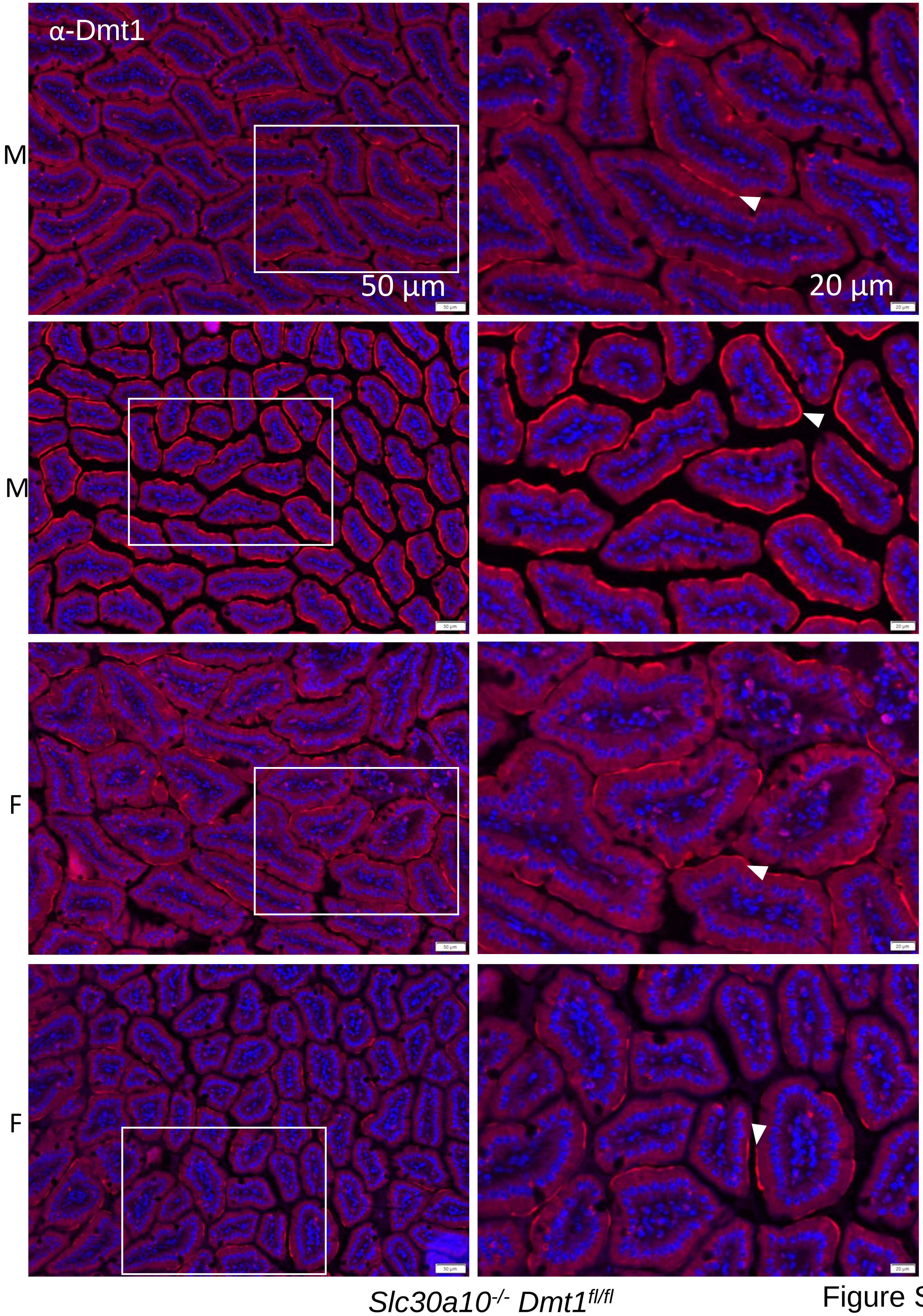


Figure S13

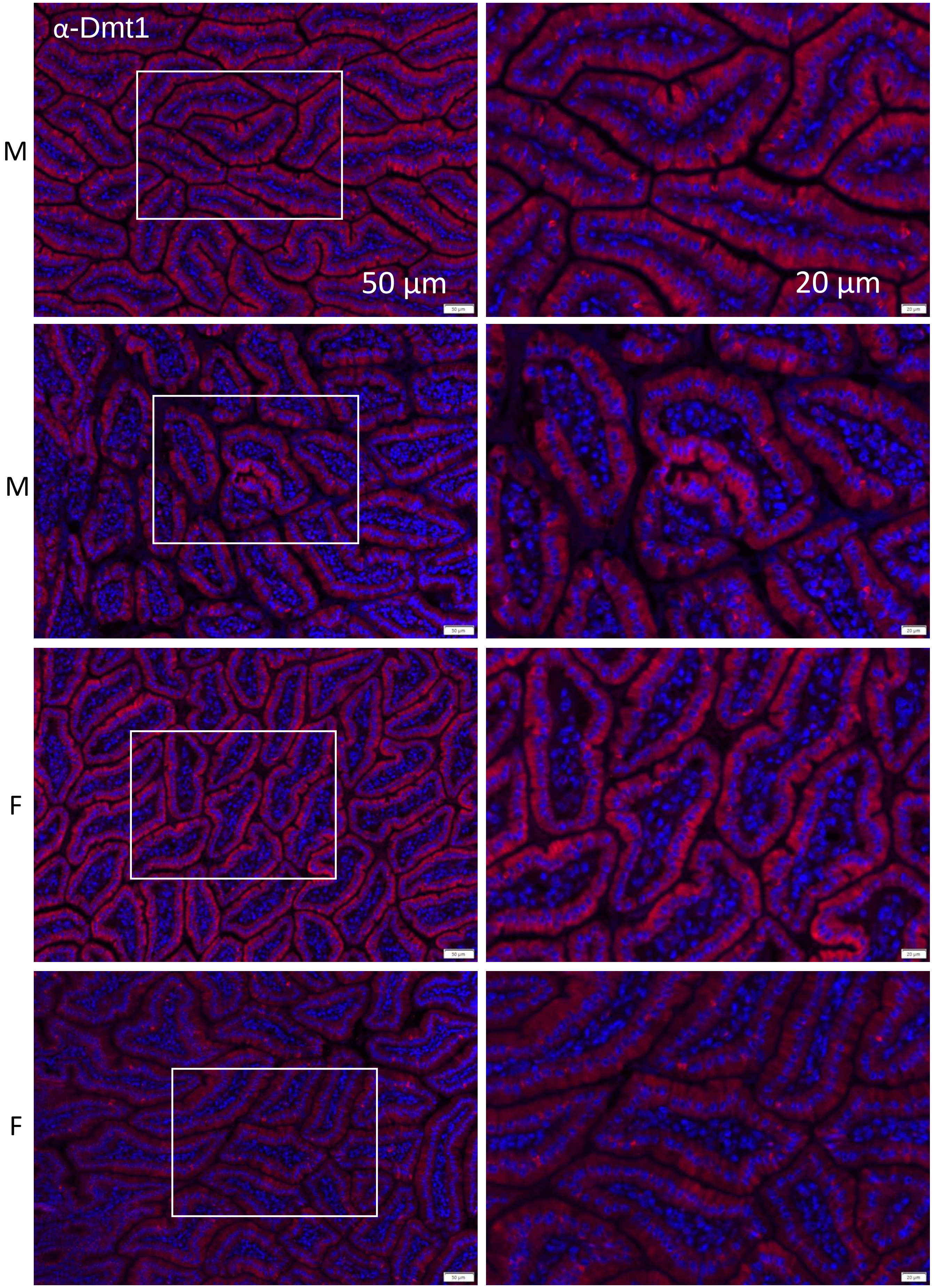


*Slc30a10*$^{-/-}$ *Dmt1*$^{fl/fl}$ *Vil-ERT2*

Figure S14

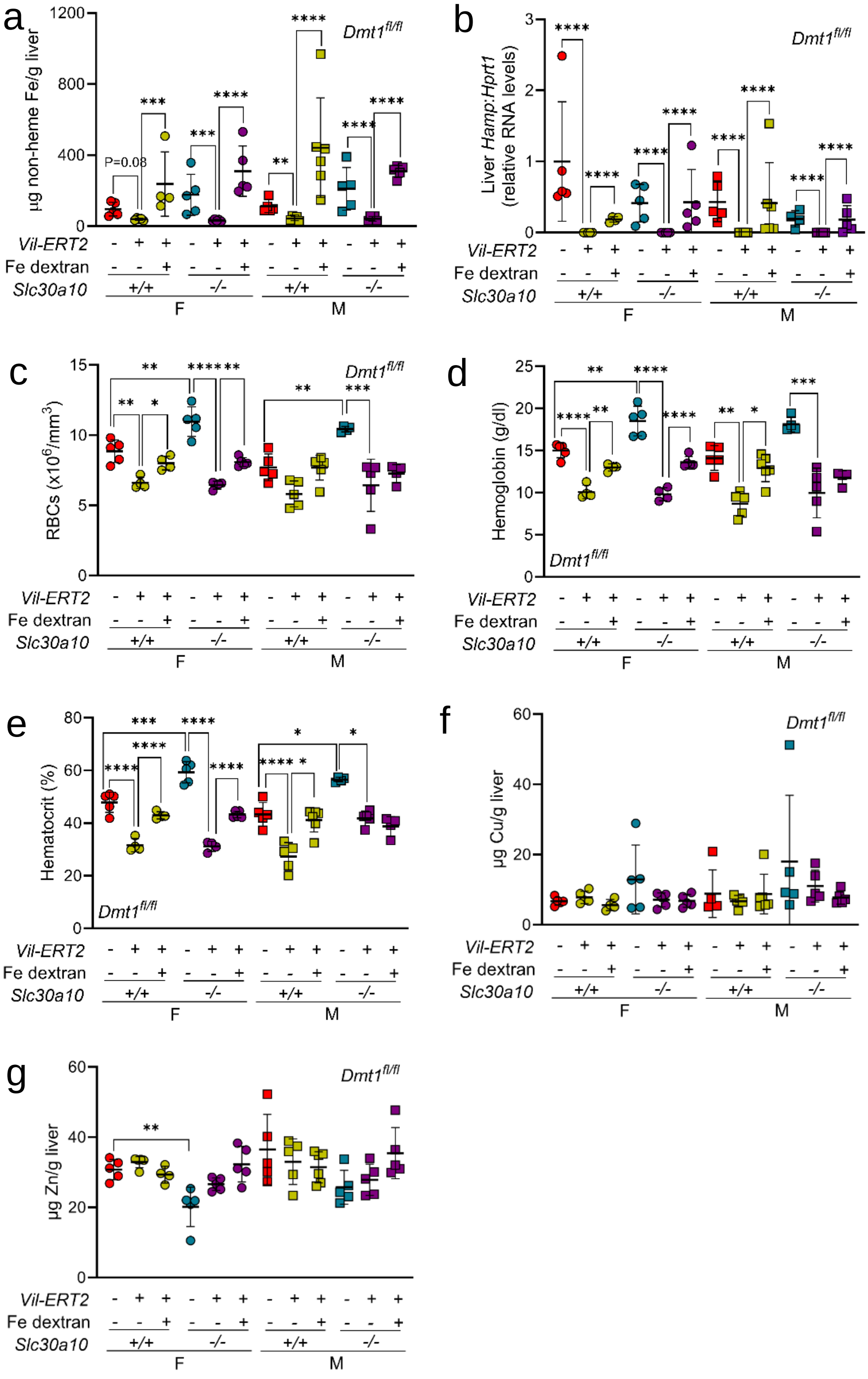
a
Dmt1fl/fl
µg non-heme Fe/g liver
P=0.08
b
Liver Hamp:Hprt1 (relative RNA levels)
c
RBCs (x10^6/mm^3)
d
Hemoglobin (g/dl)
e
Hematocrit (%)
f
µg Cu/g liver
g
µg Zn/g liver
Vil-ERT2
Fe dextran
Slc30a10
+/+
-/-
F
M

Figure S15

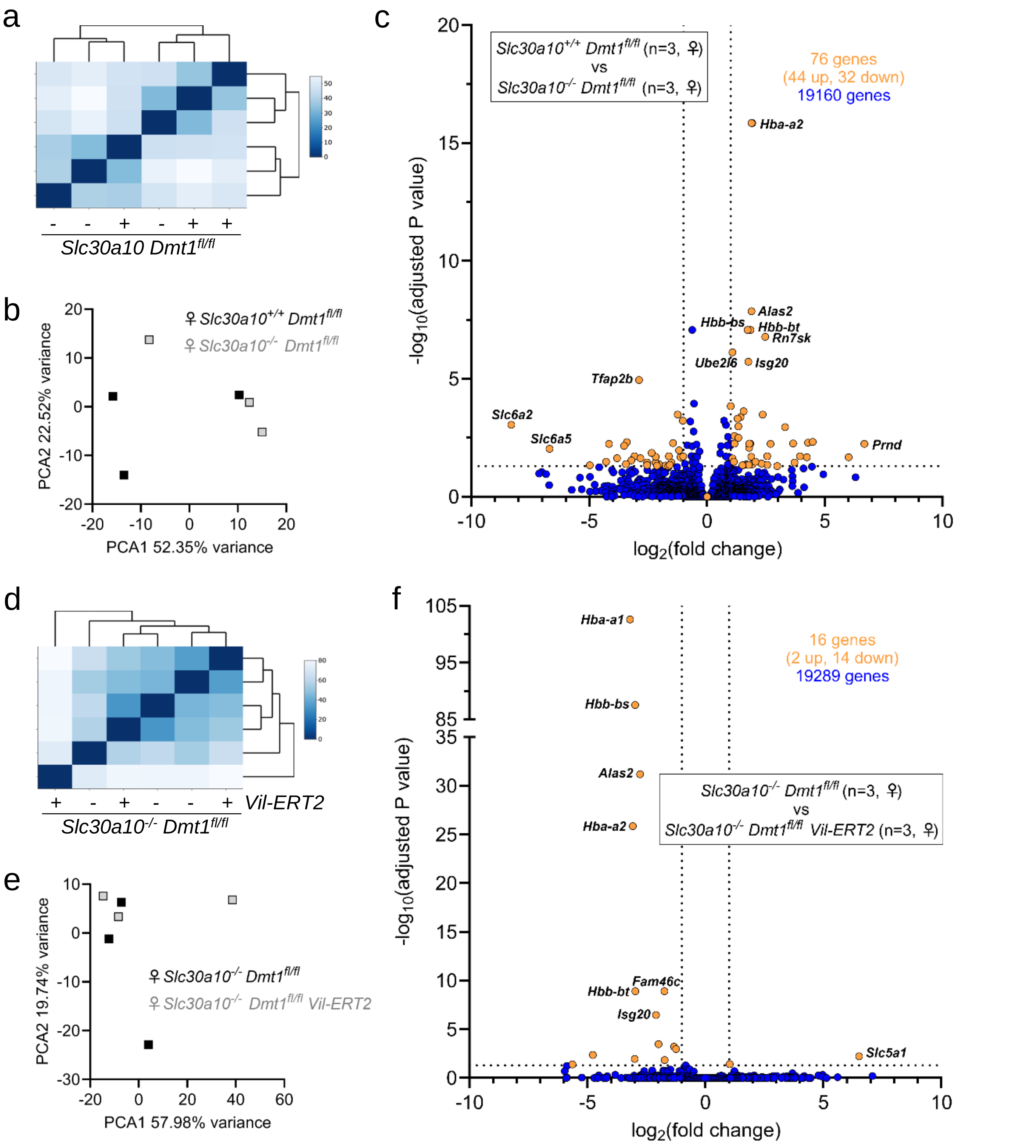
a
b
c
d
e
f
Slc30a10 Dmt1fl/fl
♀ Slc30a10+/+ Dmt1fl/fl
♀ Slc30a10-/- Dmt1fl/fl
PCA2 22.52% variance
PCA1 52.35% variance
Slc30a10+/+ Dmt1fl/fl (n=3, ♀)
vs
Slc30a10-/- Dmt1fl/fl (n=3, ♀)
76 genes
(44 up, 32 down)
19160 genes
Hba-a2
Alas2
Hbb-bs
Hbb-bt
Rn7sk
Ube2l6
Isg20
Tfap2b
Slc6a2
Slc6a5
Prnd
-log10(adjusted P value)
log2(fold change)
Vil-ERT2
Slc30a10-/- Dmt1fl/fl
♀ Slc30a10-/- Dmt1fl/fl
♀ Slc30a10-/- Dmt1fl/fl Vil-ERT2
PCA2 19.74% variance
PCA1 57.98% variance
16 genes
(2 up, 14 down)
19289 genes
Hba-a1
Hbb-bs
Alas2
Hba-a2
Fam46c
Hbb-bt
Isg20
Slc5a1
Slc30a10-/- Dmt1fl/fl (n=3, ♀)
vs
Slc30a10-/- Dmt1fl/fl Vil-ERT2 (n=3, ♀)

Figure S16

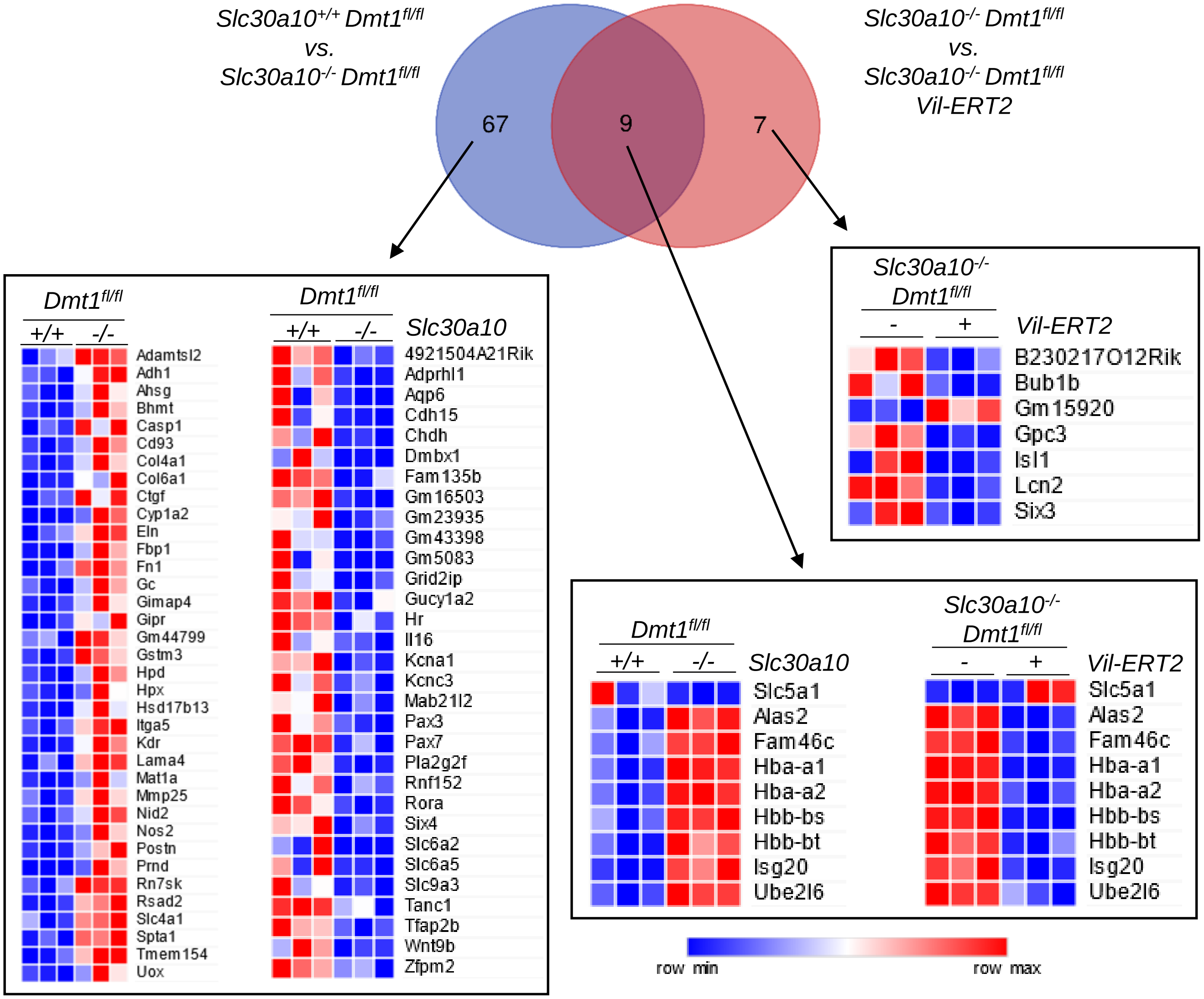

Slc30a10+/+ Dmt1fl/fl vs. Slc30a10-/- Dmt1fl/fl
Slc30a10-/- Dmt1fl/fl vs. Slc30a10-/- Dmt1fl/fl Vil-ERT2
67
9
7
Dmt1fl/fl
+/+
-/-
Slc30a10
Adamtsl2
Adh1
Ahsg
Bhmt
Casp1
Cd93
Col4a1
Col6a1
Ctgf
Cyp1a2
Eln
Fbp1
Fn1
Gc
Gimap4
Gipr
Gm44799
Gstm3
Hpd
Hpx
Hsd17b13
Itga5
Kdr
Lama4
Mat1a
Mmp25
Nid2
Nos2
Postn
Prnd
Rn7sk
Rsad2
Slc4a1
Spta1
Tmem154
Uox
4921504A21Rik
Adprhl1
Aqp6
Cdh15
Chdh
Dmbx1
Fam135b
Gm16503
Gm23935
Gm43398
Gm5083
Grid2ip
Gucy1a2
Hr
Il16
Kcna1
Kcnc3
Mab21l2
Pax3
Pax7
Pla2g2f
Rnf152
Rora
Six4
Slc6a2
Slc6a5
Slc9a3
Tanc1
Tfap2b
Wnt9b
Zfpm2
Slc30a10-/- Dmt1fl/fl
-
+
Vil-ERT2
B230217O12Rik
Bub1b
Gm15920
Gpc3
Isl1
Lcn2
Six3
Slc5a1
Alas2
Fam46c
Hba-a1
Hba-a2
Hbb-bs
Hbb-bt
Isg20
Ube2l6
row min
row max


### Figure S17

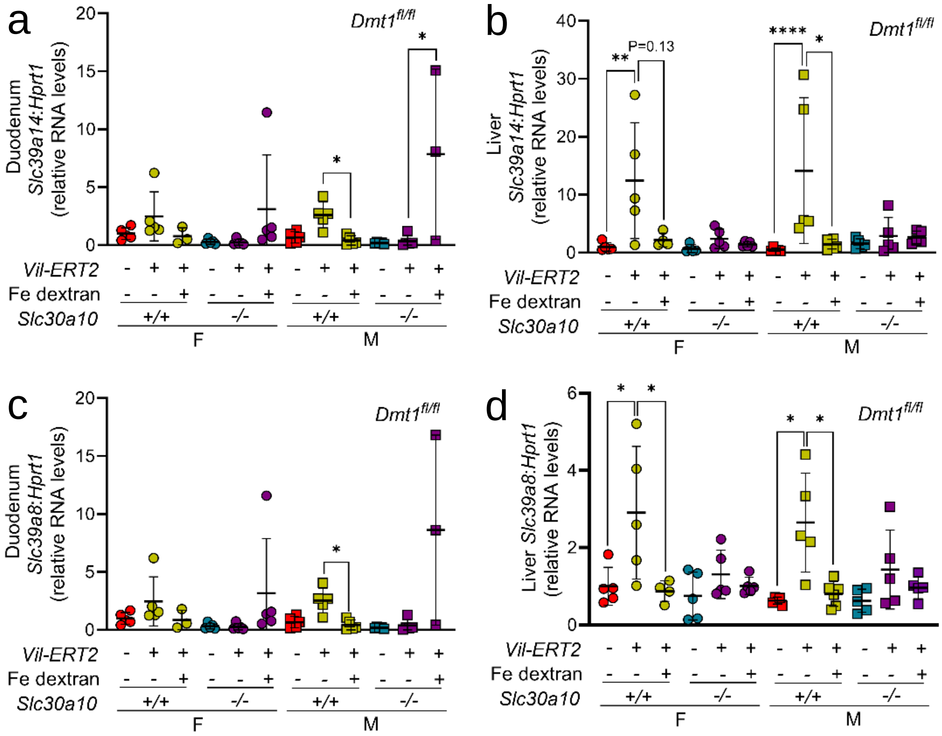
a
Duodenum Slc39a14:Hprt1 (relative RNA levels)
Dmt1fl/fl
P=0.13
b
Liver Slc39a14:Hprt1 (relative RNA levels)
c
Duodenum Slc39a8:Hprt1 (relative RNA levels)
d
Liver Slc39a8:Hprt1 (relative RNA levels)
Vil-ERT2
Fe dextran
Slc30a10
+/+
-/-
F
M

Figure S18

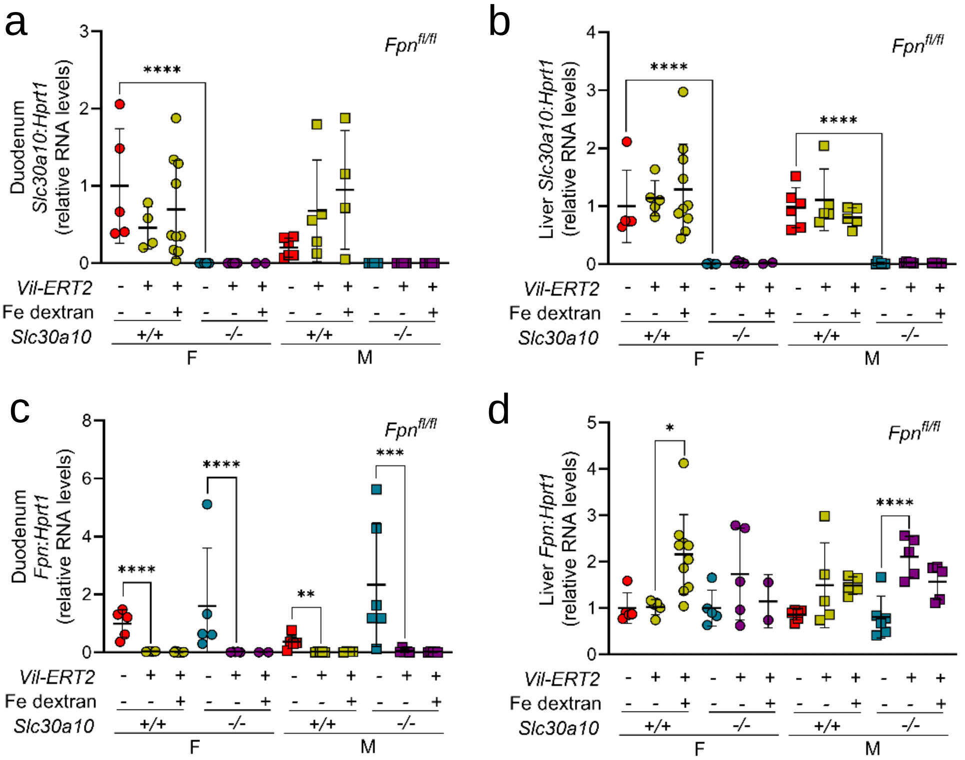
a
Fpn fl/fl
Duodenum Slc30a10:Hprt1 (relative RNA levels)
****
Vil-ERT2
Fe dextran
Slc30a10
+/+
-/-
F
M
b
Fpn fl/fl
Liver Slc30a10:Hprt1 (relative RNA levels)
c
Fpn fl/fl
Duodenum Fpn:Hprt1 (relative RNA levels)
d
Fpn fl/fl
Liver Fpn:Hprt1 (relative RNA levels)

Figure S19

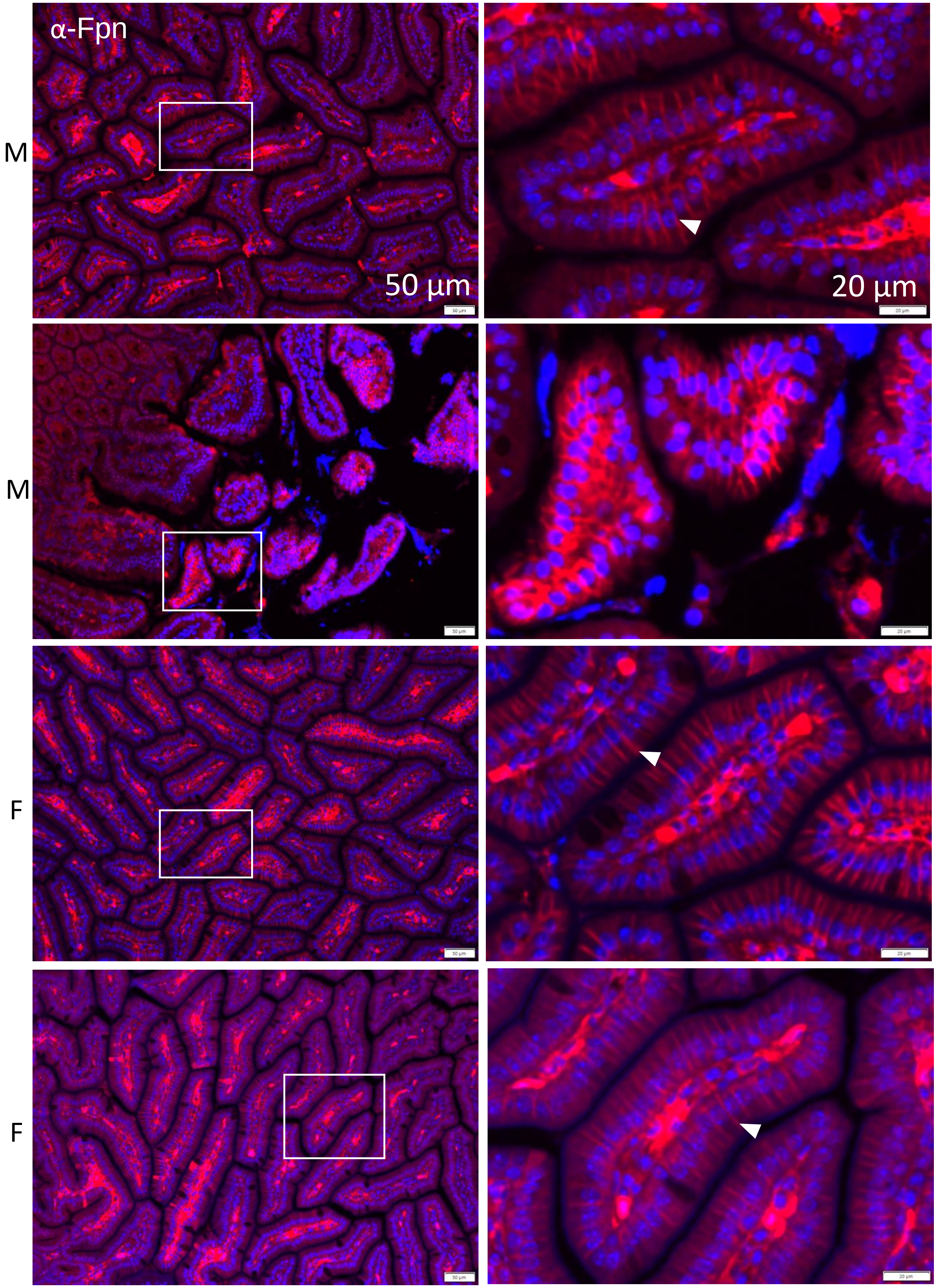


*Slc30a10*$^{+/+}$ *Fpn*$^{fl/fl}$

Figure S20

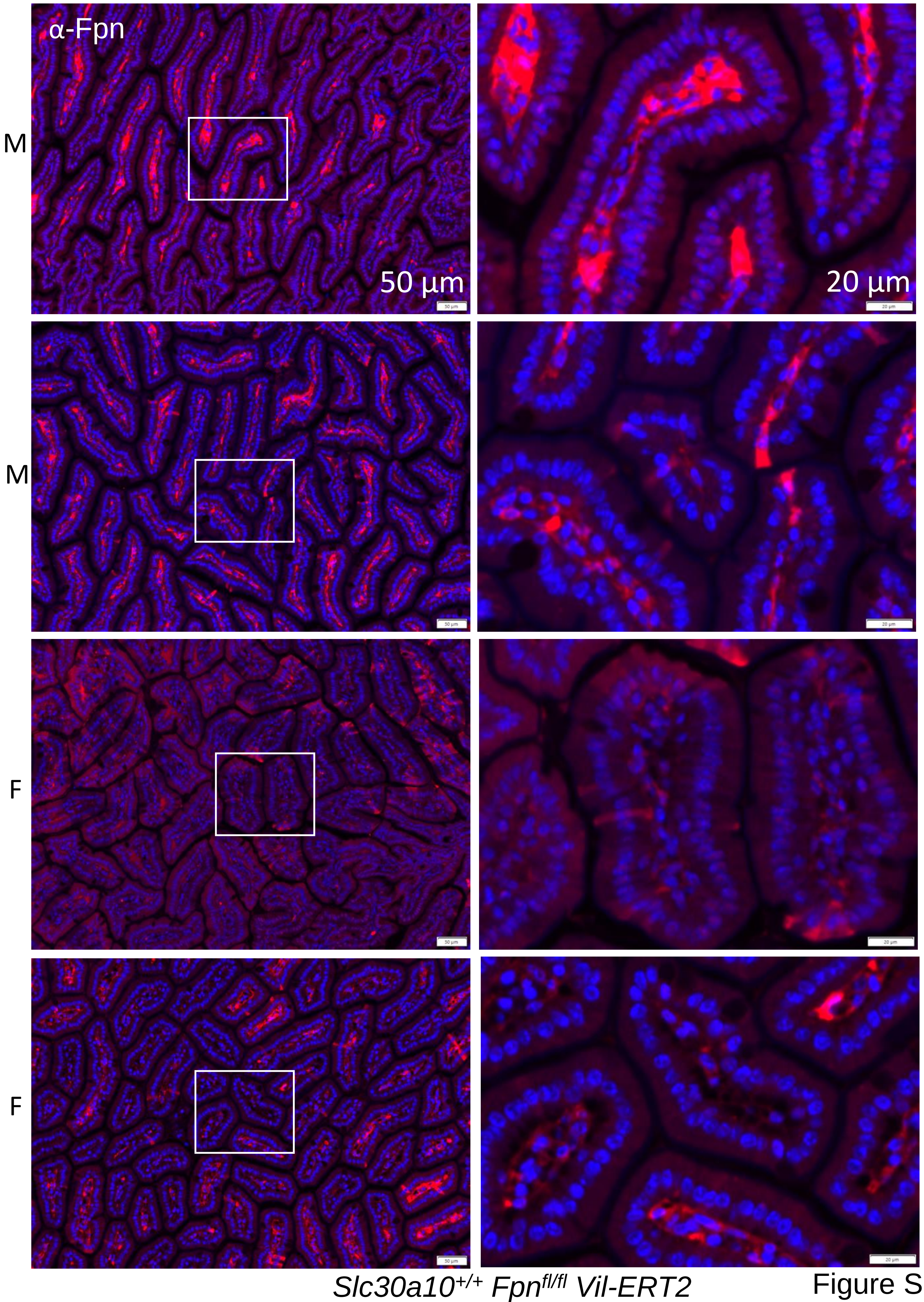


Figure S21

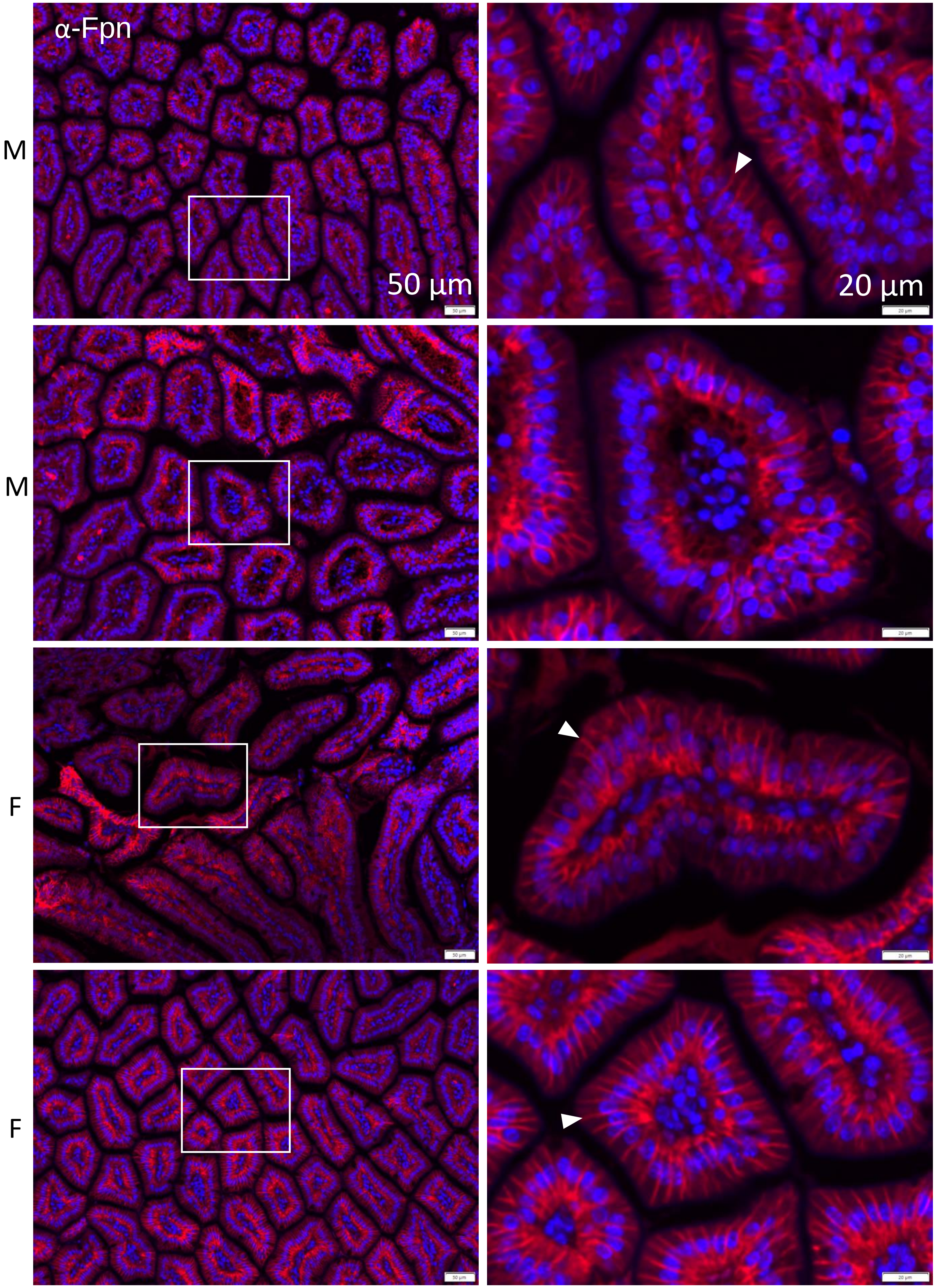


*Slc30a10*$^{-/-}$ *Fpn*$^{fl/fl}$

Figure S22

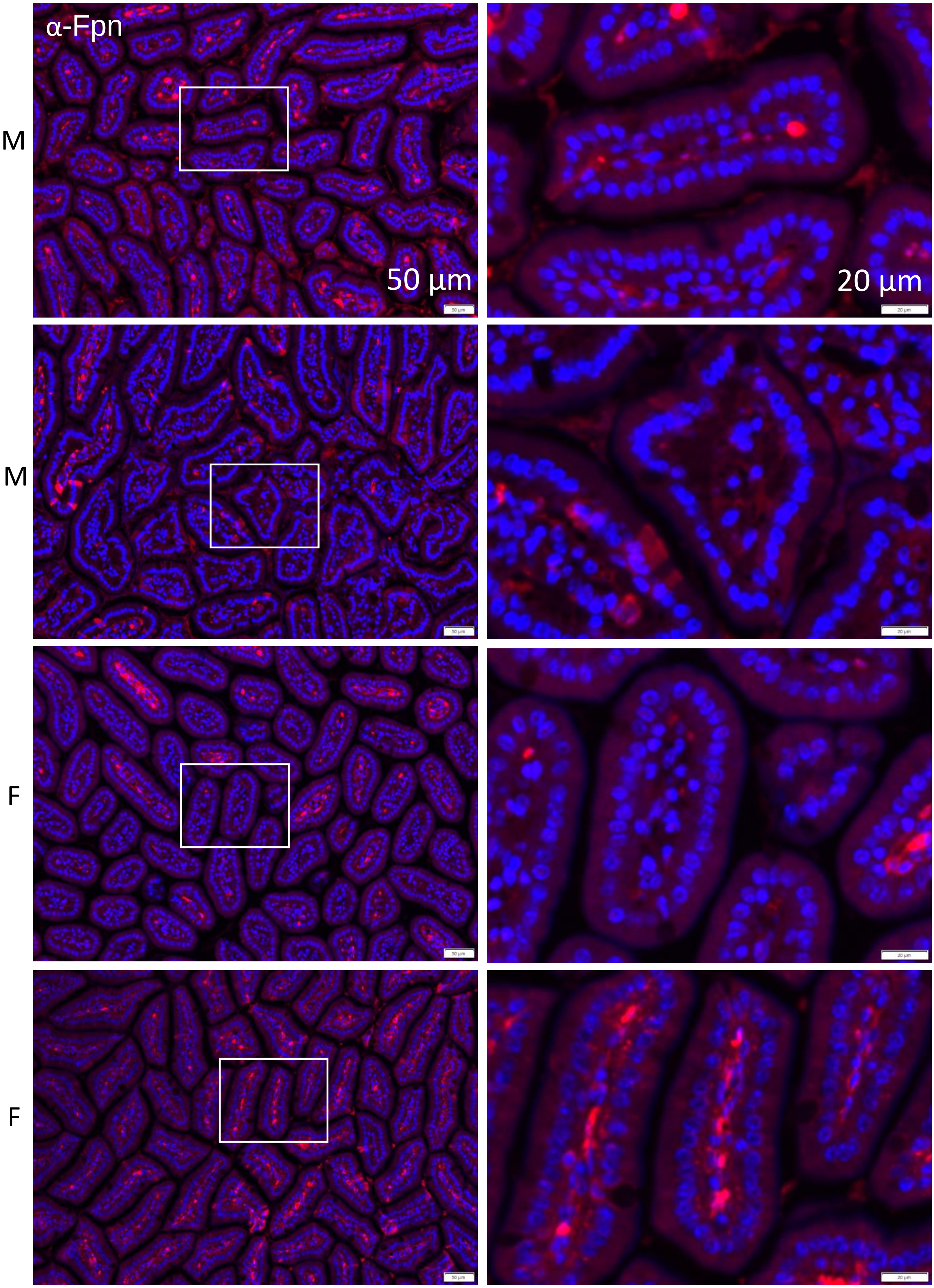


*Slc30a10$^{-/-}$ Fpn$^{fl/fl}$ Vil-ERT2*

Figure S23

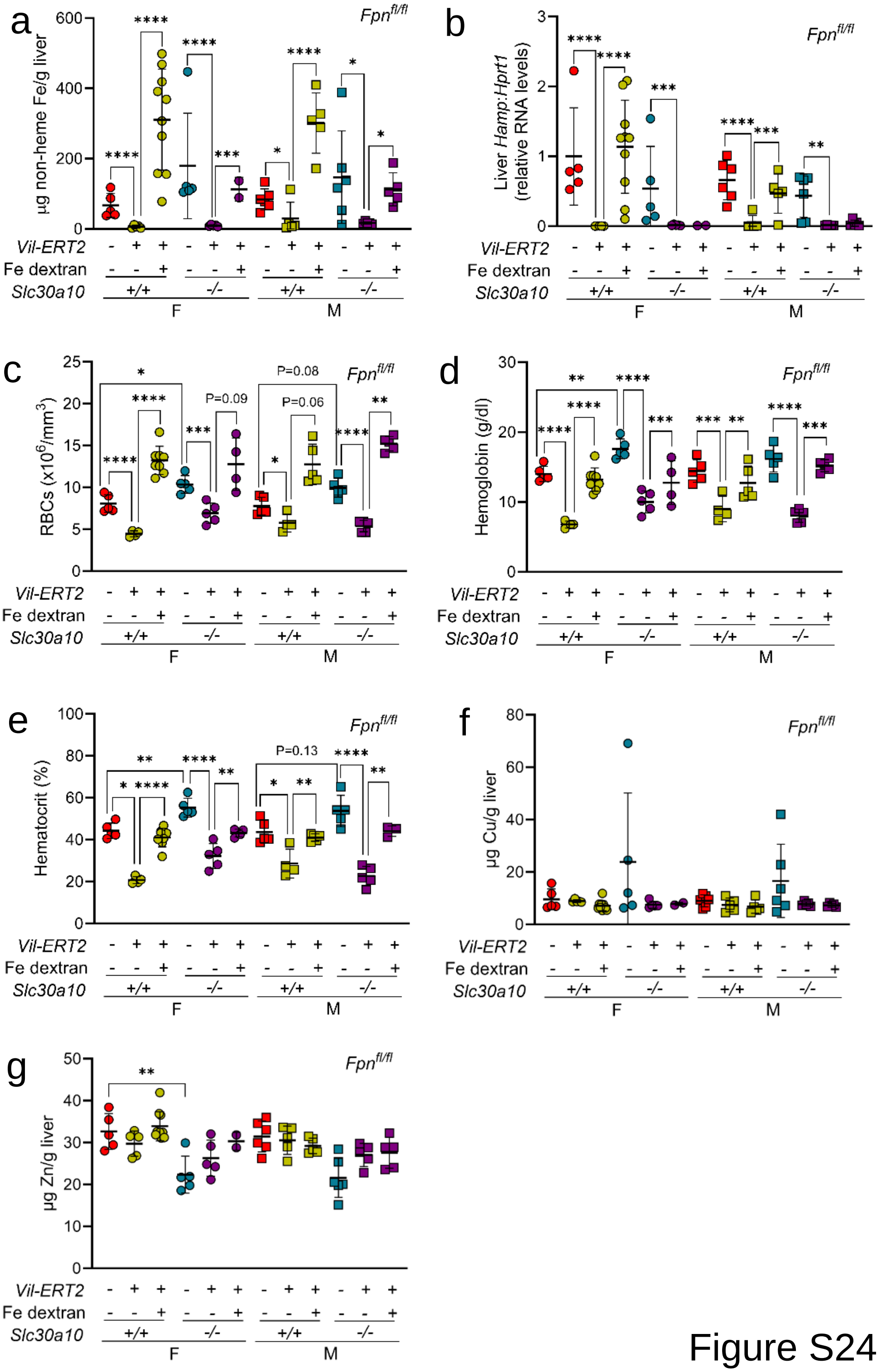

a
b
c
d
e
f
g
Fpnfl/fl
µg non-heme Fe/g liver
Liver Hamp:Hprt1 (relative RNA levels)
RBCs (x10⁶/mm³)
Hemoglobin (g/dl)
Hematocrit (%)
µg Cu/g liver
µg Zn/g liver
P=0.08
P=0.06
P=0.09
P=0.13
Vil-ERT2
Fe dextran
Slc30a10
+/+
-/-
F
M


Figure S24

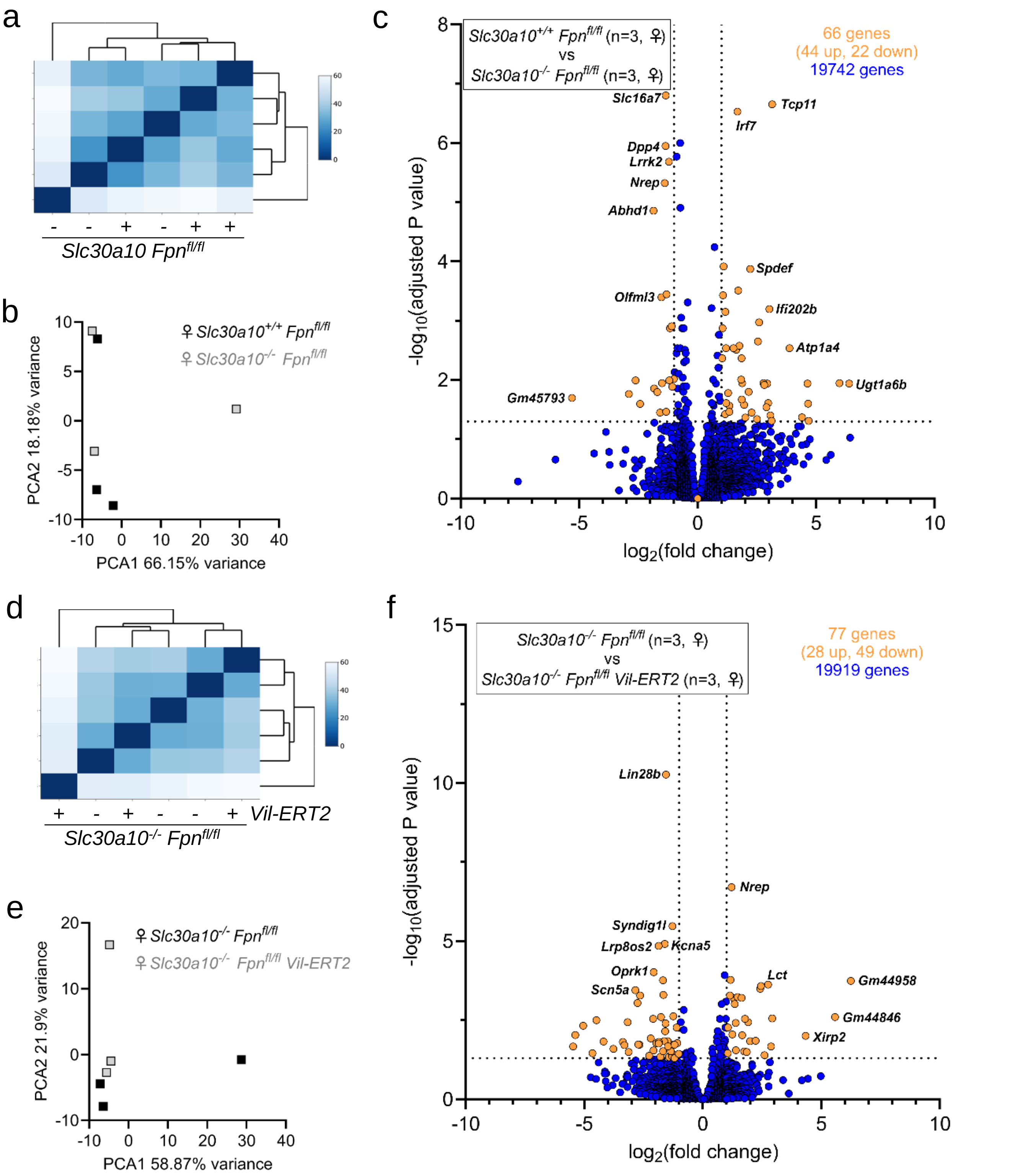

a
- - + - + +
Slc30a10 Fpn^fl/fl
b
♀ Slc30a10^+/+ Fpn^fl/fl
♀ Slc30a10^-/- Fpn^fl/fl
PCA2 18.18% variance
PCA1 66.15% variance
c
Slc30a10^+/+ Fpn^fl/fl (n=3, ♀)
vs
Slc30a10^-/- Fpn^fl/fl (n=3, ♀)
66 genes
(44 up, 22 down)
19742 genes
Slc16a7
Tcp11
Irf7
Dpp4
Lrrk2
Nrep
Abhd1
Spdef
Olfml3
Ifi202b
Atp1a4
Ugt1a6b
Gm45793
-log10(adjusted P value)
log2(fold change)
d
+ - + - - + Vil-ERT2
Slc30a10^-/- Fpn^fl/fl
e
♀ Slc30a10^-/- Fpn^fl/fl
♀ Slc30a10^-/- Fpn^fl/fl Vil-ERT2
PCA2 21.9% variance
PCA1 58.87% variance
f
Slc30a10^-/- Fpn^fl/fl (n=3, ♀)
vs
Slc30a10^-/- Fpn^fl/fl Vil-ERT2 (n=3, ♀)
77 genes
(28 up, 49 down)
19919 genes
Lin28b
Nrep
Syndig1l
Lrp8os2
Kcna5
Oprk1
Lct
Gm44958
Scn5a
Gm44846
Xirp2
-log10(adjusted P value)
log2(fold change)


Figure S25

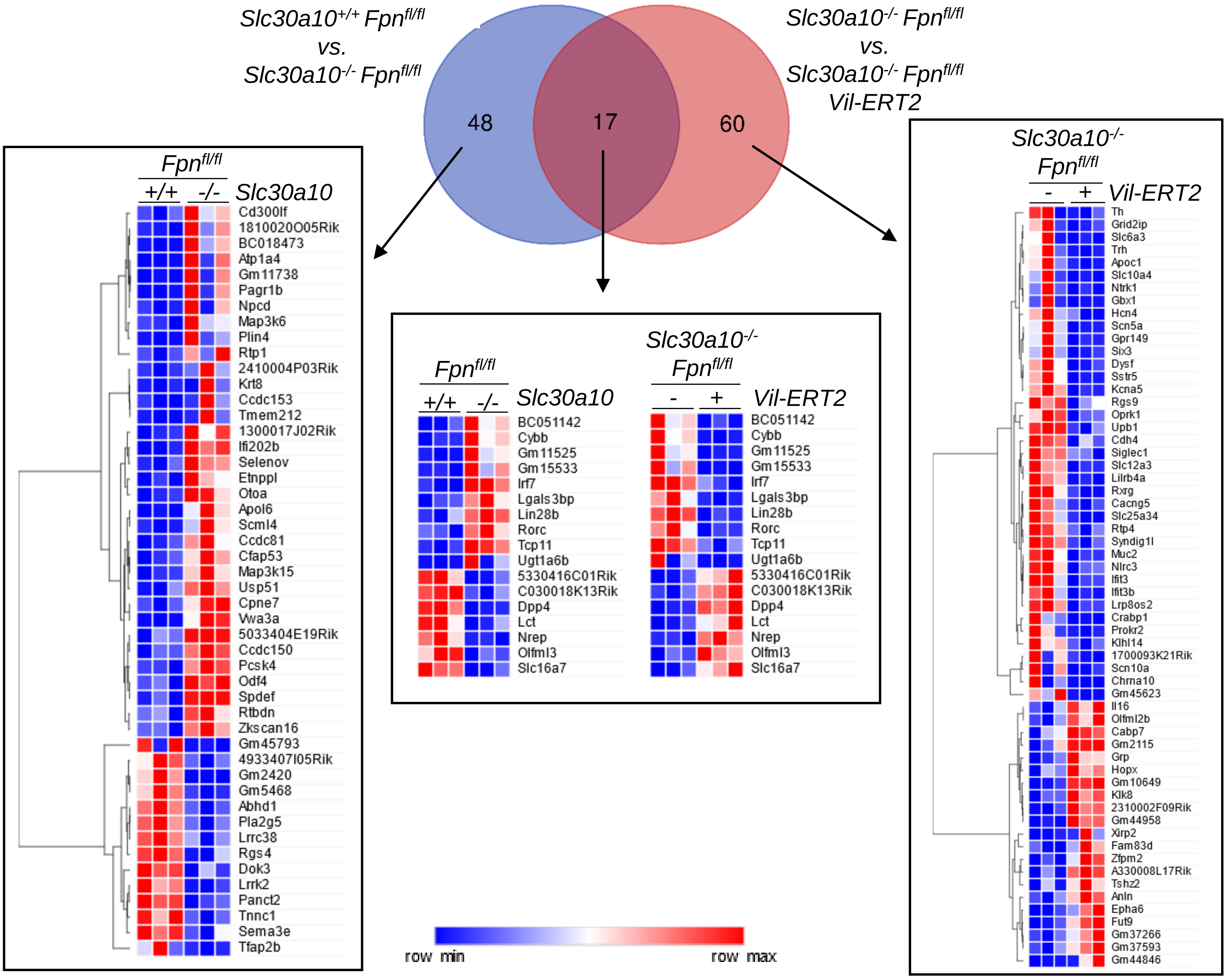

Slc30a10+/+ Fpnfl/fl vs. Slc30a10-/- Fpnfl/fl
Slc30a10-/- Fpnfl/fl vs. Slc30a10-/- Fpnfl/fl Vil-ERT2
48
17
60
Fpnfl/fl
+/+ -/- Slc30a10
Cd300lf
1810020O05Rik
BC018473
Atp1a4
Gm11738
Pagr1b
Npcd
Map3k6
Plin4
Rtp1
2410004P03Rik
Krt8
Ccdc153
Tmem212
1300017J02Rik
Ifi202b
Selenov
Etnppl
Otoa
Apol6
Scml4
Ccdc81
Cfap53
Map3k15
Usp51
Cpne7
Vwa3a
5033404E19Rik
Ccdc150
Pcsk4
Odf4
Spdef
Rtbdn
Zkscan16
Gm45793
4933407I05Rik
Gm2420
Gm5468
Abhd1
Pla2g5
Lrrc38
Rgs4
Dok3
Lrrk2
Panct2
Tnnc1
Sema3e
Tfap2b
Fpnfl/fl
+/+ -/- Slc30a10
Slc30a10-/- Fpnfl/fl
- + Vil-ERT2
BC051142
Cybb
Gm11525
Gm15533
Irf7
Lgals3bp
Lin28b
Rorc
Tcp11
Ugt1a6b
5330416C01Rik
C030018K13Rik
Dpp4
Lct
Nrep
Olfml3
Slc16a7
row min
row max
Slc30a10-/- Fpnfl/fl
- + Vil-ERT2
Th
Grid2ip
Slc6a3
Trh
Apoc1
Slc10a4
Ntrk1
Gbx1
Hcn4
Scn5a
Gpr149
Six3
Dysf
Sstr5
Kcna5
Rgs9
Oprk1
Upb1
Cdh4
Siglec1
Slc12a3
Lilrb4a
Rxrg
Cacng5
Slc25a34
Rtp4
Syndig1l
Muc2
Nlrc3
Ifit3
Ifit3b
Lrp8os2
Crabp1
Prokr2
Klhl14
1700093K21Rik
Scn10a
Chrna10
Gm45623
Il16
Olfml2b
Cabp7
Gm2115
Grp
Hopx
Gm10649
Klk8
2310002F09Rik
Gm44958
Xirp2
Fam83d
Zfpm2
A330008L17Rik
Tshz2
Anln
Epha6
Fut9
Gm37266
Gm37593
Gm44846


### Figure S26

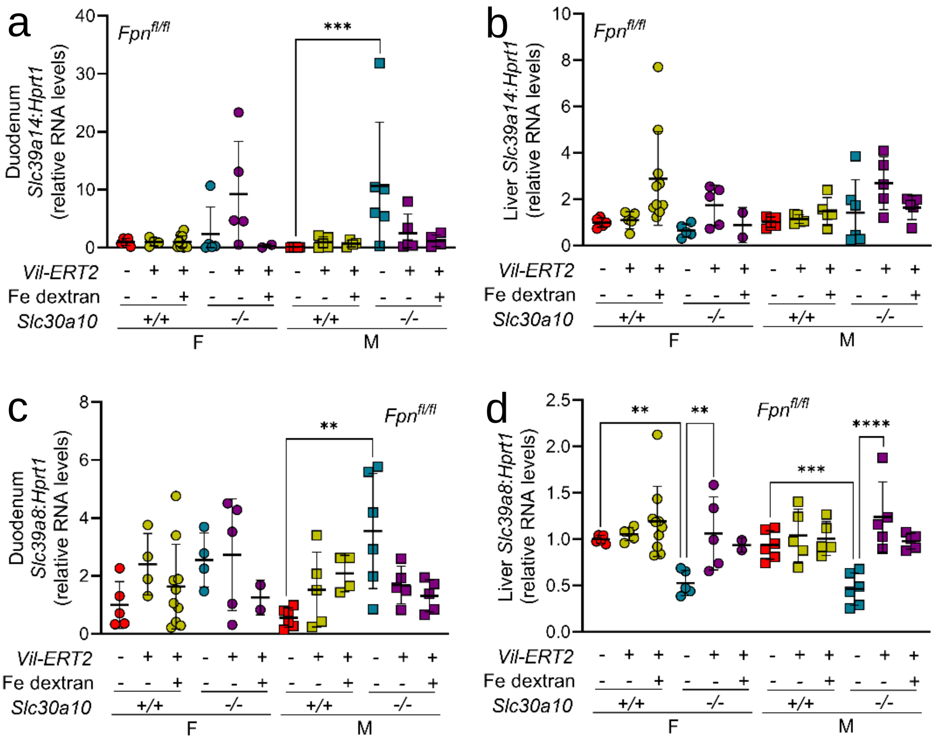
a
Duodenum Slc39a14:Hprt1 (relative RNA levels)
b
Liver Slc39a14:Hprt1 (relative RNA levels)
c
Duodenum Slc39a8:Hprt1 (relative RNA levels)
d
Liver Slc39a8:Hprt1 (relative RNA levels)
Fpn fl/fl
Vil-ERT2
Fe dextran
Slc30a10
+/+
-/-
F
M

Figure S27

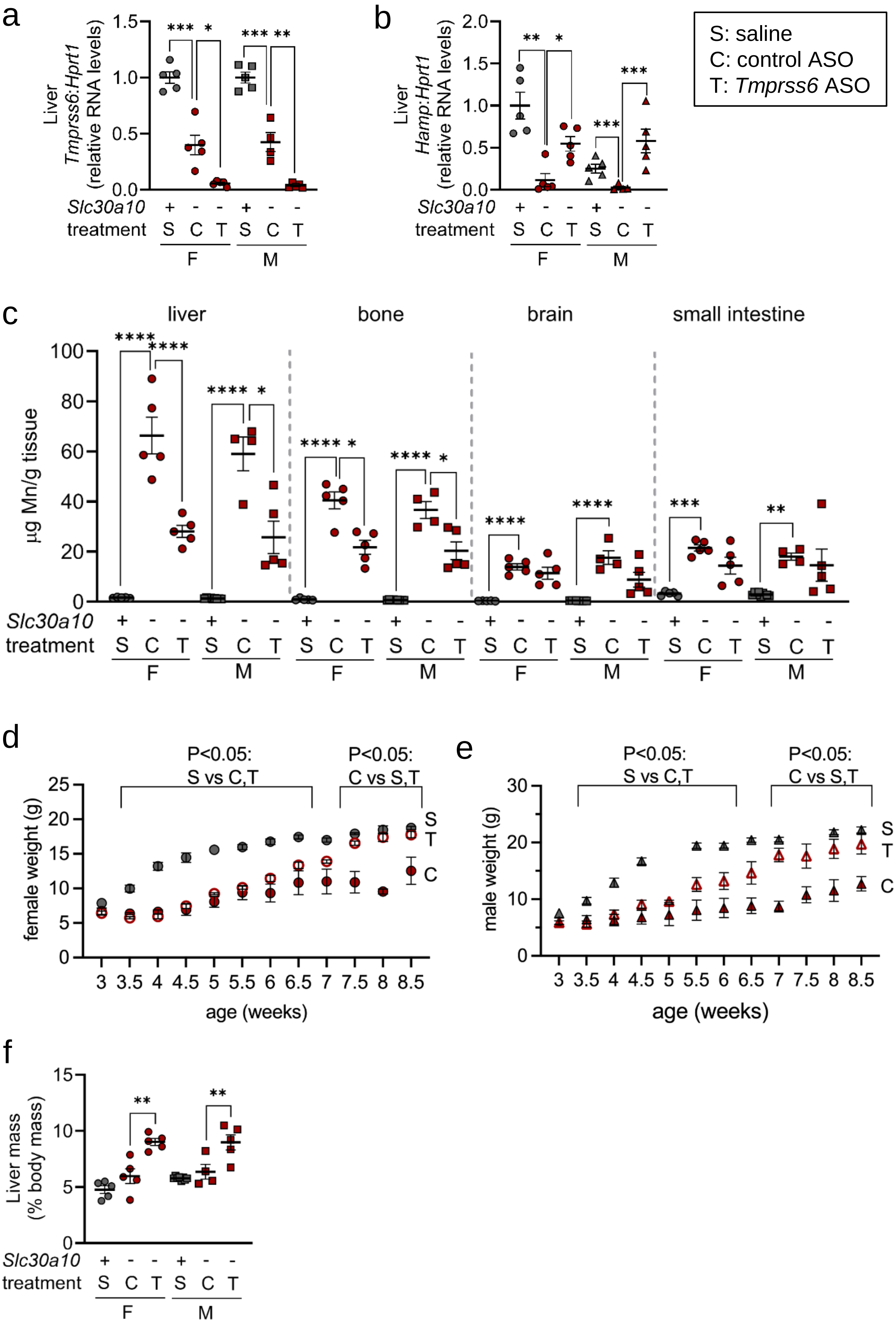

a
Liver
Tmprss6:Hprt1
(relative RNA levels)
Slc30a10
treatment
F
M
b
Liver
Hamp:Hprt1
(relative RNA levels)
S: saline
C: control ASO
T: Tmprss6 ASO
c
liver
bone
brain
small intestine
μg Mn/g tissue
d
P<0.05:
S vs C,T
P<0.05:
C vs S,T
female weight (g)
age (weeks)
e
male weight (g)
f
Liver mass
(% body mass)


Figure S28

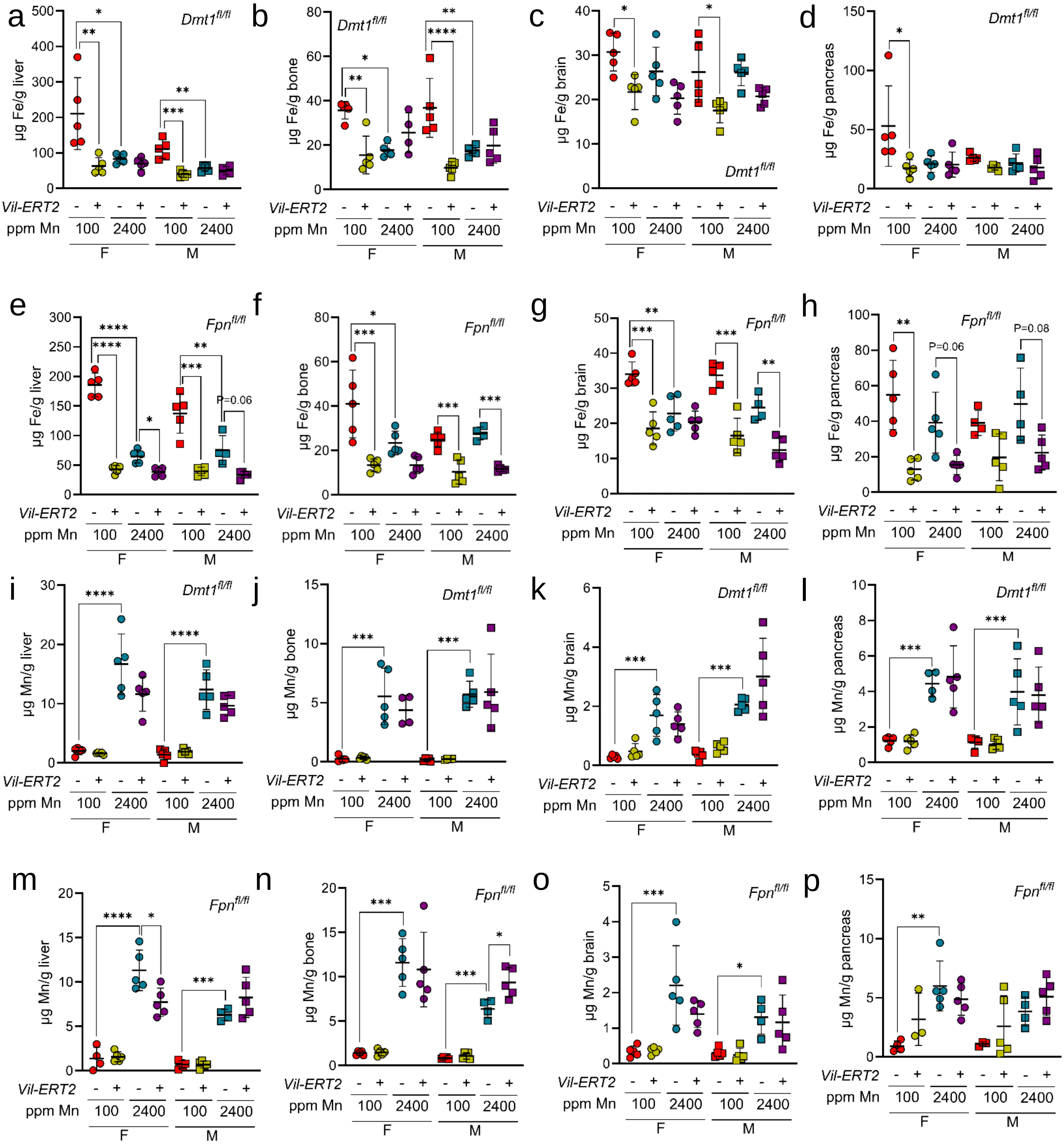
a
b
c
d
e
f
g
h
i
j
k
l
m
n
o
p
Dmt1fl/fl
Fpnfl/fl
µg Fe/g liver
µg Fe/g bone
µg Fe/g brain
µg Fe/g pancreas
µg Mn/g liver
µg Mn/g bone
µg Mn/g brain
µg Mn/g pancreas
Vil-ERT2
ppm Mn
100
2400
F
M
P=0.06
P=0.08

Figure S29

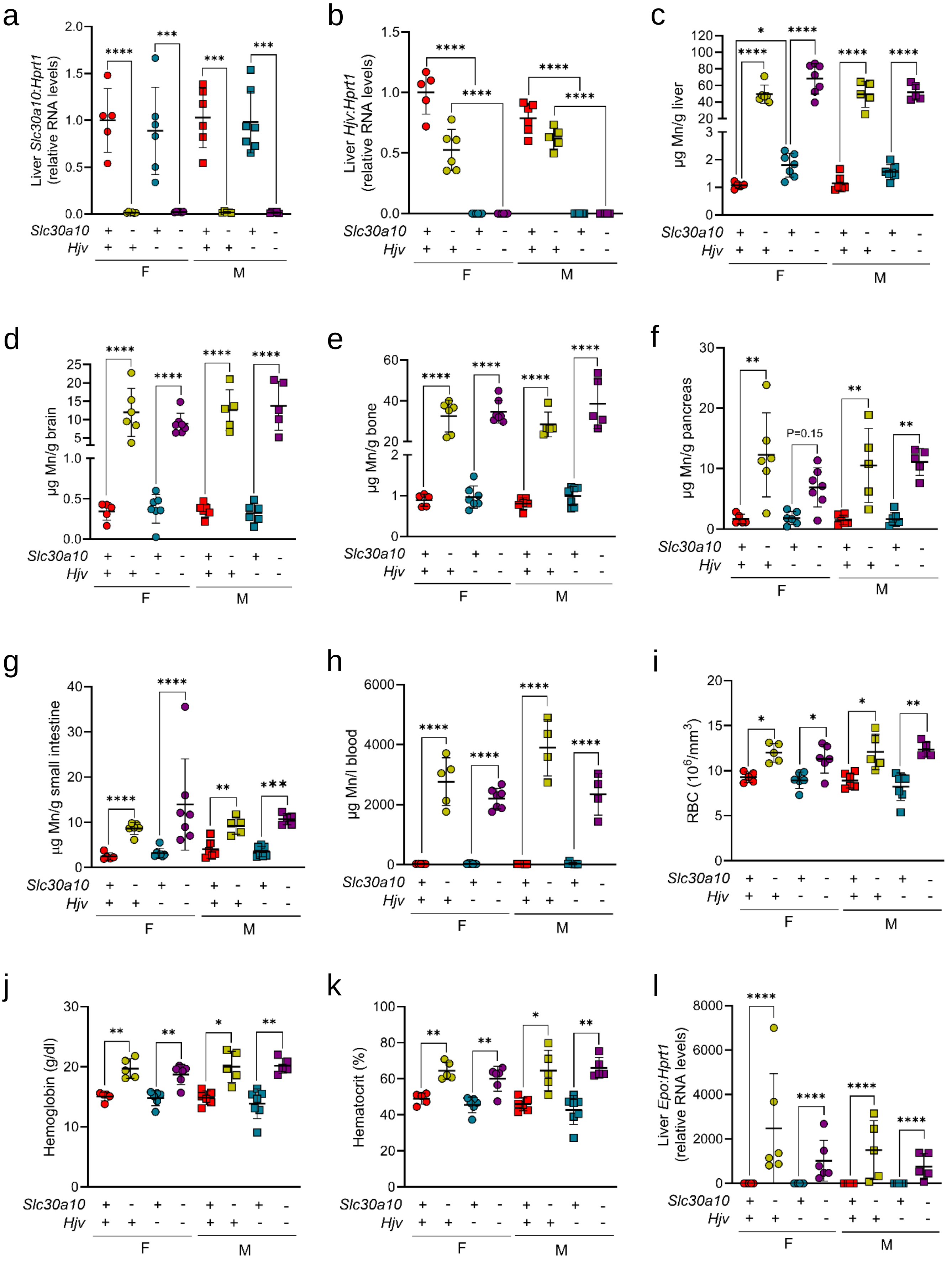
a
Liver Slc30a10:Hprt1 (relative RNA levels)
b
Liver Hjv:Hprt1 (relative RNA levels)
c
μg Mn/g liver
d
μg Mn/g brain
e
μg Mn/g bone
f
μg Mn/g pancreas
P=0.15
g
μg Mn/g small intestine
h
μg Mn/l blood
i
RBC (10^6/mm^3)
j
Hemoglobin (g/dl)
k
Hematocrit (%)
l
Liver Epo:Hprt1 (relative RNA levels)
Slc30a10
Hjv
F
M

Figure S30

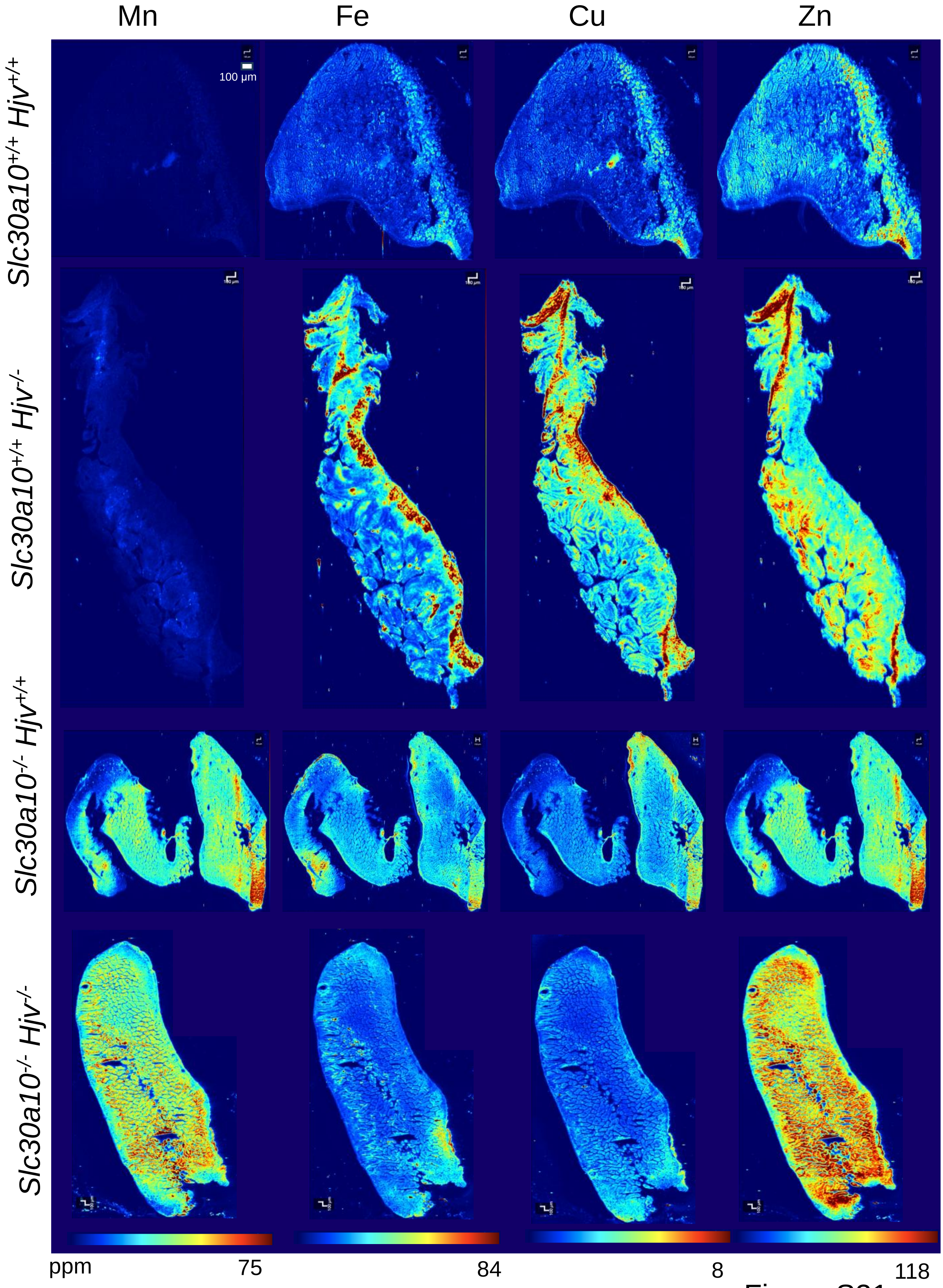

Mn
Fe
Cu
Zn
Slc30a10+/+ Hjv+/+
Slc30a10+/+ Hjv-/-
Slc30a10-/- Hjv+/+
Slc30a10-/- Hjv-/-
100 μm
ppm
75
84
8
118


Figure S31

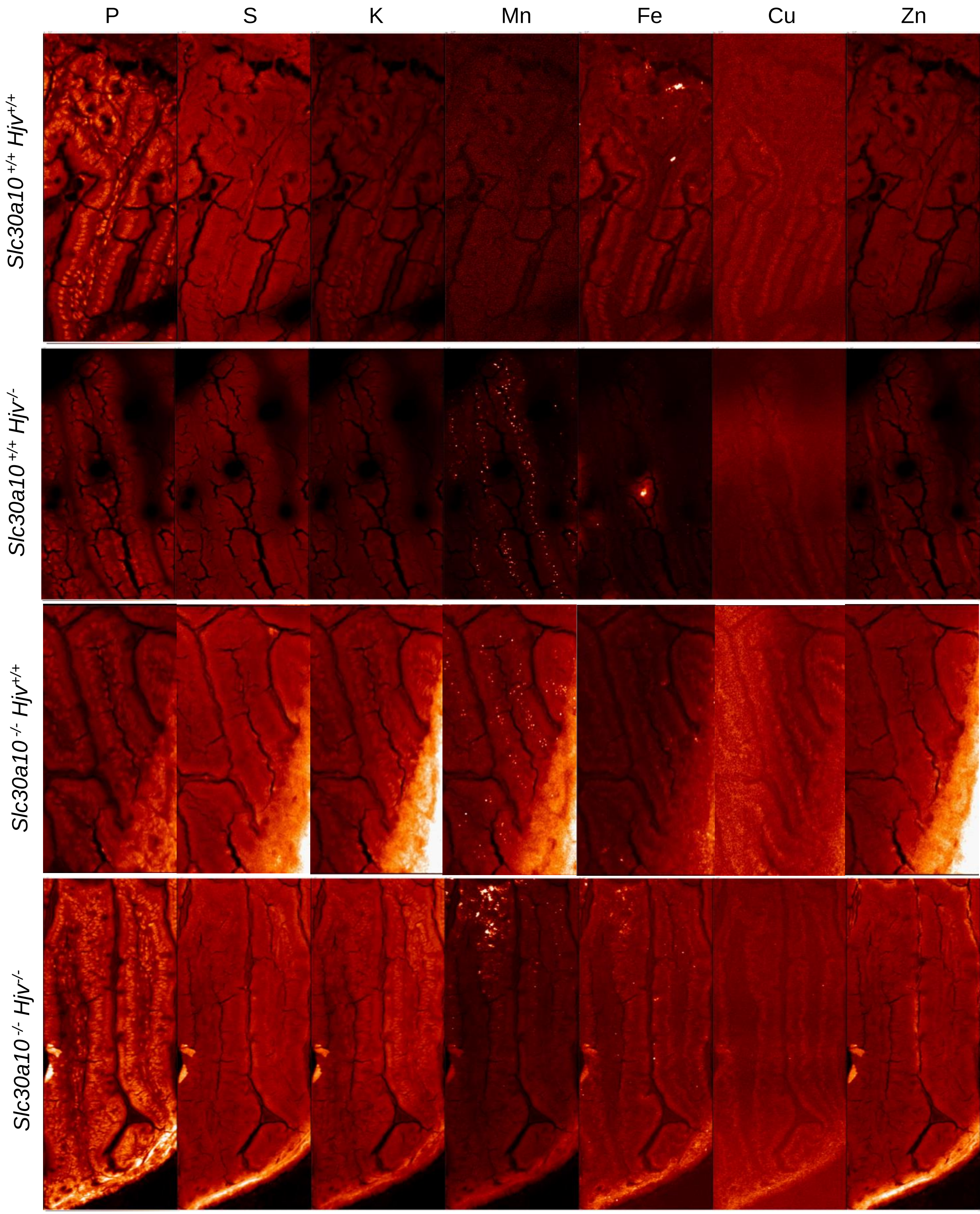
P
S
K
Mn
Fe
Cu
Zn
Slc30a10 +/+ Hjv+/+
Slc30a10 +/+ Hjv-/-
Slc30a10 -/- Hjv+/+
Slc30a10 -/- Hjv-/-

Figure S32

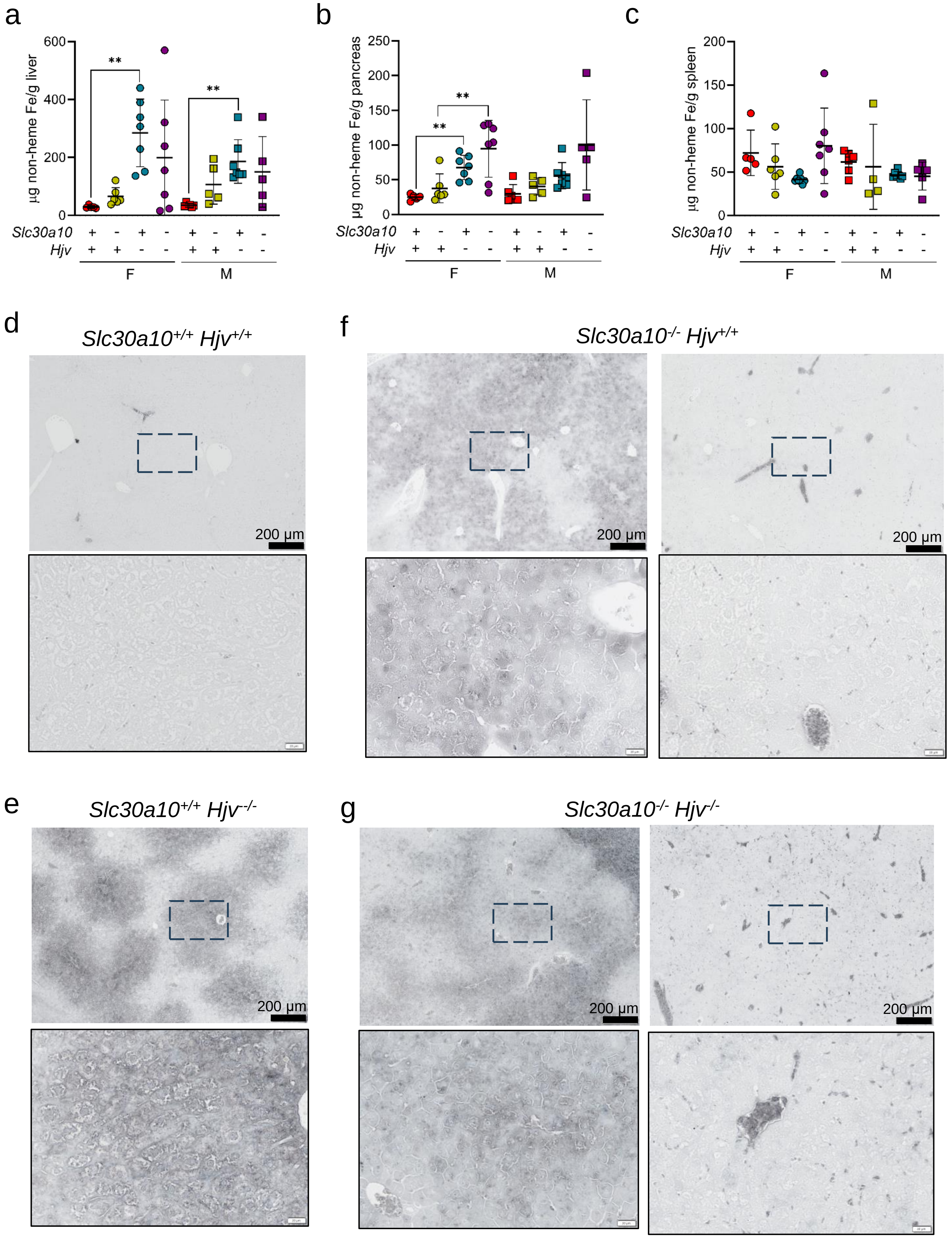

a
μg non-heme Fe/g liver
600
400
200
0
**
**
Slc30a10 + - + - + - + -
Hjv + + - - + + - -
F
M
b
μg non-heme Fe/g pancreas
250
200
150
100
50
0
**
**
Slc30a10 + - + - + - + -
Hjv + + - - + + - -
F
M
c
μg non-heme Fe/g spleen
200
150
100
50
0
Slc30a10 + - + - + - + -
Hjv + + - - + + - -
F
M
d
Slc30a10+/+ Hjv+/+
200 μm
f
Slc30a10-/- Hjv+/+
200 μm
200 μm
e
Slc30a10+/+ Hjv-/-
200 μm
g
Slc30a10-/- Hjv-/-
200 μm
200 μm


Figure S33

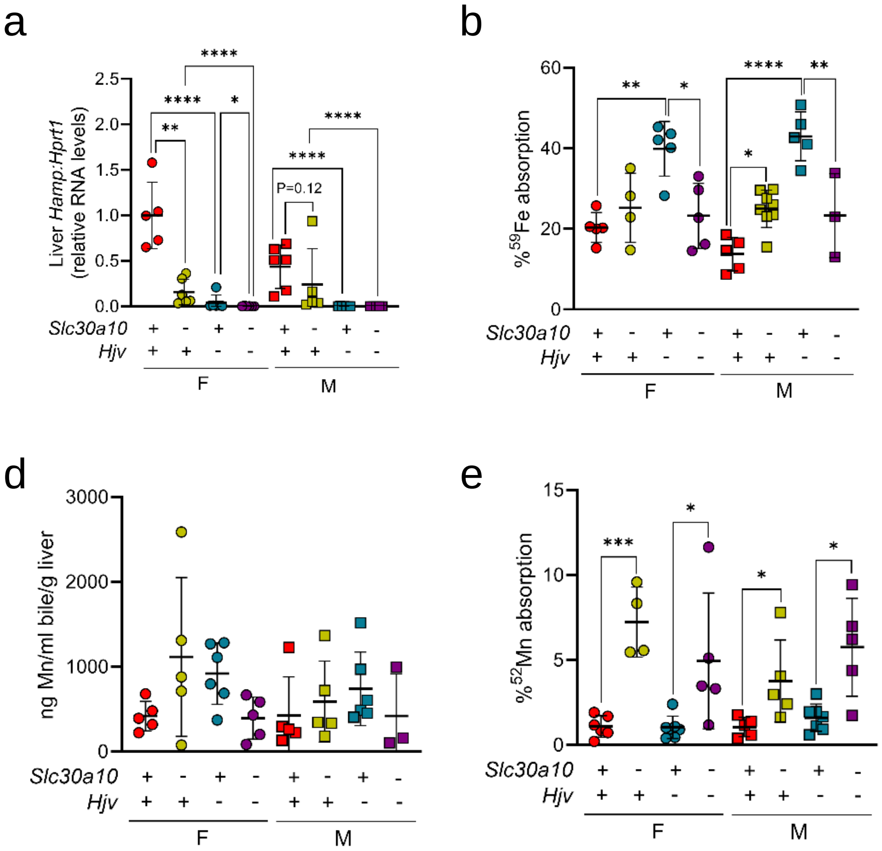
a
Liver Hamp:Hprt1 (relative RNA levels)
P=0.12
b
% 59Fe absorption
c
µl bile/g liver/minute
d
ng Mn/ml bile/g liver
e
% 52Mn absorption
Slc30a10
Hjv
F
M

Figure S34

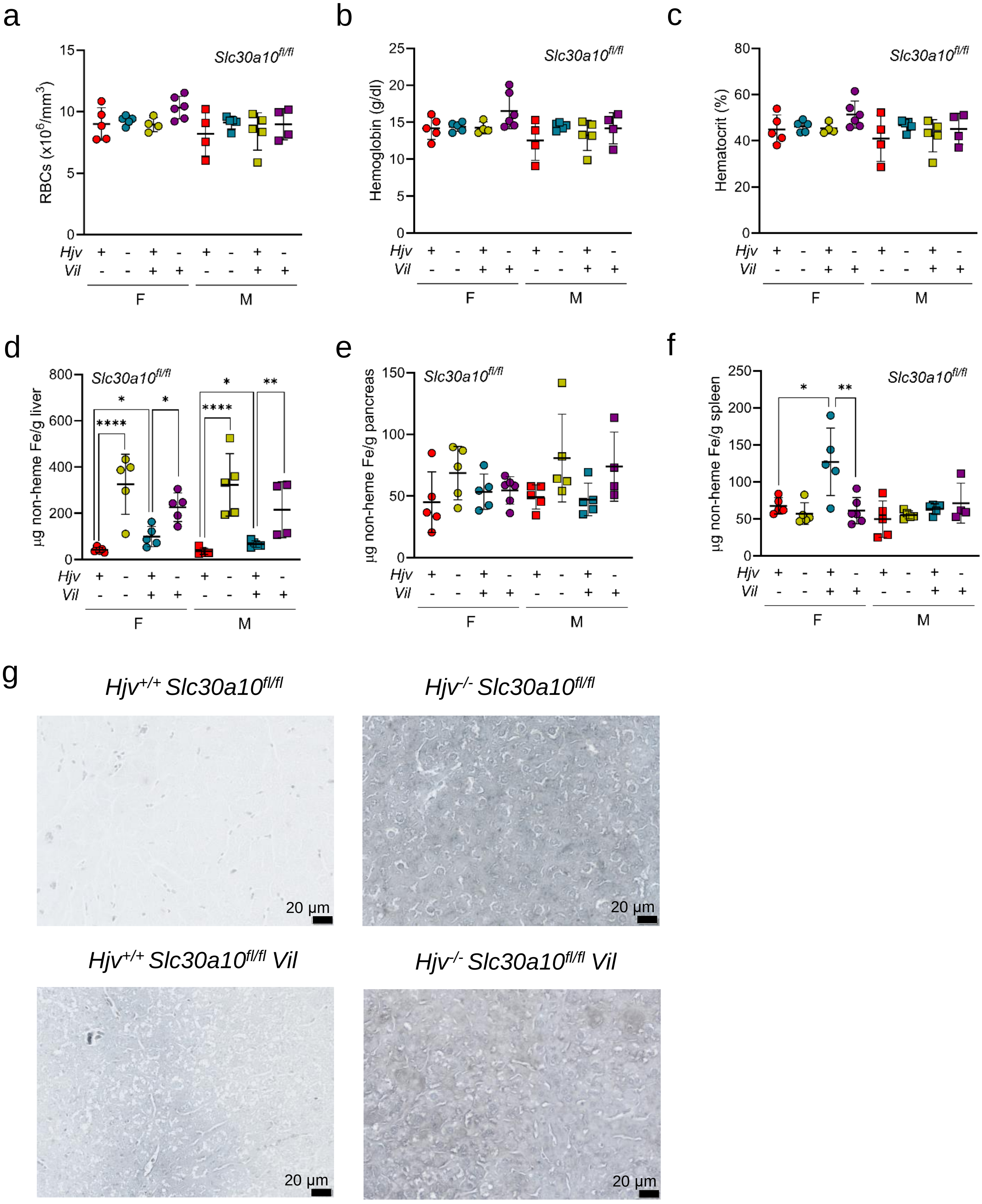
a
b
c
Slc30a10fl/fl
RBCs (x10^6/mm^3)
Hemoglobin (g/dl)
Hematocrit (%)
Hjv
Vil
F
M
d
e
f
µg non-heme Fe/g liver
µg non-heme Fe/g pancreas
µg non-heme Fe/g spleen
g
Hjv+/+ Slc30a10fl/fl
Hjv-/- Slc30a10fl/fl
Hjv+/+ Slc30a10fl/fl Vil
Hjv-/- Slc30a10fl/fl Vil
20 µm

Figure S35
